# Molecular glues of the regulatory ChREBP/14-3-3 complex protect beta cells from glucolipotoxicity

**DOI:** 10.1101/2024.02.16.580675

**Authors:** Liora S. Katz, Emira J. Visser, Kathrin F. Plitzko, Marloes A.M. Pennings, Peter J. Cossar, Isabelle L. Tse, Markus Kaiser, Luc Brunsveld, Christian Ottmann, Donald K. Scott

## Abstract

The Carbohydrate Response Element Binding Protein (ChREBP) is a glucose-responsive transcription factor (TF) with two major splice isoforms (α and β). In chronic hyperglycemia and glucolipotoxicity, ChREBPα-mediated ChREBPβ expression surges, leading to insulin-secreting β-cell dedifferentiation and death. 14-3-3 binding to ChREBPα results in cytoplasmic retention and suppression of transcriptional activity. Thus, small molecule-mediated stabilization of this protein-protein interaction (PPI) may be of therapeutic value. Here, we show that structure-based optimizations of a ‘molecular glue’ compound led to potent ChREBPα/14-3-3 PPI stabilizers with cellular activity. In primary human β-cells, the most active compound retained ChREBPα in the cytoplasm, and efficiently protected β-cells from glucolipotoxicity while maintaining β-cell identity. This study may thus not only provide the basis for the development of a unique class of compounds for the treatment of Type 2 Diabetes but also showcases an alternative ‘molecular glue’ approach for achieving small molecule control of notoriously difficult to target TFs.

**Graphical Abstract:** 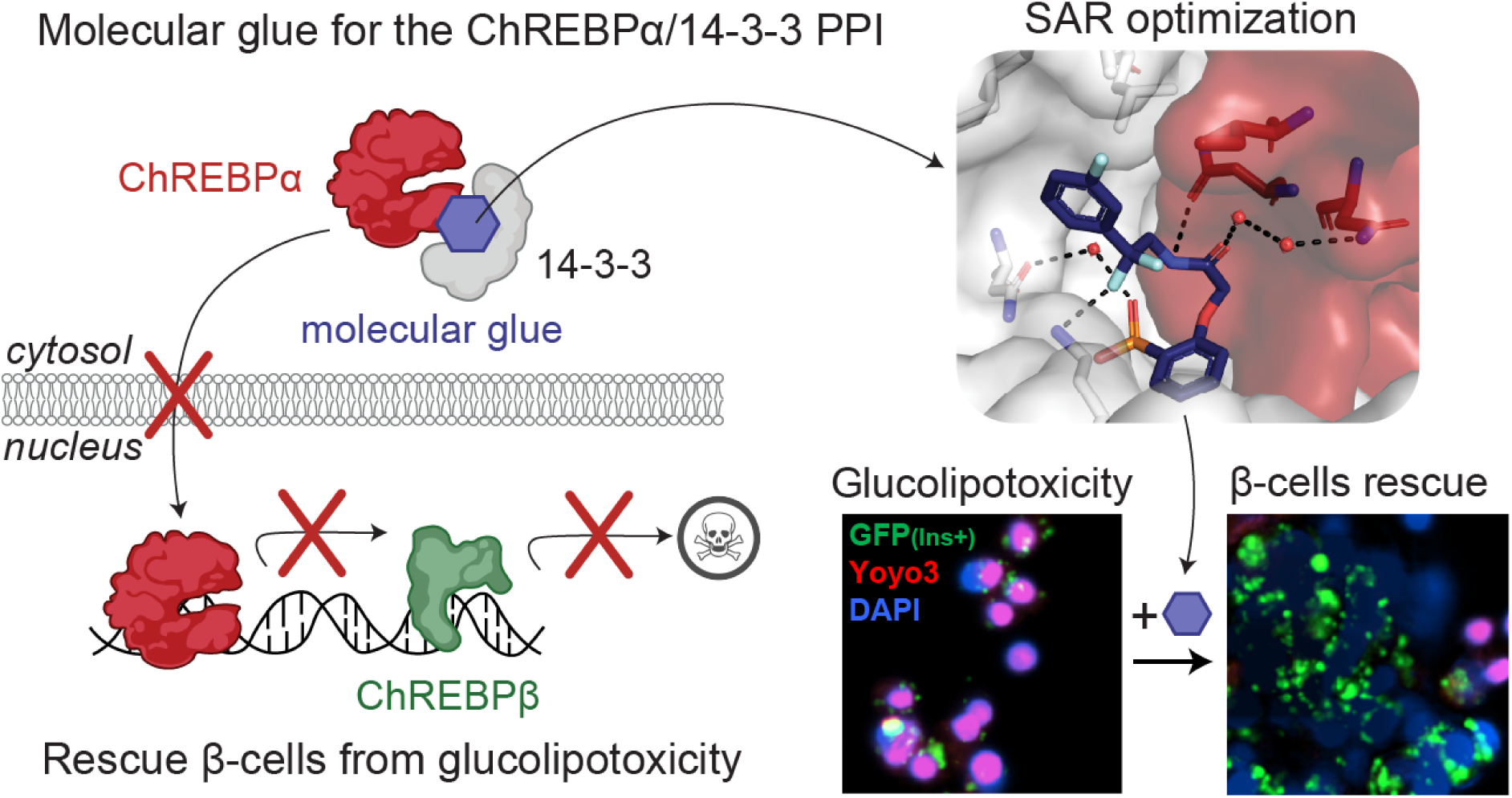

## Introduction

Type 2 diabetes (T2D) imposes an escalating burden on global health and economy. In 2021 537 million adults worldwide had diabetes, and this is expected to rise to 693 million by 2045^1^ or even 1.31 billion by 2050^2^, making it one of the leading causes of global morbidity^3^. Both major types of diabetes are characterized by insufficient functional β-cell mass to meet the increased demand for insulin. T2D develops due to decreased insulin response^4^, triggering a vicious cycle of increased insulin demand and decreased peripheral tissue response. While lifestyle adjustments can alleviate insulin resistance^5^, inadequate hyperglycemic control can lead to β-cell exhaustion, dedifferentiation, and eventual demise due to metabolic stress. This final stage is irreversible due to the restricted proliferative capacity of adult β-cells. Current T2D therapies mainly target insulin resistance and secretion, yet many T2D patients eventually become insulin dependent due to the loss of functional β-cells. Preventing the loss of functional β-cell mass remains one of the most important unmet needs in the armamentarium for the treatment of diabetes.

An emerging potential target to treat T2D is the glucose-responsive transcription factor (TF) Carbohydrate Response Element Binding Protein (ChREBP)^6, 7^. ChREBP is a key mediator in the response to glucose in pancreatic β-cells, controlling the expression of glycolytic and lipogenic genes^8^. Small molecule modulation of ChREBP function may thus represent a promising approach to combat T2D; however, many TFs such as ChREBP are known as notoriously difficult to target with small molecules due to their lack of suitable ligand binding sites^9^. However, several proteins interact with ChREBP to regulate its activation mechanism, including the 14-3-3 proteins^10, 11^. The protein-protein interaction (PPI) between ChREBP and 14-3-3 potentially offer long-sought entries to address ChREBP via molecular glues^12, 13^. The “hub” protein 14-3-3 is involved in numerous signaling pathways, as well as in the pathophysiologic state of diabetes^14^. Molecular scaffold proteins belonging to the 14-3-3 protein family are widely conserved among eukaryotes^15, 16^. In mammals, this family comprises seven isoforms, namely, β, γ, ζ, η, τ, σ, and ε^17^. In the context of pancreatic β-cells, 14-3-3 proteins are integral to the regulation of insulin secretion, cell proliferation, and survival^14^. They contribute to maintaining glucose homeostasis and influence key aspects of the cell cycle, impacting β-cell mass^18, 19^. The diverse functions of 14-3-3 proteins encompass their involvement in mitochondrial activity^18^, cell cycle progression^14^, contribution to apoptosis and cell survival pathways^20^. ChREBP interacts with 14-3-3, *via* a PKA responsive phosphorylation site S196 of ChREBP, and also *via* an alpha helix in its N-terminal domain (residues 117-137), which is one of the few known phosphorylation-independent 14-3-3 binding motifs^10, 21, 22^. The molecular basis of this PPI has been studied by X-ray crystallography, revealing the presence of a free sulfate or phosphate ion binding in the 14-3-3 phospho-accepting pocket that interacts with both proteins (**Fig. S1a**)^23^. Adenosine monophosphate (AMP) has also been reported to bind to this phospho-accepting pocket, weakly stabilizing the protein complex and enhancing the 14-3-3-mediated cytoplasmic sequestration of ChREBP (**Fig. S1b**)^22^. Next to inorganic phosphate and AMP, ketone bodies regulate ChREBP activity by increasing its binding to 14-3-3^10, 22^. This may suggest that small-molecule stabilizers – molecular glues – of the ChREBP/14-3-3 PPI are potentially valuable tools to suppress glucolipotoxicity in T2D by inhibiting the transcriptional activity of ChREBP, thereby overcoming the current limitations of ‘direct’ targeting of TF functions.

MLXIPL, the gene that expresses ChREBP, produces two major splice isoforms: ChREBPα and ChREBPβ. The ChREBPα isoform is the full length-isoform, containing the low glucose inhibitor domain (LID, including the nuclear export signal (NES)) at its N-terminal region. One of the mechanisms that retains ChREBPα in the cytosol is *via* its interaction with 14-3-3 (**Fig. 1a**)^10, 22^. ChREBPβ lacks this N-terminal domain, resulting in a nuclear, constitutively active, hyper-potent TF^24, 25^. In healthy conditions, glucose flux leads to dissociation of CHREBPα from 14-3-3 resulting in its translocation to the nucleus and induction of ChREBPβ transcription, ultimately leading to glucose-stimulated β-cell proliferation to meet the demand for insulin (**Fig. 1b**). However, under prolonged hyperglycemic conditions, a robust positive feedback loop is triggered in which the newly produced ChREBPβ binds to its own promoter sites leading to even more ChREBPβ synthesis eventually resulting in glucolipotoxicity and β-cell apoptosis (**Fig. 1c**)^10, 22, 26^. These molecular mechanisms suggest that small molecule stabilization of the ChREBPα/14-3-3 PPI *via* a suitable molecular glue may prevent the nuclear import of ChREBPα under prolonged hyperglycemic conditions and thereby avert glucolipotoxicity (**Fig. 1d**). Indeed, our groups recently developed biochemically active stabilizers of the ChREBPα/14-3-3 PPI^12^; these compounds were only moderately active and more importantly lacked cellular activity, preventing cellular validation and downstream analyses of our hypothesis^12^. Here, we aimed to further improve these ChREBPα/14-3-3 PPI small-molecule stabilizers by using structure-activity relationship (SAR) analysis. This resulted in small-molecule stabilizers with a cooperativity factor (α) up to 220 for the ChREBPα/14-3-3 PPI. The introduction of a *geminal*-difluoro group into the pharmacophore not only significantly enhanced their stabilization efficiency, most probably by slight bending of the pharmacophore arrangement, but was also crucial for obtaining cellular permeation and activity. X-ray crystallography confirmed engagement of the molecular glues at the composite interface formed by the 14-3-3 binding peptide motif of ChREBPα and 14-3-3. Immunofluorescence showed retention of ChREBPα, by 14-3-3, in the cytoplasm of primary human β-cells upon addition of the stabilizers. Concomitantly, the transcriptional activity of ChREBPα was suppressed, hence preventing the upregulation of ChREBPβ in response to glucose and glucolipotoxicity. By exploiting this mechanism of action, β-cells were rescued from glucolipotoxicity, both in terms of viability and in terms of β-cell identity, including insulin production, by keeping ChREBP transcriptionally inactive at high glucose levels. This study provides the foundation for a potential new class of compounds for regulating ChREBP activity in T2D^12, 22^.

**Fig. 1:**
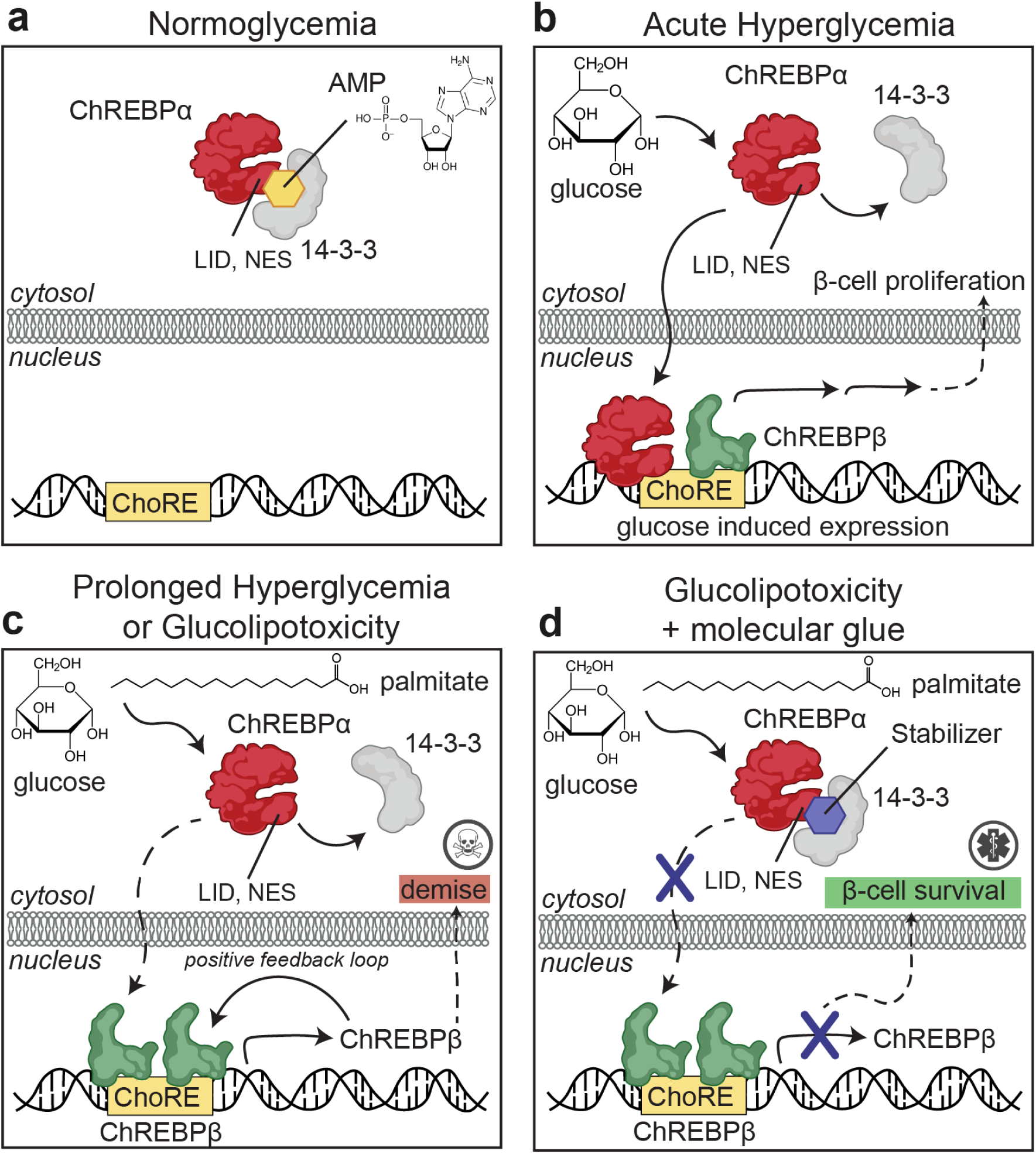
Protein-Protein Interaction between 14-3-3 and ChREBPα regulates β-cell fate. **a**. Under normoglycemic conditions, ChREBPα remains mostly cytoplasmic by binding to 14-3-3. ChREBPα is one of very few phosphorylation-independent 14-3-3 partner proteins and binds *via* a pocket containing a phosphate or sulfate ion, ketone, or AMP. **b.** In acute hyperglycemia, ChREBPα dissociates from 14-3-3 and transiently translocates into the nucleus where it binds multiple carbohydrate response elements (ChoREs) and promotes adaptive β-cell expansion. **c**. In prolonged hyperglycemia or hyperglycemia combined with hyperlipidemia (glucolipotoxicity), ChREBPα initiates and maintains a feed-forward surge in ChREBPβ expression, leading to β-cell demise. **d.** A novel class of molecular glue drugs specifically stabilize ChREBPα/14-3-3 interaction, prevent surge of ChREBPβ expression in glucolipotoxicity, and protect β-cell identity and survival.

## Results

### Differential stability and regulatory roles of ChREBP isoforms under high glucose

^24, 25^ ChREBPβ is a constitutively active transcription factor that promotes its own production by binding to its own promoter^24, 25^. This raises questions of how ChREBPβ production gets turned off and what is its relationship to ChREBPα once it begins the positive-feedback mechanism. To address this, we analyzed the two isoforms’ half-lives. ChREBPβ, under high glucose concentrations, exhibits substantial instability, with a half-life as short as 12 minutes. In contrast, ChREBPα displays increased stability, with its half-life extending beyond 30 minutes (**Fig. S2**). These findings suggest that without continuous priming by ChREBPα activity, ChREBPβ production would be self-limiting, and provides a mechanistic explanation of how the positive-feedback production of ChREBPβ can be turned off when glucose levels decrease, or when ChREBPα is prevented from entering the nucleus when glucose levels are high.

### Focused library based on 1 established a crucial SAR

To investigate the hypothesis that cytosolic sequestration of ChREBPα via molecular glue stabilization of the ChREBPα/14-3-3 PPI inhibits downstream glucotoxicity and β-cell apoptosis it was critical that our previously discovered stabilizer **1** (**Fig. 2a**) was first optimized to improve activity, but more importantly cellular efficacy. Stabilizer **1** was previously found to bind to the ChREBPα-peptide/14-3-3β interface (**Fig. S3a-c**)^12^. A challenge to optimization of **1** was the poor resolution of the ligand’s electron density in the X-ray co-crystal structure, particularly around the phenyl ring and linker of **1**. This poor electron density limited structure-guided optimization and indicated sub-optimal PPI stabilization (**Fig. S3d**). To address this technical challenge our efforts were directed towards the development of an alternative crystallization approach. Gratifyingly, this novel co-soak crystallization approach resulted in a new co-crystal structure with an improved resolution of 1.6 Å. The improved resolution enabled a more reliable model building, including the atomic fitting of the phenyl ring and ethylamine linker of **1**. (**Fig. 2a**, **S3e**). In addition to the phosphonate of **1** interacting with the phosphate-accepting pocket of 14-3-3 (R56, R129, Y130), and R128 of ChREBPα (**Fig. 2b**), it was now observed that both the phenyl and phenyl phosphonate rings of **1** make hydrophobic contacts with side chains of both 14-3-3 and ChREBPα. Further, an intramolecular hydrogen bond between the amide moiety and the phosphonate group of **1** was resolved. Finally, the improved resolution of this novel crystal structure enabled detection of a water-mediated hydrogen bond between the carbonyl amide of **1** and the main chain amide of I120 of ChREBPα (**Fig. 2b**). Subsequent Fluorescence Anisotropy (FA) measurements revealed that compound **1** stabilized the ChREBPα/14-3-3 complex with a cooperativity factor (α) of 35 (**Fig. 2c**). The cooperativity factor (α) and the intrinsic compound affinity to 14-3-3 (K_D_^II^) were determined by fitting, using a thermodynamic equilibrium model, which allows for an objective comparison of different stabilizers (**Fig. S4, S5**).^32^

**Fig. 2:**
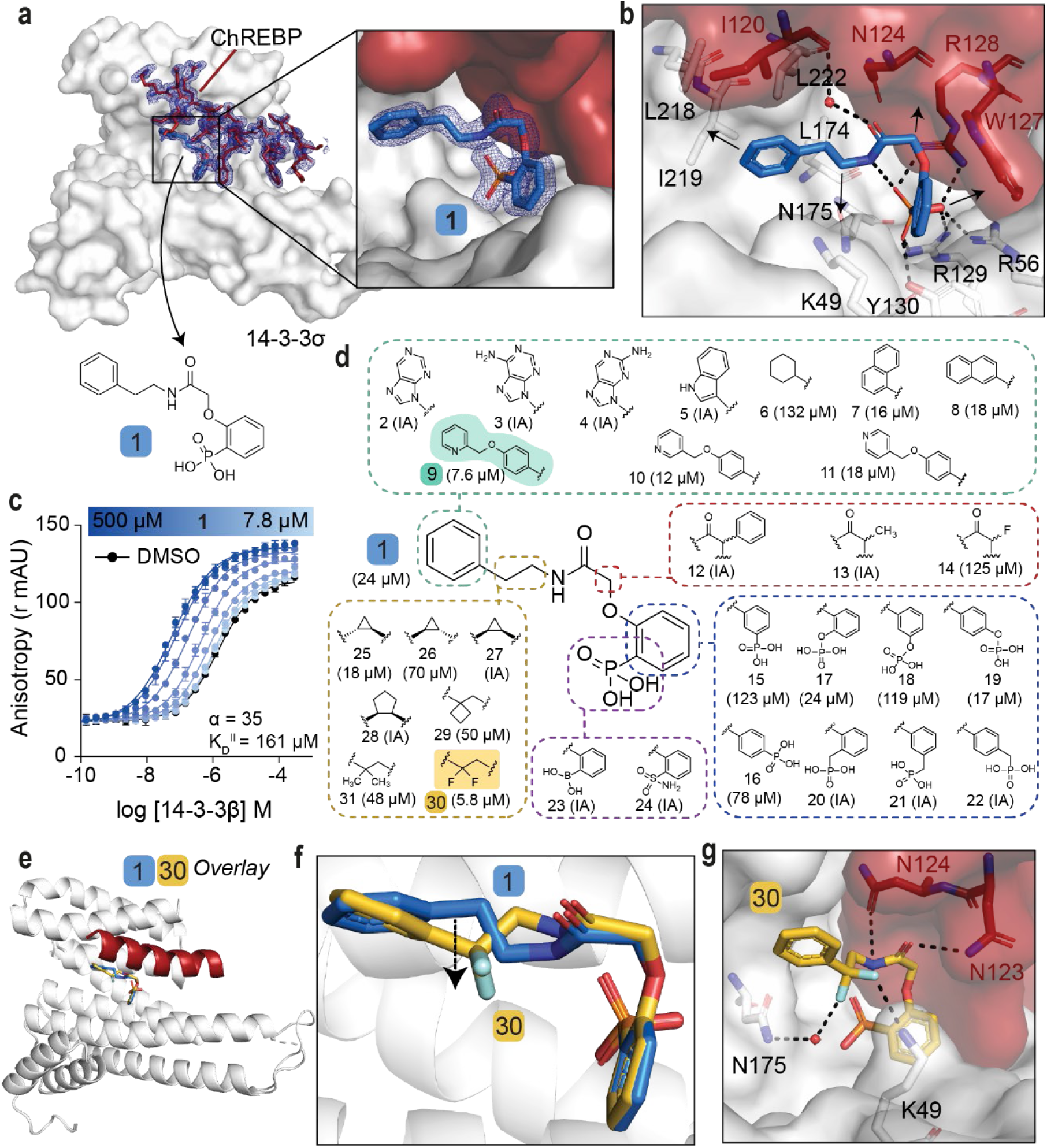
SAR around analog 1 results in improved stabilizer 30. **a**. Crystal structure of compound **1** (blue sticks) in complex with 14-3-3σ (white surface) and ChREBPα (red sticks and surface). Final 2F_o_-F_c_ electron density contoured at 1.0σ. **b.** Interactions of **1** (blue sticks) with 14-3-3σ (white) and ChREBPα (red) residues (relevant side chains are displayed in stick representation, polar contacts are shown as black dashed lines). **c.** FA 2D protein titration of 14-3-3β in FITC-labeled ChREBPα peptide (10 nM) and varied but fixed concentrations of **1** (0–500 µM), including the cooperativity factor (α, determined, by fitting, using a thermodynamic equilibrium model^32^) and intrinsic affinity of **1** to 14-3-3 (K_D_^II^). **d.** Structure and activity analogs of **1**. The two best compounds are marked in cyan and yellow. EC_50_ in parenthesis with mean ± SD, n = 2. For FA titration graphs see **Fig. S4, S5**. **e, f.** Crystallographic overlay of **1** (blue sticks) with **30** (yellow sticks) in complex of 14-3-3σ (white cartoon) and ChREBPα (red cartoon). **g.** Interactions of **30** (yellow) with 14-3-3 σ (white) and ChREBPα (red) (relevant side chains are displayed in stick representation, polar contacts are shown as black dashed lines).

With structural characterization of the ternary ChREBPα/14-3-3/**1** complex in hand, attention was shifted to the optimization of compound activity. To gain a greater SAR understanding, a focused library was synthesized aiming to improve the stabilization efficiency and selectivity for this class of molecular glues, in which compounds were compared based on their EC_50_ value derived from FA-based compound titrations (**Fig. 2d, S6, S7, S8**, **Table S1**) ^27–29^. To mimic the structure of the natural stabilizer **AMP**, the phenyl substituent of **1** was replaced by a purine (**2**, **3**, **4**). However, these modifications were not tolerated. Similarly, no stabilization was observed upon replacement of the phenyl moiety with either an indole (**5**), or a cyclohexane moiety (**6**), while naphthalene groups (**7**, **8**) led to slightly more active compounds. Extension of the phenyl group with a methylpyridine functional group (**9, 10, 11**) increased the stabilization efficiency, with the N-2 position (**9**, EC_50_ = 7.6 ± 0.5 µM) eliciting the greatest response. Substitutions at the central acetamide were not tolerated (**12, 13, 14**). Furthermore, systematic shifting of the phosphonate group along the phenyl phosphonate ring showed that the *ortho* position is most favorable (**1, 15, 16**). Interestingly, replacement of the phosphonate with a phosphate group was tolerated (**17**, **18**, **19**) and the *para* positioned analog led to improved compound activity (EC_50_ = 17.4 ± 3.2 µM). Conversion of the phenyl phosphonate into a benzyl phosphonate was however not tolerated (**20**, **21**, **22**). Phosphonate mimics, e.g. a boronic acid (**23**) or a sulfonamide (**24**) group were also inactive compounds. Lastly, we focused on modifications of the alkyl fragment of the phenyl ethyl amine moiety. Investigation of the SAR around the ethylene linker showed that replacement of the ethylene linker with a constrained (1*R*, 2*S*) cyclopropyl ring (**25**, EC_50_ = 17.6 ± 1.7 µM) elicited similar activity to **1**, while stereoisomers **26** (1*S*, 2*R*) and **27** (1*R*, 2*R*) were not tolerated. Furthermore, ring expansion to the cyclopentyl ring (**28**, 1*R*, 2*R*) was inactive and installation of a cyclobutyl (**29**) or *geminal*-dimethyl groups (**31**) resulted in diminished activity relative to **1**. Interestingly, the introduction of a *geminal*-difluoro group however significantly enhanced (six-fold relative to **1**) the stabilization of the ChREBPα/14-3-3 PPI, resulting in the best stabilization observed so far (**30**, EC_50_ = 5.8 ± 0.4 µM). Unfortunately, this analog was not amendable for phosphonate replacement with a sulfonamide or boronic acid, since this resulted in inactivity (**58**, **68**, **Table S2**). Merging the two best stabilizers (**9** and **30**) resulted in analog **73** with a similar EC_50_ value compared to each individual substitution (**73**, EC_50_ = 11.0 ± 1.5 µM, **Table S3**). Co-crystallization of ChREBPα and 14-3-3 was successful for **30**, showing a clear density of both the ChREBPα peptide and **30** (**Fig. S9a**). Overlaying the crystal structure of **30** with **1** revealed a high conformational similarity of 14-3-3σ and the ChREBPα peptide, with a comparable positioning of the phosphonate of **30** in the ChREBPα/14-3-3 phospho-accepting pocket (**Fig. 2e**, **2f**). The *geminal*-difluoro group of **30** was, however, directed downwards into the 14-3-3 binding groove, causing an alternative bend of the phenyl ethyl amine moiety (**Fig. 2f**). Both fluorine atoms form interactions with 14-3-3σ; one directly contacts K49 and the other engages in a water-mediated polar interaction with N175 (**Fig. 2g**), thereby recapitulating other literature reports showing that fluorinated small molecules can form potent hydrogen bonds ^27–29^. Additionally, due to the alternative bend of the linker, the amide of **30** was now positioned to interact with N123 and N124 of ChREBPα (**Fig. 2g**). These additional contacts of the molecular glue with both 14-3-3 and ChREBPα explain the increase in ternary complex formation by the introduction of the *geminal*-difluoro group.

### Fluorination of compounds improves stabilization potency

Encouraged by the enhanced stabilizing activity of the fluorinated analog **30**, a focused library of fluorinated compounds was next synthesized and compared to their non-fluorinated analogs (**Fig. 3a**, **Table S3**). Fluorination of analogs increased their stabilization potency consistently across all library members. Even the previously inactive *p*-Cl (**32**) and *p*-Br (**33**) substituted compounds now elicited a high activity when combined with the *geminal*-difluoro group substitution (**40:** EC_50_ = 6.6 ± 0.9 µM, **41**: EC_50_ = 4.1 ± 0.4 µM, respectively). The fluorine functionality has a common place in medicinal chemistry as reflected by 20-25% of drugs in the pharmaceutical pipeline containing at least one fluorine atom^30, 31^. Introduction of a fluoro group at the *meta* position of the phenyl ring even further improved compound activity (**43**: EC_50_ = 3.8 ± 0.2 µM). Additional to single fluoro-substitutions at the phenyl ring of **30**, double fluoro-substituted analogs were synthesized (panel IV in **Fig. 3a**). A 2,4-difluoro substitution resulted in the most potent stabilizer observed so far (**53**, EC_50_ = 3.1 ± 0.6 µM).

**Fig. 3:**
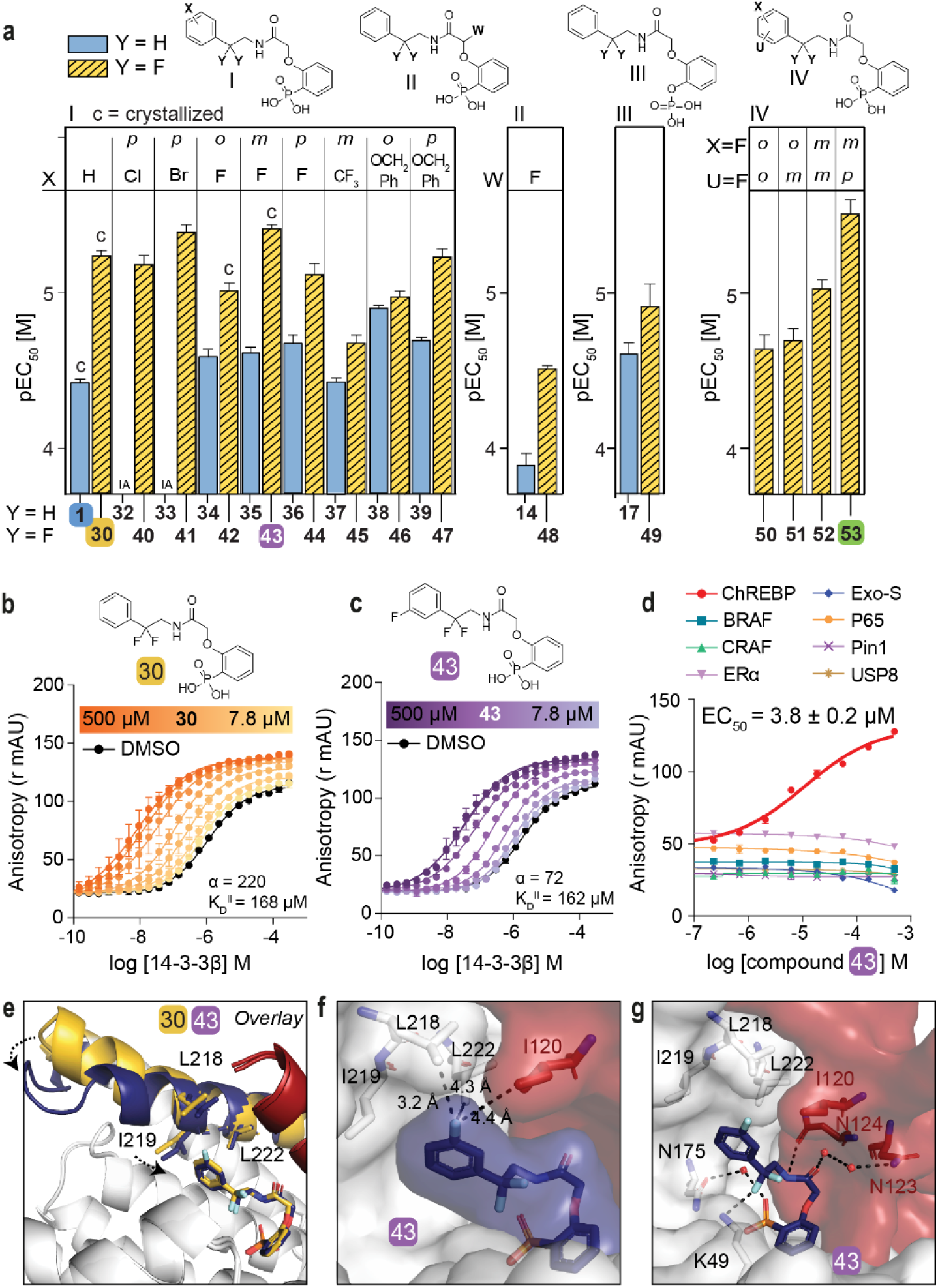
Fluorination of compounds enhances stabilizing potency. **a**. Structures and bar graphs of pEC_50_ values derived from FA compound titrations, for Y=H (blue bars) and Y=F (yellow bars). (For graphs see **Fig. S4**, **S5**, for EC_50_ values see **Table S3**) (mean ± SD, n=2). **b, c.** Titration of 14-3-3β to FITC-labeled ChREBPα peptide (10 nM) against varying fixed concentrations of **30** or **43** (0–500 µM) (mean ± SD, n = 2), including the cooperativity factor (α, determined, by fitting, using a thermodynamic equilibrium model^32^) and intrinsic affinity of the stabilizers to 14-3-3 (K_D_^II^). **d.** Selectivity studies by titrating **43** to 14-3-3β and eight different 14-3-3 interaction FITC-labeled peptides (all 10 nM) (mean ± SD, n = 2). **e.** Crystallographic overlay **30** (yellow) and **43** (purple) in complex with 14-3-3σ (white cartoon) and ChREBPα (red cartoon). Helix 9 of 14-3-3σ is colored in the same color as the corresponding compound, showing a helical ‘clamping’ effect when **43** (purple) is present. **f.** Surface representation of **43** (purple) in complex with 14-3-3σ (white) and ChREBPα (red), showing the distances (black dashes) of the **43** m-F substitution to the residues (sticks) of 14-3-3σ and ChREBPα. **g.** Interactions of **43** (purple) with 14-3-3σ (white) and ChREBPα (red) (relevant side chains are displayed in stick representation, polar contacts are shown as black dashed lines).

Two-dimensional FA titrations were performed to investigate the cooperativity in ChREBPα/14-3-3/stabilizer complex formation. 14-3-3β was titrated to FITC-labeled ChREBPα peptide (10 nM) in the presence of different, but constant, concentrations of **30**, **43** or **53** (0–500 µM) (**Fig. 3b**, **3c, S10a**). The change in apparent K_D_ of the ChREBPα/14-3-3 complex, over different concentrations of stabilizer (**Fig. S10b**), showed that all three fluorinated compounds (**30**, **43** and **53**) elicit a similar stabilizing profile, with a 2.4-to 6-fold increase in stabilization compared to the non-fluorinated parent analog **1**. Fitting the two-dimensional data using the thermodynamic equilibrium model^32^ showed increased cooperativity for all three fluorinated compounds (α = 220 (**30**), 72 (**43**), 126 (**53**)) compared to their defluorinated parent compound **1** (α = 35), while their intrinsic affinity for 14-3-3 was barely affected (K_D_^II^ = 168 µM, 162 µM, 110 µM for **30**, **43**, **53** and 161 µM for **1**) (**Fig. 3b**, **3c**, **S5**, **S10a**). This indicates that the compound optimization has mainly improved the interaction with ChREBPα and not the binding affinity to 14-3-3. Of note, the mean squared error landscape of the two fitted parameters showed for some stabilizers that the α and K_D_^II^ parameters are interconnected, meaning that a weaker intrinsic affinity, K_D_^II^, correlates to an increased cooperativity factor, α. (**Fig. S5**).

Phosphate- and phosphonate-based compounds may also act as inhibitors of other 14-3-3-client protein complexes, often acting as low micromolar (IC_50_ ∼ 1-20 µM) inhibitors^33, 34^. To test the specificity of the ChREBPα/14-3-3 PPI stabilizers, we selected an array of eight representative 14-3-3 client-derived peptide motifs with differentiating binding sequences and included internal binding motifs BRAF^35^, CRAF^36^, p65^37^, USP8^38, 39^, one C-terminal binding motif ERα^40^, one special binding mode Pin1^41^ and another reported non-phosphorylated motif ExoS^21^. Strikingly, all three compounds (**30**, **43**, **53**) displayed a high selectivity for stabilizing ChREBPα, without affecting any other client peptide up to 100 µM (**Fig. 3d**, **S10c**). Furthermore, titration of all seven 14-3-3 isoforms to ChREBPα in the presence of **43** showed PPI stabilization among all isoforms (**Fig. S11**). The ChREBPα stabilization selectivity of 43 was also preserved among different 14-3-3 isoforms (**Fig. S12**). These data demonstrate the highly selective nature of the molecular glue activity of these compounds by addressing a unique pocket only present in the ChREBPα/14-3-3 complex and via a stabilization mechanism, involving a large cooperativity effect.

Crystallization of the new fluorinated compounds with the ChREBPα/14-3-3σ complex was also successful for **42** and **43**, resulting in a clear density for the entire molecule (**Fig. S9b, c**). A crystallographic overlay with **30** revealed an additional conformational ‘clamping’ effect of helix-9 of 14-3-3σ in the presence of **43** (**Fig. 3e**), causing the hydrophobic residues of 14-3-3 (I219, L218) to be closer to the phenyl ring of **43**. This clamping effect of helix-9 is in line with previous observations for molecular glues with high cooperativity factors, for other 14-3-3 PPIs ^42^. A more detailed analysis showed that the *meta*-fluoro group of **43** is positioned at the rim of the hydrophobic interface of 14-3-3 and ChREBPα (**Fig. 3f**). The rest of stabilizer **43** has similar interactions with both ChREBPα and 14-3-3 as **30** (**Fig. 3g**, **S13**). An *ortho-* fluoro substitution at the phenyl ring was less favorable (**42**, EC_50_ = 9.6 ± 1.1 µM) although it did result in a similar ‘clamping’ effect of helix-9 of 14-3-3 (**Fig. S14**). This *ortho* substituted fluoro was not positioned at the rim of the ChREBPα/14-3-3 interaction interface, instead it made a polar interaction with N175 of 14-3-3 (**Fig. S14**). While no crystal structure could be solved for the double fluoro-substituted analog **53**, the structure of **43** shows room for a fluoro-substitution at the *para* position of its phenyl ring, explaining the high potency of **53** (**Fig. 3g**). Likely, the electron withdrawing effects of the fluorine substitutions are enhancing hydrophobic contacts with the hydrophobic amino acids at the roof of the 14-3-3 groove. Concluding, fluorination of the scaffold enhanced stabilizing efficiency of the ChREBPα/14-3-3 complex, by strengthening the interactions at the rim of the PPI interface.

### Molecular glues stabilizing the ChREBPα/14-3-3 interaction rescue human β-cells from glucolipotoxic death

We tested six compounds (**1**, **12**, **30**, **43**, **53**, **66**) for cytotoxicity and their ability to protect β-cells from glucolipotoxicity (summarized in **Fig. 4a**). Compound **1**, our previously reported parent compound that was moderately active in the biochemical assay, displayed cytotoxicity in INS-1 cells. In contrast, compounds **30**, **43** and **53**, which were all active in biochemical assays, demonstrated no cytotoxicity in INS-1 cells and protected the β-cells from glucolipotoxicity, turning them into candidates for further cellular evaluation. Compounds **12** and **66**, which served as control compounds, were not active in the biochemical assay and were also neither generally cytotoxic nor rescued the cells from glucolipotoxicity. To test the efficacy of our compounds in mitigating β-cell death from glucolipotoxicity, we conducted dose-response experiments in rat insulinoma, β-cell-like INS-1 cells^43^. We observed a significant attenuation of cell death in response to glucolipotoxic conditions at 5 and 10 μM of active compounds **30**, **43**, and **53**, but not in the presence of the inactive compounds **12** or, **66** (**Fig. S15a**). To investigate cadaveric human islets, we optimized the single-cell and population-level analyses using real-time kinetic labeling (SPARKL) assay^44^, that measures the kinetics of overall proliferation (cell count, using NucBlue) or cell death rates (using YoYo3) specifically in β-cells using the rat insulin promoter (RIP)-ZSGreen adenovirus^45^ (**Fig 4b**, **Fig S15a-c**). The active compounds (**30**, **43**, **53**) all exhibited a remarkable decrease in β-cell death in glucolipotoxic conditions, as observed in **Fig 4c**, **4d, 4e** while the ‘inactive’ control compounds showed no activity. Notably, for all five tested compounds, there was no discernible impact on β-cell number, indicating that over the course of 72 h, no meaningful proliferation occurred in treatment groups (**Fig S15b-c**). We also performed glucose-stimulated insulin secretion (GSIS) in primary human islets pre-treated for 24 h with **43,** in low or high glucose conditions. Remarkably, we found that **43** improved glucose-stimulated insulin secretion (GSIS) stimulation index (SI) in high glucose (**Fig 4f**) with no effect on SI in response to KCl or on insulin content (**Fig S15d-e**).

**Fig 4:**
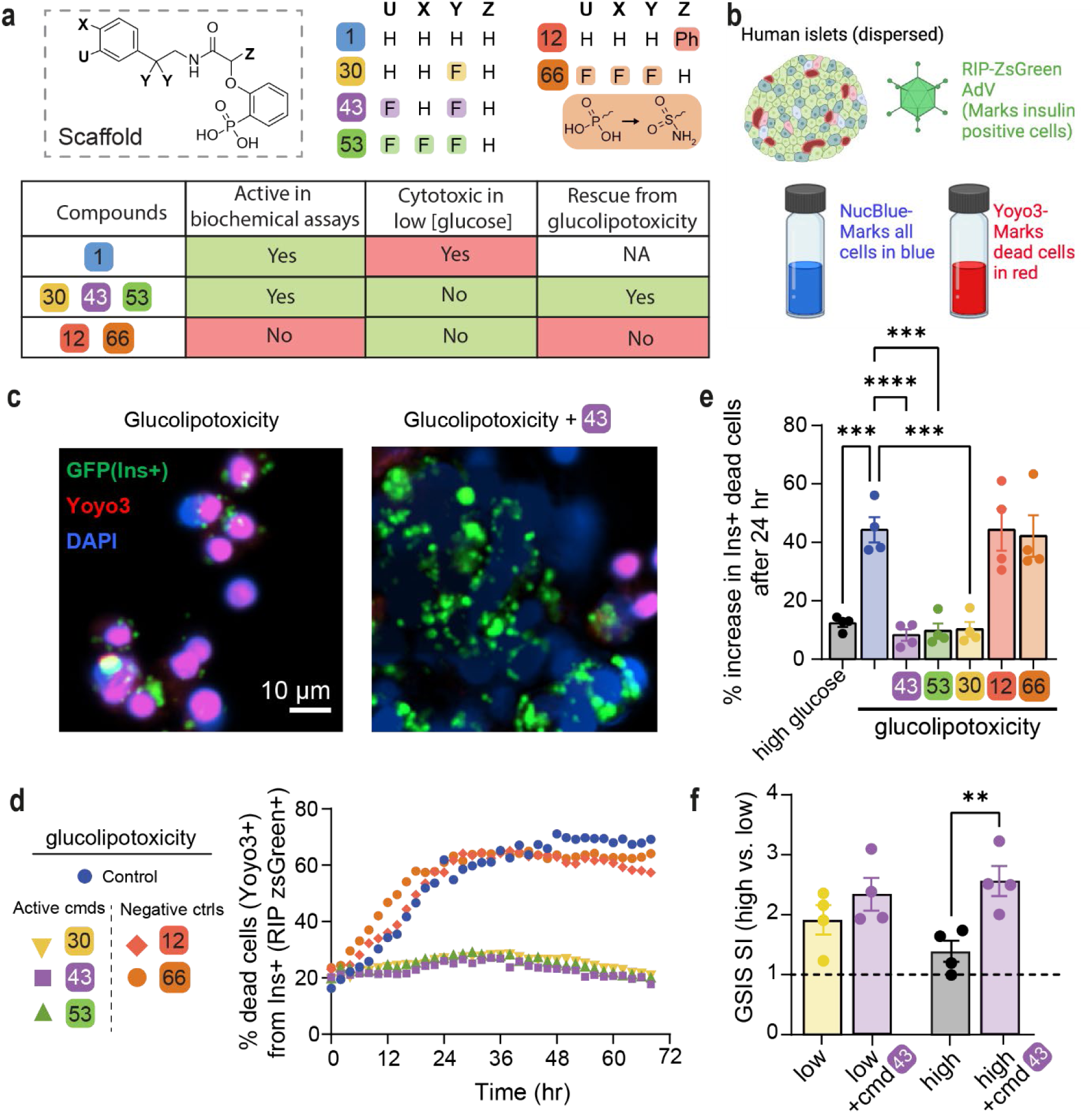
Active compounds protect human β-cells from glucolipotoxicity. **a.** Overview of compounds included in cellular assays. Table shows results of cytotoxicity and β-cell rescue from glucolipotoxicity in the presence of the compounds (green indicates positive outcome and red cytotoxicity). **b.** Schematic of adaptation to the SPARKL assay in human islets to specifically monitor β-cells. **c.** Representative figures from **d** at 48 h with **43**. The results are representative from 4 different human cadaveric donors. **d.** Representative kinetics of β-cell death in glucolipotoxicity (20 mM glucose+500 μM palmitate), in the presence of 10 µM of the indicated compounds. **e.** Quantification of β-cell death (assessed by Yoyo3+% of GFP+ cells) at 24 h from **d**. **f.** Human islets were treated for 24 h as indicated, followed by quantification of glucose-stimulated insulin secretion (GSIS) in KREBS buffer (2.8 mM glucose, 1% BSA) over 30 min. The corresponding GSIS-stimulation index (SI) was obtained by determining the ratio of insulin release at high vs low glucose. Data are means +/- SEM; n=4; p<0.01**, p<0.01; ***, p<0.005; ****, pP<0.001.

Our method was validated using TUNEL staining at 48 h using 10 µM **43** to show similar cell death patterns as the kinetic measurement (**Fig S16**). Indeed, **43**, which we focus on for the rest of this study, also prevented glucose-stimulated proliferation, confirming our previous studies^8, 25, 46^ and glucolipotoxic cell death in INS-1 cells (**Fig S17**). Together, these studies demonstrated that the three active compounds (**30**, **43**, **53**) prevent β-cell death while maintaining and even enhancing β-cell function in the context of glucolipotoxicity. Importantly. we also demonstrated that the effects of **43** were mediated by ChREBPβ as silencing of ChREBPβ had a similar effect on protection from glucolipotoxicity in human islet cells, while overexpression of ChREBPβ led to increased cell death under low, high, and glucolipotoxic conditions, and **43** was unable to protect human islet cells from apoptosis due to overexpression of ChREBPβ (**Fig S18**).

### Compound 43 stabilizes the ChREBPα/14-3-3 interaction *in situ* preventing ChREBPα’s nuclear translocation in response to glucose or glucose + palmitate

To explore the mechanism by which **43** acts *in cellulo* as a molecular glue, we performed three sets of experiments. First, a proximity ligation assay (PLA) was employed in INS-1 cells using an antibody against ChREBPα and a pan 14-3-3 antibody. PLA assays show a fluorescent signal only when proteins are in close (<40 nm) and stable proximity to each other ^47^. In low glucose, we observed an interaction between 14-3-3 and ChREBPα. By contrast, following exposure to high glucose for 30 min, ChREBPα no longer interacted with 14-3-3, but the interaction was restored after 2 h (**Fig 5a**). These observations are consistent with our previous study showing that ChREBPα transiently enters the nucleus to begin the feed-forward induction of ChREBPβ ^25^. Remarkably, in the presence of **43**, the interaction between 14-3-3 and ChREBPα remained unchanged at high glucose concentrations, thus confirming that **43** stabilized the interaction between 14-3-3 and ChREBPα in high glucose. In a second set of experiments, in which nuclear localization of ChREBPα was studied using a ChREBPα-specific antibody, we again found the same dynamics; ChREBPα entered the nucleus after 30 min of high glucose and exited the nucleus after 2 h. However, **43** blocked the transient translocation of ChREBPα in response to glucose (**Fig. 5b**, **5c**). To confirm specificity of 14-3-3 binding to ChREBPα, we also studied the dissociation of 14-3-3 from NFATC1, previously demonstrated to translocate into the nucleus in response to glucose stimuli ^48, 49^. In the presence and absence of **43,** glucose promoted 14-3-3 dissociation from NFATC1, demonstrating the selectivity of our molecular glue to stabilize the interaction of ChREBPα and 14-3-3 (**Fig S19**). Under glucolipotoxic conditions (20 mM glucose + 500 µM palmitate), we observed a similar translocation of ChREBPα to the nucleus after 30 min, however, the nuclear clearance of ChREBPα under glucolipotoxic conditions took longer compared to high glucose alone (**Fig. 5d**, **5e**). AMPK activity cannot mediate **43**’s effects since in the presence of 1 mM AICAR, a widely used AMPK activator, translocation of ChREBPα is not blocked (**Fig 5c**, **5e**).

**Fig. 5:**
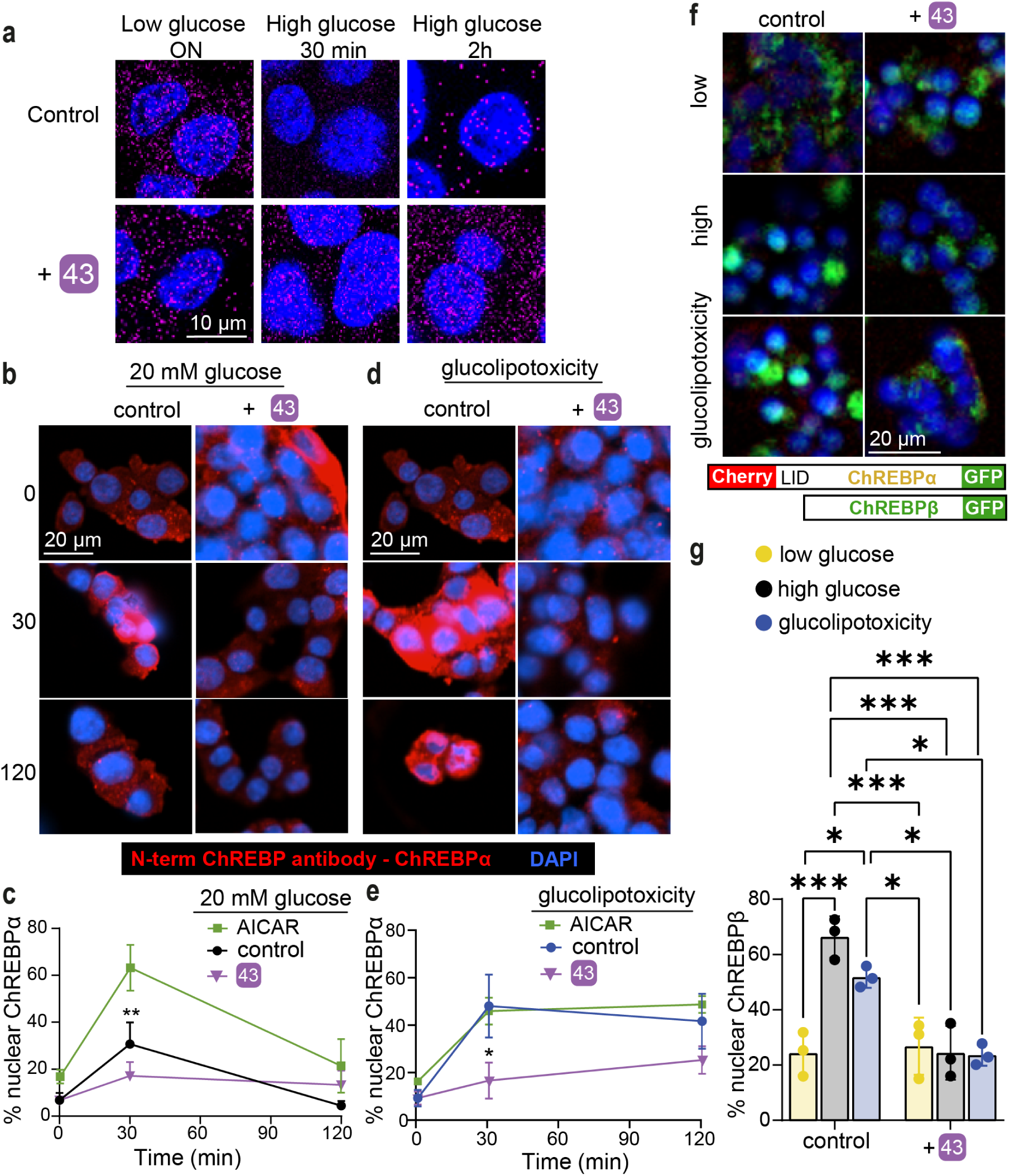
43 stabilizes ChREBPα/14-3-3 interaction and thus retains cytoplasmic ChREBPα localization in response to glucose and glucolipotoxicity. **a.** Proximity ligation assay demonstrating increased interaction between 14-3-3 and ChREBPα. INS-1 cells were cultured overnight (ON) at low (5.5 mM) glucose and exposed to high (20 mM) glucose for the indicated times **b, d.** Representative figures showing the nuclear localization of ChREBPα after exposure to high glucose (**b**) or glucolipotoxic (**d**) conditions. **c, e.** Time course of nuclear localization of ChREBPα based on figures **b**, **d**, respectively in addition to 1mM AICAR. **f, g.** CRISPR/Cas9 engineered INS-1 cells treated with the indicated compounds for 24 h; Low-5.5 mM glucose, High-20 mM glucose, glucolipotoxicity-20 mM glucose+500 μM palmitate. **f.** Representative images at 24 h. **g.** Quantification of % nuclear ChREBP at 24 h. Data are the means +/-SEM, n=3-5, *p<0.05, **p<0.01.

Because our nuclear translocation studies are based on immunodetection, this positive outcome could be a result of epitope masking, e.g. by a ChREBPα interaction with the transcription machinery or other heteropartners, which could block antibody binding. Therefore, a third approach used CRISPR/Cas9 engineered INS-1 cells ^2525252525252525252525252525^ wherein fluorescent tags to the 5′ and 3′ ends of ChREBP were added^25^. An mCherry tags the N-terminal LID domain, which identifies ChREBPα exclusively, and an eGFP tag on the C-terminus represents both ChREBPα and ChREBPβ. In these cells, ChREBPα appears as yellow (red+green), and ChREBPβ appears green. Compound **43** clearly prevents the built up of nuclear ChREBPβ in both high glucose and glucolipotoxic conditions, thus confirming the immunostaining results (**Fig. 5f**, **5g**). Together, these studies demonstrate that **43** stabilizes the interaction of 14-3-3 and ChREBPα under conditions of hyperglycemia and glucolipotoxicity.

### Effect of stabilizers on ChREBP downstream genes and preservation of β-cell identity

Next, we assessed the effect of stabilizers on transcription of the ChREBP splice isoforms in human islets. While ChREBPα mRNA levels remained relatively stable in all conditions tested (**Fig. 6a**), ChREBPβ, which contains a well characterized carbohydrate response element (ChoRE) on its promoter^24, 46^, was upregulated in response to high glucose, or to high glucose and high palmitate (glucolipotoxicity). Interestingly, compound **43** prevented upregulation of ChREBPβ both at mRNA level (**Fig. 6b**) and at protein level (**Fig. 6c**, **6d**), indicating that inhibition of the nuclear translocation of ChREBPα by strengthening the ChREBPα/14-3-3 interaction with **43** is required for the blocking of the upregulation of glucose responsive genes such as ChREBPβ. TXNIP is another glucose responsive gene, which also contains a well characterized ChoRE^50^ and its upregulation is implicated in oxidative stress and β-cell death^51^. Promoter luciferase assays demonstrated that **43** prevents the glucose response of the TXNIP gene (**Fig. 6e**). Importantly, β-cell identity marker genes, INS (**Fig. 6f**) and PDX1 (**Fig. 6g**), were both downregulated at the mRNA level in human islets exposed to glucolipotoxicity, consistent with the previously reported de-differentiation phenotype pertinent to β-cell demise^52, 53^. Yet, in the presence of **43** this de-differentiation was prevented (**Fig. 6f**, **6g**). Similarly, immunostaining for PDX1 showed a marked decrease in PDX1 protein levels in glucolipotoxicity, which was rescued by **43** (**Fig. 6c**, **6h**). We also tested by qPCR in human islets other known 14-3-3 interactors, including USP8, RELA, PIN1, NFATC1 and BRAF (**Fig S20a-e**). Other β-cell maturity markers or glucose sensing and processing genes were unchanged by **43** (**Fig S20f-i**). However, the β-cell dedifferentiation marker-ALDH1A3^54^ was significantly decreased by **43** in low glucose concentrations but unaffected in high glucose or glucolipotoxicity (**Fig S20j**). To test functional consequences, we measured the effects of **43** on Ca^+2^ signaling in INS-1 cells, where **43** had no effect on calcium upregulation in response to glucose (**Fig S21a**). However, **43** enhanced mitochondrial activity in high glucose as evidenced by both active mitochondria staining (**Fig S21b**) and ATP/ADP ratio (**Fig S21c**). Together these data suggest that silencing ChREBPβ by **43** decreases *de novo* lipogenesis leading to less accumulation of toxic lipid species like ceramides, improving mitochondrial function.

**Fig. 6:**
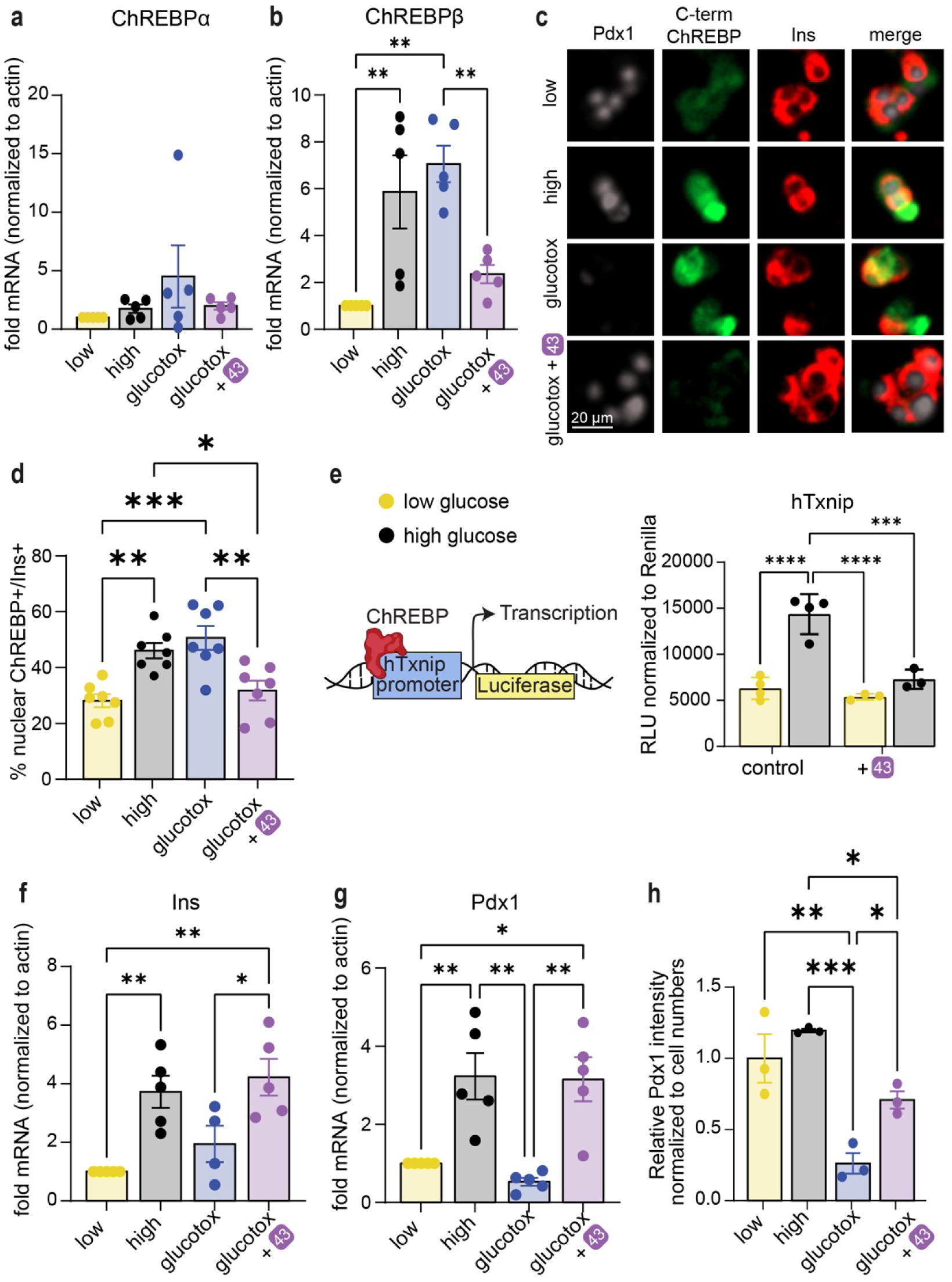
43 preserves β-cell identity in glucolipotoxicity and prevents upregulation of ChREBPβ in high glucose and glucolipotoxicity. **a,b,f,g.** mRNA fold-enrichment over control low glucose in human islets, treated with 10 µM **43** for 24 h; Low-5.5 mM glucose, High-20 mM glucose, glucotox-20 mM glucose+500 μM palmitate. **c, d, h**. Immunostaining for Pdx1, c-term ChREBP, and insulin in dispersed islets treated for 48 h with the indicated treatments. **e.** INS-1 cells expressing luciferase driven by the human TXNIP promoter were incubated for 24 h at the indicated glucose concentrations, in the presence or absence of 10 μM **43**. Data are the means +/-SEM, n=3-7, *p<0.05, **p<0.01***, p<0.005, ****, p<0.001.

## Discussion

The global epidemic of T2D necessitates the development of novel therapeutic and preventive strategies to arrest the progressive nature of this disease^55^. ChREBP is an important regulator of glucose levels and is increasingly recognized as a potential target for T2D treatment^56^. A small molecule regulation of its function is however challenging as TFs are notoriously difficult to target with ‘classical’ small molecules. Here, we sought to develop a molecular glue approach, based on the use of novel small molecule PPI stabilizers, molecular glues, for the ChREBPα/14-3-3 PPI. These compounds were then used to test if retention of ChREBPα in the cytoplasm may lead to a persistent, chemotherapeutically exploitable inactivation of its transcriptional activity. Through the development of cellularly active molecular glues, we were able to demonstrate that this unconventional strategy for targeting TF functions *via* TF retention in the cytoplasm indeed resulted in the desired, consistent, and efficacious glucolipotoxicity rescue phenotype. We also did not observe significant cytotoxicity on a cellular level. Besides its implications for T2D therapy, we therefore believe that 14-3-3 or other scaffold-mediated retention of TFs in the cytoplasm *via* customized molecular glues may represent a broader therapeutic approach that may also be extended to other, difficult to target TFs with medicinal relevance. This strategy also complements other non-conventional ‘molecular glue’ or PROTAC TF approaches such as TRAFTACs or other TF targeted degradation systems^57, 58^.

Sequence alignments^59^ of the interacting regions between 14-3-3 proteins and ChREBPα display the conserved sequences of the α2 and α3 helices of ChREBPα across multiple species, including human, chimpanzee, gorilla, rat, and mouse (**Fig S22a)**. The alignment of specific 14-3-3 helices (helices 9, 7, and 3) that interact with ChREBPα’s α2 helix, highlighted in red within the crystal structure model demonstrates sequence conservation across the seven human 14-3-3 isoforms (**Fig S22b top**). Cross-species alignment (human, chimpanzee, gorilla, rat, and mouse) for the 14-3-3 sigma isoform underscores the evolutionary conservation of interaction domains (**Fig S22b bottom)**, suggesting functional relevance in ChREBPα’s regulation by 14-3-3 proteins.

Our cellular active molecular glues simultaneously engage both protein partners in the composite ChREBPα/14-3-3 binding pocket, causing a cooperativity factor up to 220 for this PPI. The addition of *geminal*-difluoro groups increased the PPI stabilization efficiency significantly, overall leading to compounds with micromolar cellular potency. On a molecular level, this enhanced PPI stabilization by fluorination of the molecular glues was achieved by stimulating the clamping of helix-9 of 14-3-3 and contacting hydrophobic amino acids at the interaction surface of the ChREBPα/14-3-3 complex. This molecular mechanism also resulted in a very high selectivity of the compounds for the stabilization of the ChREBPα/14-3-3 PPI over other 14-3-3-based PPIs. Selective interfacing of these molecular glues with amino acid residues that only occur in the context of this specific PPI complex, thus result both in a large cooperative effect for PPI stabilization and high selectivity. Combined, this high cooperativity and selectivity for the ChREBPα/14-3-3 PPI, and probably increased membrane permeability, conferred cellular activity to our lead compound.

Our studies also shed further light on the function and suitability of ChREBP as a target in metabolic disease. As T2D progresses, there is a progressive decline in β-cell mass due to the metabolic overload associated with a diabetic environment. ChREBP has long been identified as a potential mediator of this decline, in part by activating the pro-oxidative protuberant, TXNIP^50^. Indeed, clinical trials have been launched with compounds that inhibit the induction of TXNIP in T1D^60^. We recently described the destructive feed-forward production of ChREBPβ in β-cells in the context of prolonged hyperglycemia and diabetes ^25, 60–62^. Unrestrained ChREBPβ leads to increased oxidative stress, which in turn leads to decreased expression of transcription factors necessary to maintain β-cell identity and function. Importantly, we found that deletion of the ChREBPβ isoform, using the Cre/Lox system, completely rescues β-cells from cell death and dedifferentiation due to glucolipotoxicity^25^. Here, we found molecular glues, which prevented ChREBPα from dissociating from 14-3-3 and initiating the feed-forward production of ChREBPβ, that were also remarkably effective at preventing cell death and dedifferentiation from glucolipotoxicity. One additional consequence of the retention of ChREBPα was that the molecular glues attenuated cell proliferation (**Fig. S17f**). This result was consistent with our previous findings that a modest induction of ChREBPβ is required for adaptive β-cell expansion in mice^8, 25, 46^. Considering molecular glues as a potential therapeutic, it is likely that preservation of β-cell mass is more clinically relevant compared to the loss of an extremely low proliferation rate of human β-cells^63, 64^.

ChREBPα may be anchored in the cytoplasm by elements other than 14-3-3 including sorcin, a Ca^++^-binding ER protein, and *via* C-terminal amphiphilic interactions with lipid droplets ^65, 66^. In addition, numerous post translational modifications^6, 67^ as well as interaction with importin α^68^ may play important roles in regulating ChREBP cellular location. Since 14-3-3 and importin α compete for the same ChREBP binding region, ChREBP/importin α-PPI inhibitors, as developed by Uyeda *et al*.^69^ could potentially work synergistically with ChREBPα/14-3-3 stabilizers in suppressing the progression of T2D. Importantly, upregulation of ChREBPβ and downstream genes (i.e. TXNIP and/or DNL genes) are implicated not only in β-cell demise but also in kidney failure^70^, cardiac hypertrophy^71^, ischemic and cardiovascular diseases^72, 73^, liver steatosis^74^, and NAFLD^75^ in particular in high glucose setting. Inhibiting ChREBPβ and downstream genes might prove to be valuable not only for preserving β-cell mass but may also be therapeutic in other tissues for the previously listed conditions.

To summarize, we demonstrated the development of highly potent, selective, active stabilizers of the ChREBPα/14-3-3 PPI with a phosphonate chemotype that display activity in both INS-1 cell lines and primary human islets. This new compound class possibly presents the first foundations for the development of novel therapeutics against T2D. More broadly, this work delineates a novel conceptual entry to the forthcoming discoveries of molecular glues, and we envision that 14-3-3 PPI stabilization can not only be applied for the interaction with ChREBP but can be translated to other TFs as well and offers an orthogonal strategy for hard-to-drug proteins and pathways in general.

## Online content

Any methods, additional references, Nature Portfolio reporting summaries, source data, extended data, supplementary information, acknowledgements, peer review information; details of author contributions and competing interests; and statements of data and code availability are available upon request

## Methods

### Protein expression and purification

The 14-3-3βFL and 14-3-3σΔC isoform (full length and truncated C-terminus after T231 (ΔC to enhance crystallization)) containing a N-terminal His6 tag were expressed and purified as described previously^12^.

### Peptide Sequences

The N-terminal FITC labeled ChREBP-derived peptide (residues 117 – 142; sequence: RDKIRLNNAIWRAWYIQYVKRRKSPV-CONH_2_) was synthesized via Fmoc solid phase peptide synthesis as described previously^12^. N-terminal acetylated ChREBP peptide used for crystallization was purchased from GenScript Biotech Corp. Peptides used for selectivity studies were purchased from GenScript Biotech Corp with the following sequences: BRAF: (5-FAM-RDRSS(pS)APNVH-CONH_2_), CRAF: (5-FAM-QRST(pS)TPNVH-CONH_2_), ERα: (5-FAM-AEGPFA(pT)V-COOH), EXO-S: (5-FAM-KKLMFK(pT)EGPDSD-CONH_2_), P65: (5-FAM-EGRSAG(pS)IPGRRS-CONH_2_), PIN1: (5-FAM-LVKHSQSRRPS(pS)WRQEK-CONH_2_), USP8: (5-FAM-KLKRSY(pS)SPDITQ-CONH_2_).

### Fluorescence Anisotropy measurements

Fluorescein labeled peptides, 14-3-3βFL protein, the compounds (100 mM stock solution in DMSO) were diluted in buffer (10 mM HEPES, pH 7.5, 150 mM NaCl, 0.1% Tween20, 1 mg/mL Bovine Serum Albumin (BSA; Sigma-Aldrich). Final DMSO in the assay was always 1%. Dilution series of 14-3-3 proteins or compounds were made in black, round-bottom 384-microwell plates (Corning) in a final sample volume of 10 µL in duplicates.

Compound Titrations were made by titrating the compound in a 3-fold dilution series (starting at 500 µM) to a mix of FITC labeled peptide (10 nM) and 14-3-3β (concentration at EC_20_ value of protein-peptide complex; 150 nM for ChREBP). For the selectivity studies concentration of 14-3-3β were: 300 nM for ERα, 3.65 µM for Exo-S, 10 µM for Pin1, 250 nM for USP8, 230 nM for B-RAF, 30 µM for P65 and 450 nM for C-RAF. Fluorescence anisotropy measurements were performed directly.

Protein 2D titrations were made by titrating 14-3-3β in a 2-fold dilution series (starting at 300 µM) to a mix of FITC-labeled peptide (10 nM) against varying fixed concentrations of compound (2-fold dilution from 500 µM), or DMSO. Fluorescence anisotropy measurements were performed directly.

Fluorescence anisotropy values were measured using a Tecan Infinite F500 plate reader (filter set lex: 485 ± nm, lem: 535 ± 25 nm; mirror: Dichroic 510; flashes:20; integration time: 50 ms; settle time: 0 ms; gain: 55; and Z-position: calculated from well). Wells containing only FITC-labeled peptide were used to set as G-factor at 35 mP. Data reported are at endpoint. EC_50_ and K_D_ values were obtained from fitting the data with a four-parameter logistic model (4PL) in GraphPad Prism 7 for Windows. Data was obtained and averaged based on two independent experiments.

### Cooperativity Analysis

The cooperativity parameters for compound 1, 30, 43 and 53 for 14-3-3 and ChREBP were determined by using the thermodynamic equilibrium system as described previously^76^. The data from 2D-titrations was provided to the model including the K_D_^I^ = 1.5 µM, P_tot = 10 nM, and the variable concentrations of 14-3-3 and stabilizer at each data point. Fit parameters were given the following initial guess values: K_D_^II^: 500 µM, α: 100.

### X-Ray crystallography data collection and refinement

A mixture of Ac-ChREBP peptide and compound (2 µL of 5 mM stock ChREBP peptide + 0.5 µL of 100 mM stock compound) was mixed in crystallization buffer (CB: 20 mM HEPES pH 7.5, 2 mM MgCl_2_, 2 mM βME) to a final concentration of 4 µL. This was added to preformed 14-3-3σ/peptide 2d crystals at 4 °C that were produced as described previously^13^. Single crystals were fished after 7 days of incubation at 4 °C, and flash-cooled in liquid nitrogen. Diffraction data was collected at either the Deutsche Elektronen-Synchrotron (DESY Petra III beamline P11 proposal 11011126 and 11010888, Hamburg, Germany (crystal structures of compound 1 and 30), or at the European Synchrotron Radiation Facility (ESRF Grenoble, France, beamline ID23-2 proposal MX2268) (crystal structures of compound 43 and 42). Initial data processing was performed at DESY using XDS^77^ or at ESRF using DIALS^78^ after which pre-processed data was taken towards further scaling steps, molecular replacement, and refinement.

Data was processed using the CCP4i2 suite^79^ (version 8.0.003). After indexing and integrating the data, scaling was done using AIMLESS^80^. The data was phased with MolRep^81^ using 4JC3 as a template. Presence of soaked ligands and ChREBP-peptide was verified by visual inspection of the Fo-Fc and 2Fo-Fc electron density maps in COOT ^82^ (version 0.9.6). If electron density corresponding to ligand and peptide was present, the ChREBP peptide was built in and the structure and restrains of the ligands were generated using AceDRG^83^, followed by model rebuilding and refinement using REFMAC5^84^. The PDB REDO^85^ server (pdb-redo.edu) was used to complete the model building and refinement. The images were created using the PyMOL Molecular Graphics System (Schrödinger LLC, version 2.2.3). For refinement statistics see Table S4.

The structures were deposited in the protein data bank (PDB) with IDs: 8BTQ (compound 1), 8C1Y (compound 30), 8BWH (compound 42) and 8BWE (compound 43).

### Synthesis of SAR library for 1

Detailed synthetic procedures and characterization of compounds marked with a star (*) in Table S1 were described previously^12^. The synthetic procedure and characterization of all remaining compounds are provided in the Supplemental Methods. Generally, all compounds obtained had been purified by reversed-phase high performance liquid chromatography (HPLC) and characterized by liquid chromatography mass spectrometry (LC-MS) and nuclear magnetic resonance (NMR; 400 MHz for ^1^H NMR and 100 MHz for ^13^C NMR). Compounds were prepared as 100 mM stock solutions in DMSO before use in experiments, and stored at -20 °C.

### Cell lines

INS-1–derived 832/13 rat insulinoma cells were maintained in RPMI 1640 medium with 10% FBS, 10 mM HEPES, 2 mM L-glutamine, 1 mM sodium pyruvate, and 50 mM β-mercaptoethanol 100U/mL penicillin, 100 mg/mL streptomycin and further supplemented with 11 mM glucose, at 37°C in a 5% CO_2_ incubator. Human islets were isolated from human cadaveric islets donors provided by the NIH/NIDDK-supported Integrated Islet Distribution Program (IIDP) (https://iidp.coh.org/overview.aspx), and from Prodo Labs (https://prodolabs.com/), the University of Miami, the University of Minnesota, the University of Wisconsin, the Southern California Islet Cell Resource enter, and the University of Edmonton, as summarized in Supplementary Table 5 Informed consent was obtained by the Organ Procurement Organization (OPO), and all donor information was de-identified in accord with Institutional Review Board procedures at The Icahn School of Medicine at Mount Sinai (ISMMS). Human islets were cultured and dispersed as previously described ^8^.

### Adenovirus

Human islets were transduced with RIP-ZsGreen at MOI of 100. Overnight, the following day, islets were dispersed, and seeded in RPMI with 100U/mL penicillin, 100 mg/mL streptomycin in the presence of an additional 100 MOI of the virus for 2-4 h. Following, FBS was added to a final concentration of 10%, compounds added and cells were continuously imaged for the course of 48-72 h.

### Immunostaining

INS-1 cells or dispersed human islet cells were plated on 12-mm laminin-coated glass coverslips. Cells were treated with the indicated conditions and times. Cells were washed with PBS and fixed in 4% paraformaldehyde. Immunolabeling was performed as previously described^25^ with primary antibodies directed against N-terminus ChREBP (1:250, ^25^), C-terminus ChREBP (Novus, 1:250), Pdx1 (Abcam, 1:500), and Insulin (Fisher, 1:1000)

### Glucose stimulated Insulin Secretion (GSIS) analysis

20 IEQ primary human were placed in 12 µm pore diameter inserts (Millipore) and incubated for 24 h with indicated treatment. Following, media was replaced with KREBS containing 2.8 mM glucose and 1% BSA and GSIS analysis was performed as previously described ^48^. Insulin levels were measured using a commercial ELISA assay (Mercodia), following the manufacturer instructions.

### qPCR and PCR

mRNA was isolated using the Qiagen RNAeasy mini kit for INS-1 cells, or for islets using the Qiagen RNAeasy mini kit. cDNA was produced using the Promega m-MLV reverse transcriptase. qPCR was performed on the QuantStudio5 using Syber-Green (BioRad) and analysis was performed using the ΔΔCt method. PCR for genotyping was performed using standard methods. Primer sequences are shown in Table S6.

### Proximity Ligation Assay (PLA)

PLA was used to determine endogenous protein–protein interactions as previously described in^86^. Briefly, ChREBPα (by GenScript) or NFATC1 (cell signaling) and Pan-14-3-3 (Santa Cruz) antibodies were conjugated to Duolink oligonucleotides, PLUS and MINUS oligo arms, respectively, using Duolink® In Situ Probemaker Following a PBS wash, cells were fixed with 4% formaldehyde solution for 10 min at room temperature, and blocked with Duolink Blocking Solution for 1 h at 37 °C and then incubated with 4 μg/mL ChREBPα-Plus and 14-3-3-MINUS overnight at 4 °C. PLA was performed according to the manufacturer’s directions. No secondary antibodies were used, because PLUS and MINUS oligo arms were directly conjugated to ChREBP and 14-3-3. Cells were imaged on a Zeiss 510 NLO/Meta system (Zeiss, Oberkochen, Germany), using a Plan-Apochromat 63×/1.40 oil differential interference contrast objective.

### Proliferation and cell death

Cellular proliferation and cell death was quantified using Real-Time Kinetic Labeling (SPARKL) technique^22^. Syto21 marked all cells, Yoyo3 marked all dead cells.

### Calcium Imaging

INS-1 cells seeded on glass bottom dish at low glucose. After 24 h media was replaced with KREBs buffer containing 5.5 mM glucose. Following 1 h incubation, Fluo-4 AM was added and incubated with cells for 45 min. The cells were then imaged at low glucose concentrations for 1 min, with images captured at 6 sec intervals. Glucose was supplemented to the cells to reach a final concentration of 20 mM and imaging was continued with the same settings for 5 additional min. The images were acquired using the confocal Cytation 10 system. Fluo-4 intensities were calculated using Gen5 Software.

### Active mitochondria staining

INS-1 cells were incubated with 250 nM MitoTracker® Orange CM-H2TMRos (Life Technologies, Eugene, OR) for 30 min at 37°C. Mitochondrial activity was analyzed based on fluorescence intensity using a confocal Cytation 10 system and Gen 5 software.

### ATP/ADP measurements

Measurements were performed using a commercially available ADP/ATP bioluminescent kit (Abcam).

### Luciferase Reporter

The reporter assays were done as described previously^17^. Briefly, INS-1-derived 832/13 cells that stably express the luciferase reporter gene under control of the human TXNIP promoter. Cells were harvested after 24 h and luciferase activity was measured using the Luciferase Reporter Assay System (Promega, Cat. #E1500) on Victor Nivo luminometer (PerkinElmer). Firefly luciferase activity was normalized to *Renilla* luciferase activity.

### Statistics

All studies were performed with a minimum of three independent replications. Data were represented in this study as means ± standard error of the mean (SEM). Statistical analysis was performed using two-way ANOVA on GraphPad (Prism) V9.2.

## Acknowledgments

This work was supported by the European Union through ERC Advanced Grant PPI-Glue (101098234), the Netherlands Ministry of Education, Culture and Science (Gravity program 024.001.035). The Netherlands Organization for Scientific Research (ECHO grant 711.018.003), and by DFG-funded CRC1093 (Supramolecular Chemistry on Proteins). Views and opinions expressed are however those of the authors only and do not necessarily reflect those of the European Union or the European Research Council. Neither the European Union nor the granting authority can be held responsible for them. DKS grant support from NIH/NIDDK R01DK130300 and the Human Islet and Adenoviral Core (HIAC) of P30DK020541.

We thank Michelle Arkin (UCSF) for stimulating discussions. We thank Sr. Sarah Stanley for help with Calcium imaging and members of the Diabetes Obesity and Metabolism Institute at Icahn School of Medicine at Mount Sinai for useful discussions. We acknowledge DESY (Hamburg, Germany), a member of the Helmholtz Association HGF, for the provision of experimental facilities. Parts of this research were carried out at PETRA III and we would like to thank Johanna Hakanpää for assistance in using beam P11. Beamtime was allocated for proposals 11011126 and 11010888. Further, we acknowledge the European Synchrotron Radiation Facility (ESRF) for provision of synchrotron radiation facilities and thank Shibom Basu for assistance and support in using beamline ID23-2 (mx2407).

## List of Abbreviations

ChREBP: carbohydrate response element binding protein
ChoRE: carbohydrate response element
TF: transcription factor
PPI: protein-protein interaction
T2D: type 2 diabetes
AMP: adenosine monophosphate
LID: low glucose inhibitory domain
NES: nuclear export signal
SAR: structure-activity relationship
α: cooperativity factor
FA: fluorescence anisotropy
Kd^II^: intrinsic compound affinity
SPARKL: single-cell and population-level analyses using real-time kinetic labeling
RIP: rat insulin promoter
GSIS: glucose-stimulated gene expression
SI: stimulation index
PLA: proximity ligation assay

