## Supplemental figures, tables and information for "Molecular glues of the regulatory ChREBP/14-3-3 complex protect beta cells from glucolipotoxicity"

### Supporting Information

### SUPPLEMENTAL FIGURES

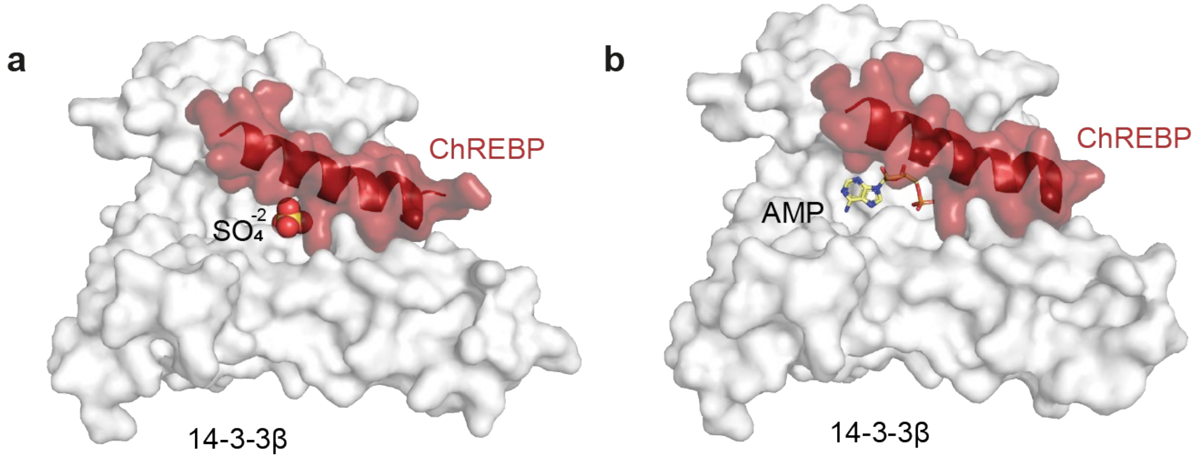


**Fig. S1 | Molecular basis of the PPI between 14-3-3 and ChREBP.** Crystal structures of 14-3-3β (white) in complex with ChREBPα (red) and (**a**) $SO_4^{2-}$ (PDB ID: 4GNT), or (**b**) AMP (PDB ID: 5F74).

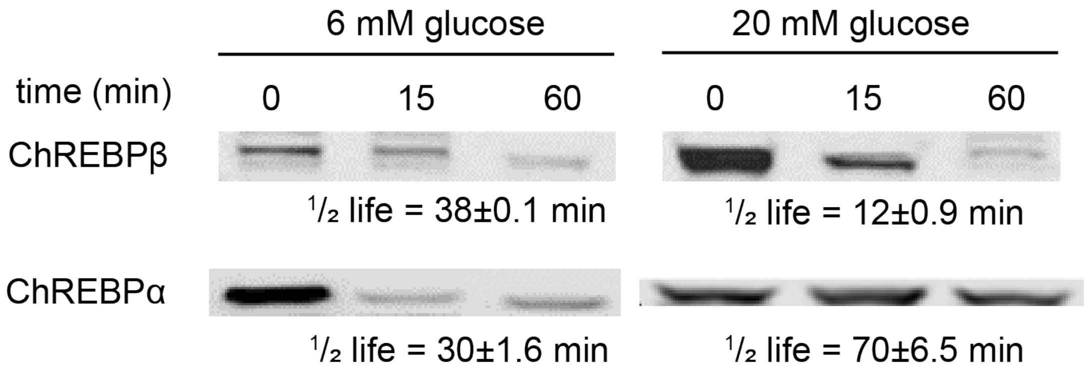


**Fig. S2 | Differential stability of ChREBPα and ChREBPβ depending on glucose exposure.** Endogenously GFP-tagged ChREBP INS-1 cells (where both ChREBPα and ChREBPβ are tagged with GFP) were cultured in 6 mM or 20 mM glucose for 2 days, and then treated with 100 µg/ml cycloheximide (CHX) and either 6 mM or 20 mM glucose for the indicated times. ChREBPα and ChREBPβ were visualized in immunoblots using GFP antibody. Half-life was calculated from densitometry as a percentage of remaining signals. Representative blots from 3 experiments.

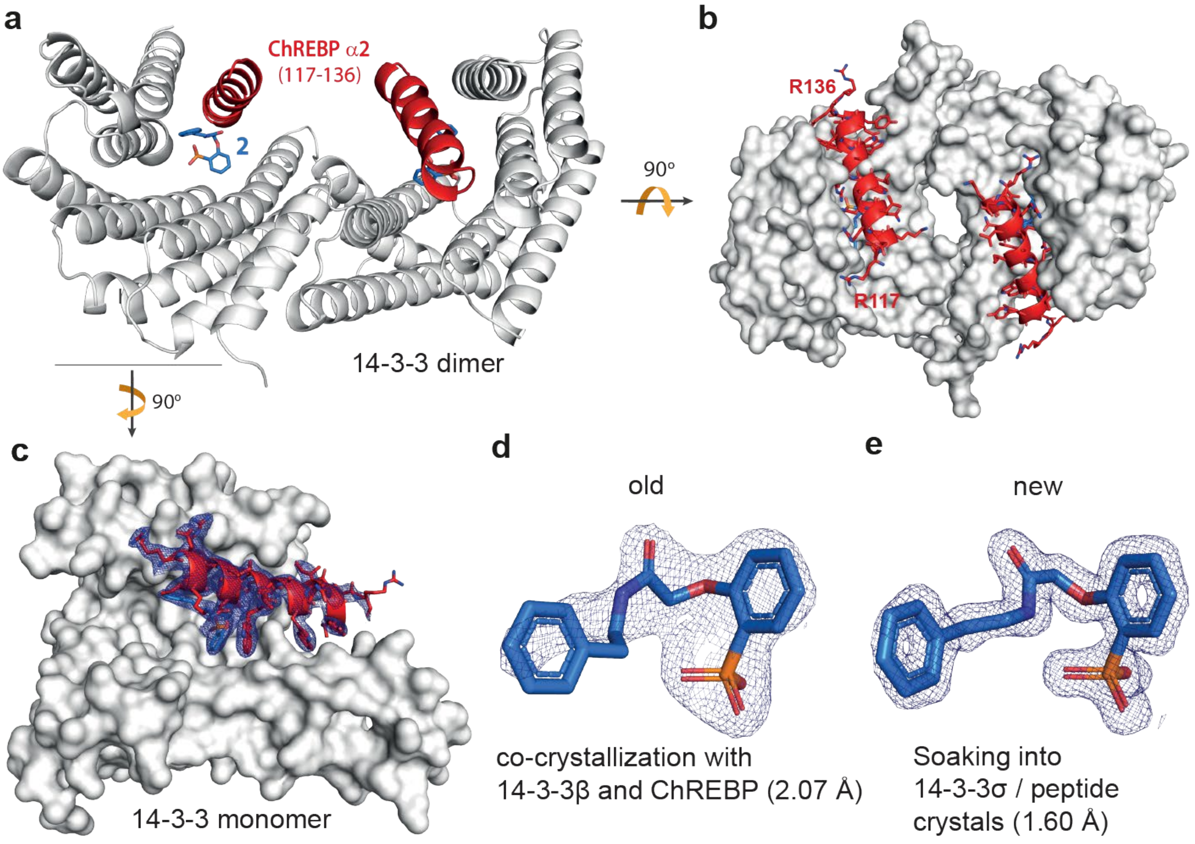


**Fig. S3 | Co-crystal structure of 14-3-3β in complex with ChREBPα and stabilizer. a**. Side view of the 14-3-3β dimer (white cartoon) bound by two ChREBPα peptides (red cartoon) and two **1** molecules (blue sticks). **b**. Top view displaying the antiparallel orientation of the ChREBP helices in the 14-3-3 dimer. **c**. Front view of one 14-3-3β monomer (white surface) bound by one ChREBP peptide (red cartoon) and **1** (blue sticks). **d.** Close-up of the electron density of **1** solved by the co-crystallization with 14-3-3β and ChREBP (PDB ID: 6YGJ, 'old' structure). **e.** Close-up of electron density of **1** solved by the soaking method into 14-3-3σ crystals ('new' structure).

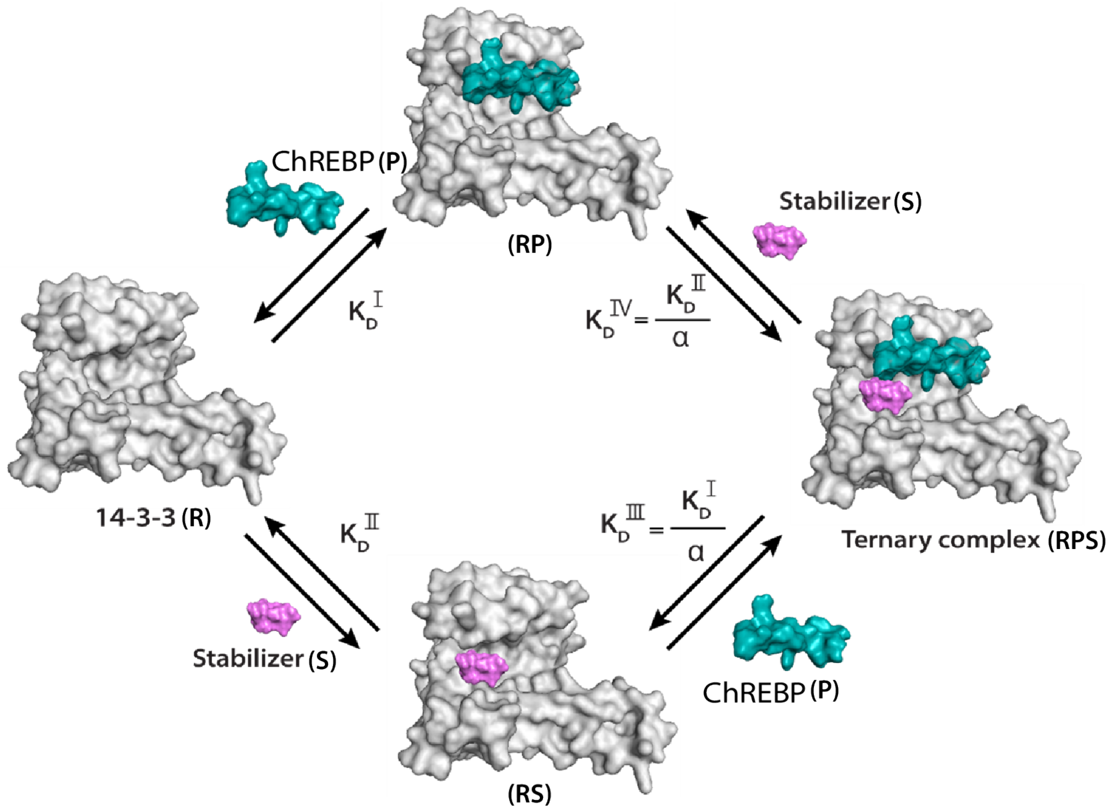


Equilibrium equations:

$KD^{I} = [R]*[P]/[RP]$

$KD^{II} = [R]*[S]/[RS]$

$KD^{III} = [RS]*[P]/[RPS]$

$KD^{IV} = [RP]*[S]/[RPS]$

The cooperativity factor α is defined as:

$\alpha = KD^{I}/KD^{III} = KD^{II}/KD^{IV}$

**Fig. S4 | Cooperativity scheme of 14-3-3/ChREBP PPI stabilization.** The ChREBP peptide (P) binds to the 14-3-3 protein (R) with $KD^{I}$ and in the presence of the stabilizer molecule (S) this affinity is altered to $KD^{I}/\alpha$. Similarly, the stabilizer binds to 14-3-3 with an intrinsic affinity $KD^{II}$ and an enhanced affinity $KD^{II}/\alpha$ when ChREBP is already bound to 14-3-3. Equilibrium equations that are used in the thermodynamic equilibrium model.

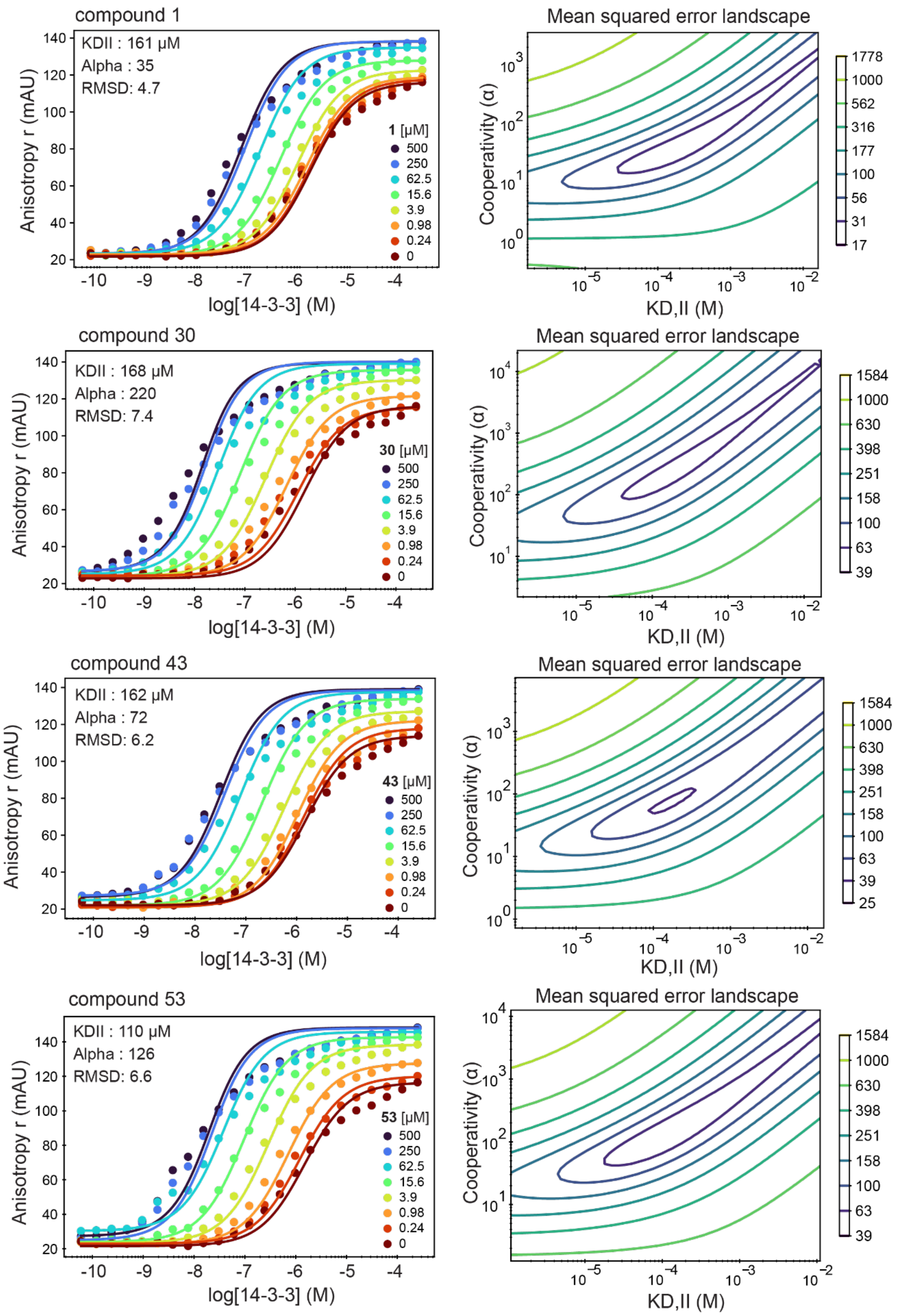


**Fig. S5 | Cooperativity analysis.** Left: experimental fluorescence anisotropy data of 14-3-3 titration to FITC-labeled ChREBPα peptide in the presence of several concentrations of compound **1**, **30**, **43**, **53**. Right: error-landscape plot centered on the determined $K_D^{II}$ and α factors.

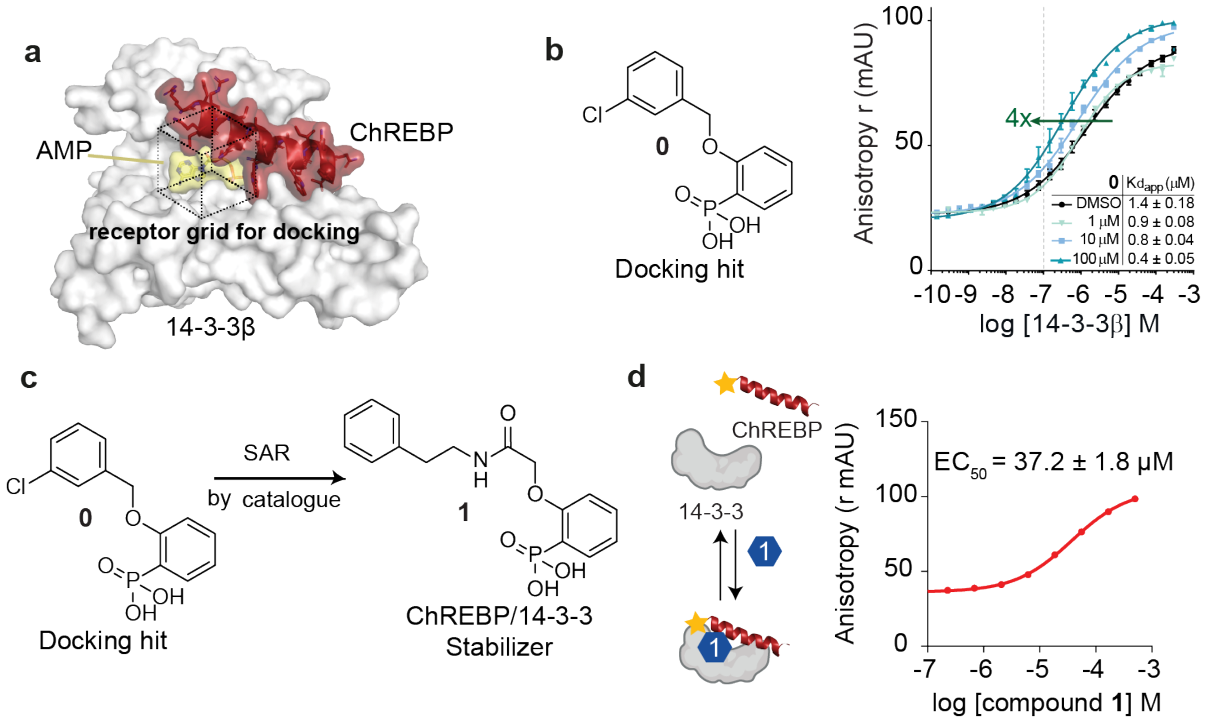


**Fig. S6 | Schema for SAR. a.** The receptor grid (black dotted box) for docking in the crystal structure of 14-3-3β (white surface), ChREBP (red cartoon and sticks) and AMP (yellow surface) (PDB entry 5f74). **b.** Molecular structure of docking hit **0** and fluorescence anisotropy (FA) data of 14-3-3β titration to FAM-labeled ChREBP peptide (100 nM) and fixed concentrations (1, 10 and 100 μM) of compound **0**, showing a 4-fold increase in stabilization with 100 μM. **c**. Molecular structure of ChREBP/14-3-3 stabilizer **1** discovered by SAR by catalogue from starting point **0**. **d.** FA compound titration of **1** into FITC-labeled ChREBP (10 nM), and 14-3-3β (150 nM).

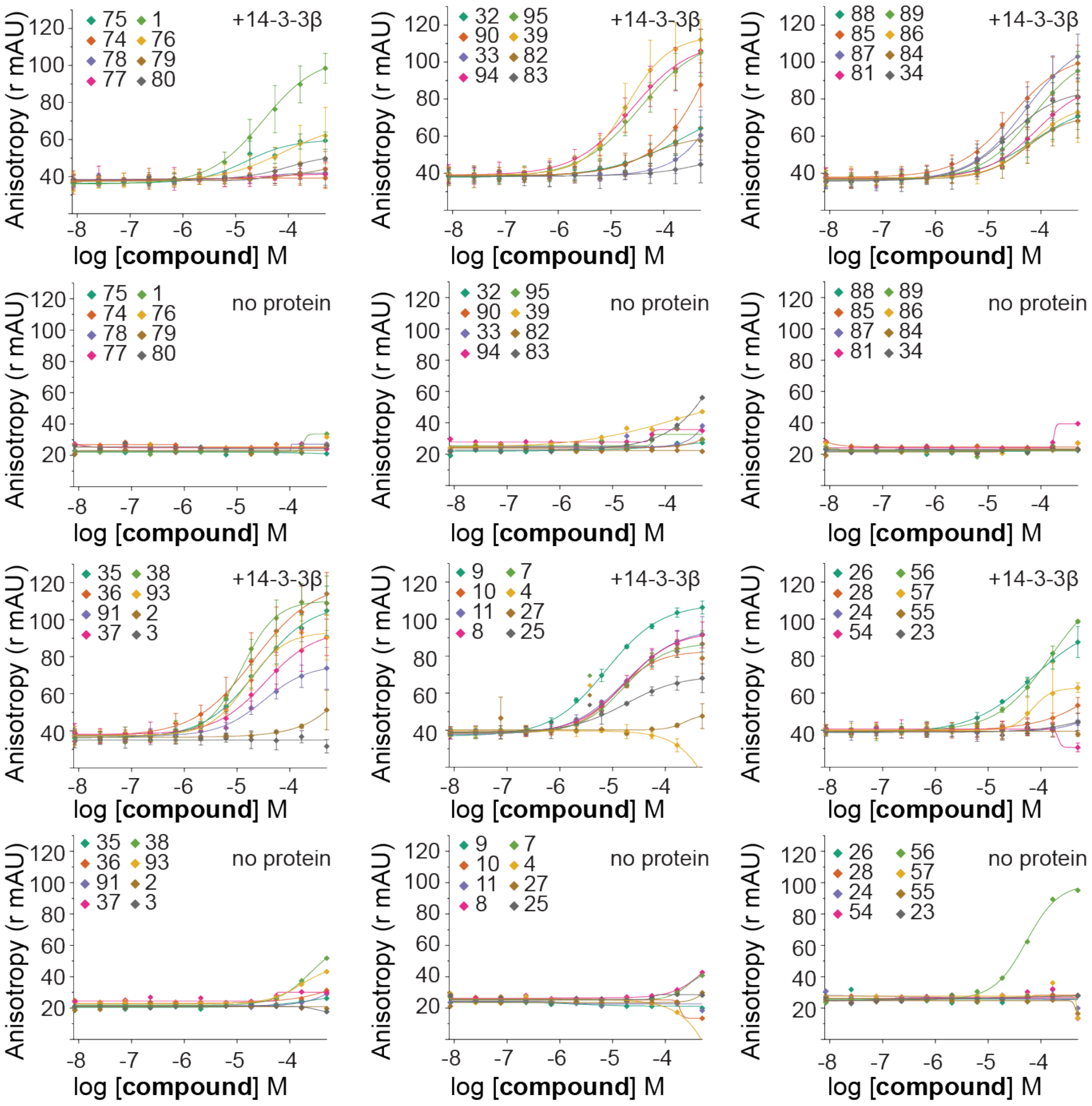


**Fig. S7 | Fluorescence Anisotropy (FA) compound titrations described in Table S1-2-3 (part I).** Compounds were titrated in FITC-labeled ChREBP peptide (10 nM) with or without the presence of 14-3-3β (150 nM).

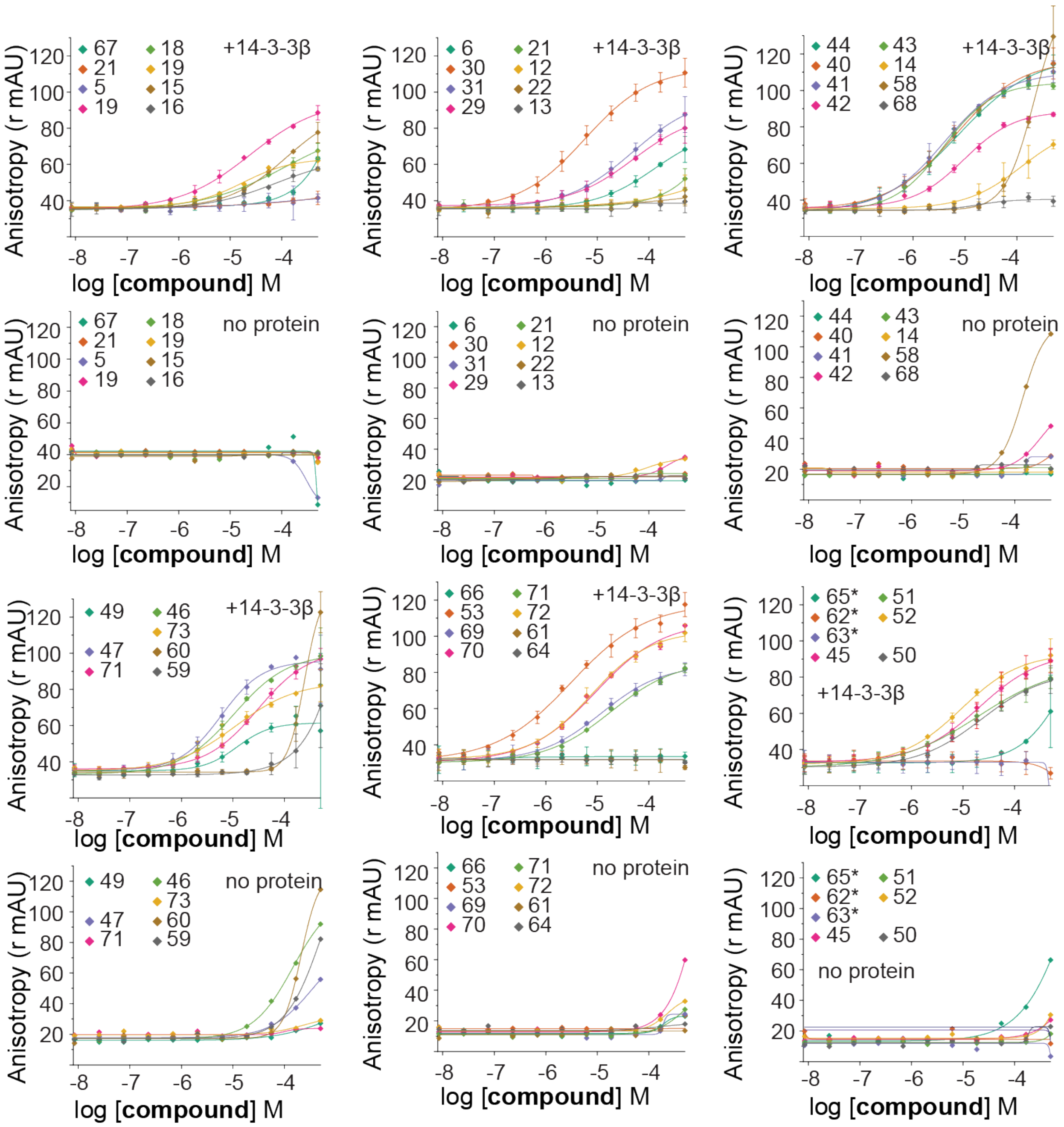


**Fig. S8 Fluorescence Anisotropy (FA) compound titrations described in Table S1-2-3 (part II).** Compounds were titrated in FITC-labeled ChREBP peptide (10 nM) with or without the presence of 14-3-3β (150 nM).

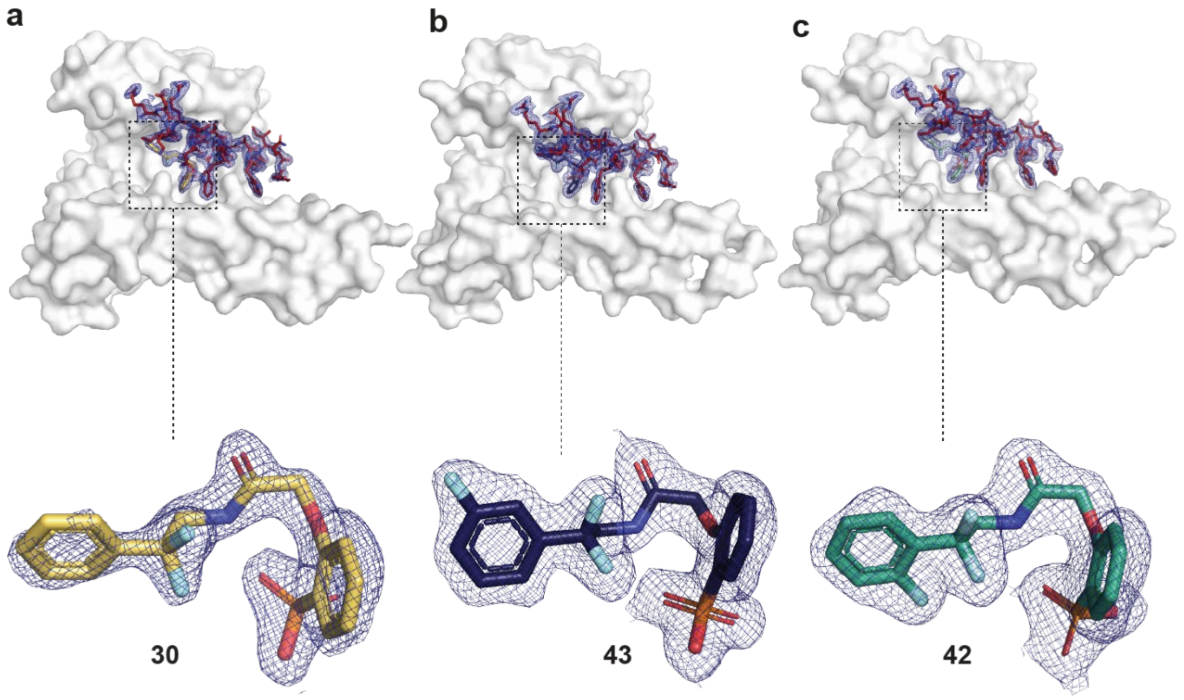


**Fig. S9 | Crystal structures** of 14-3-3σ (white surface) in complex with ChREBP (red sticks) and stabilizer. **a. 30** as yellow sticks. **b. 43** as purple sticks. **c. 42** as green sticks. Fo-Fc electron density maps (blue mesh) are contoured at 1σ.

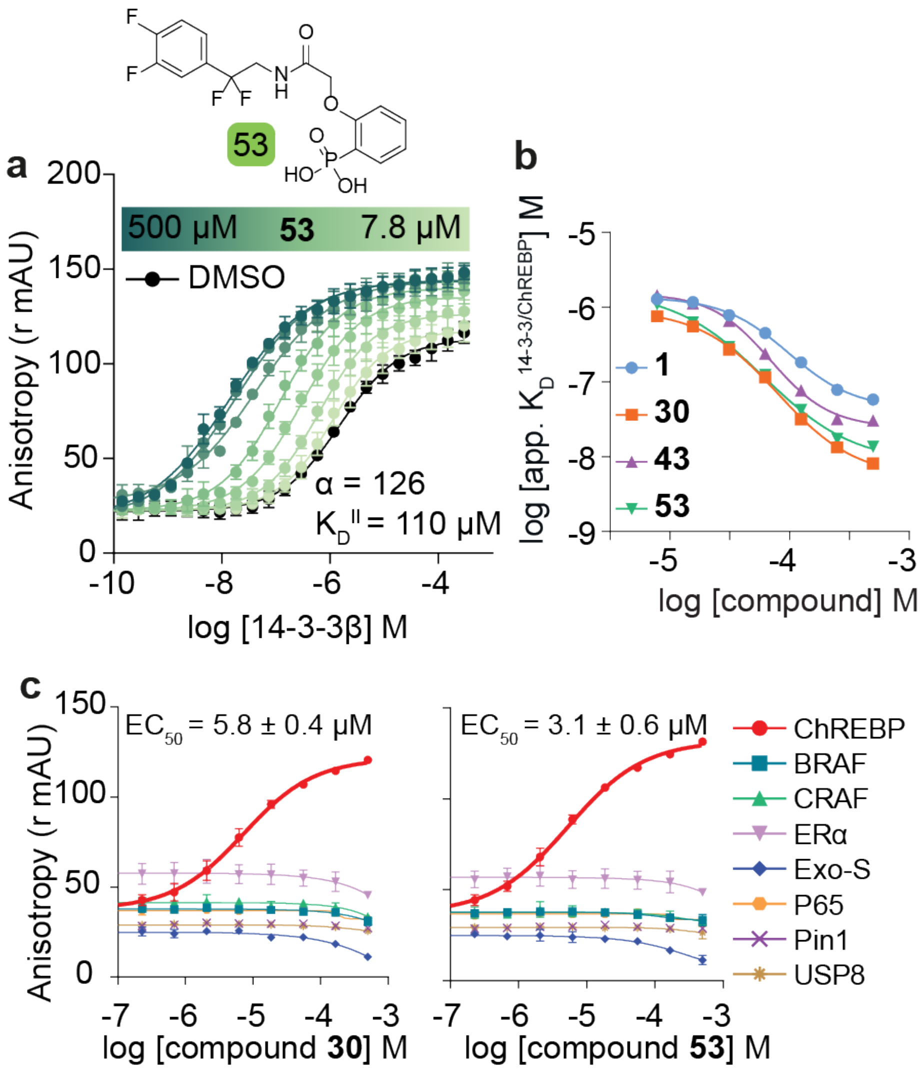


**Fig. S10 | Affinity, cooperativity and selectivity measurements for the ChREBP/14-3-3 interaction in the presence of molecular glues. a.** Titration of 14-3-3β to FITC-labeled ChREBP peptide (10 nM) against varying fixed concentrations of **53** (0–500 μM) (mean ± SD, n = 2). **b.** Apparent $K_D$ value of the ChREBPα–14-3-3 interaction (y-axis) in the presence of a range of concentrations (0–250 μM) of **1** (blue), **30** (orange), **43** (purple) and **53** (green) (x-axis) **c.** Selectivity studies by titrating **30** or **53** to 14-3-3β and eight different 14-3-3 interaction FITC-labeled peptides (all 10 nM) (mean ± SD, n = 2).

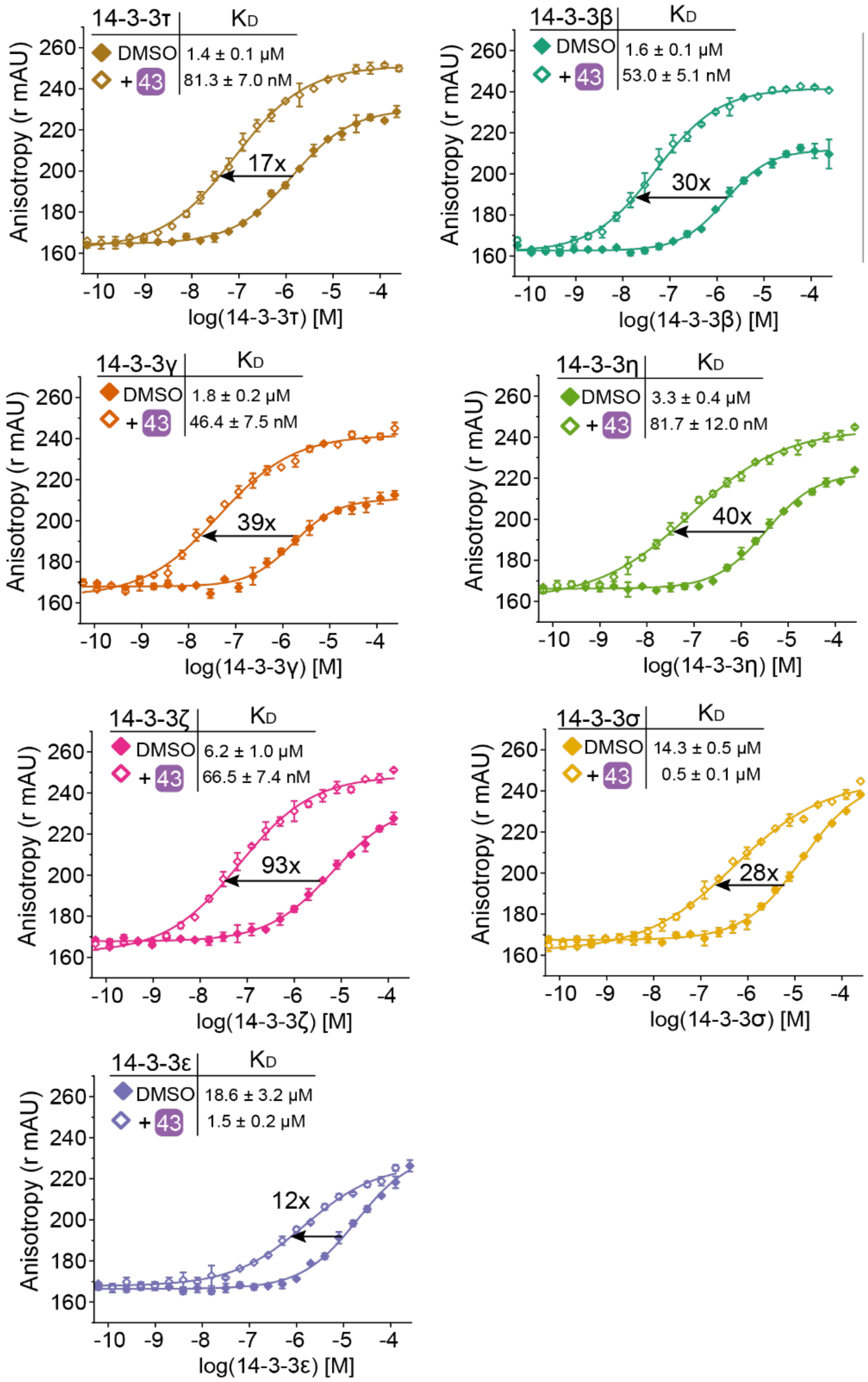


**Fig. S11 | Fluorescence Anisotropy (FA) protein titrations of 14-3-3 isoforms comparing DMSO and 43.** 14-3-3 isoforms were titrated to FITC-ChREBP (10 nM) with and without 100 μM of compound **43**. The stabilization factor ($K_D$ DMSO/$K_D$ +43) is shown above the arrow.

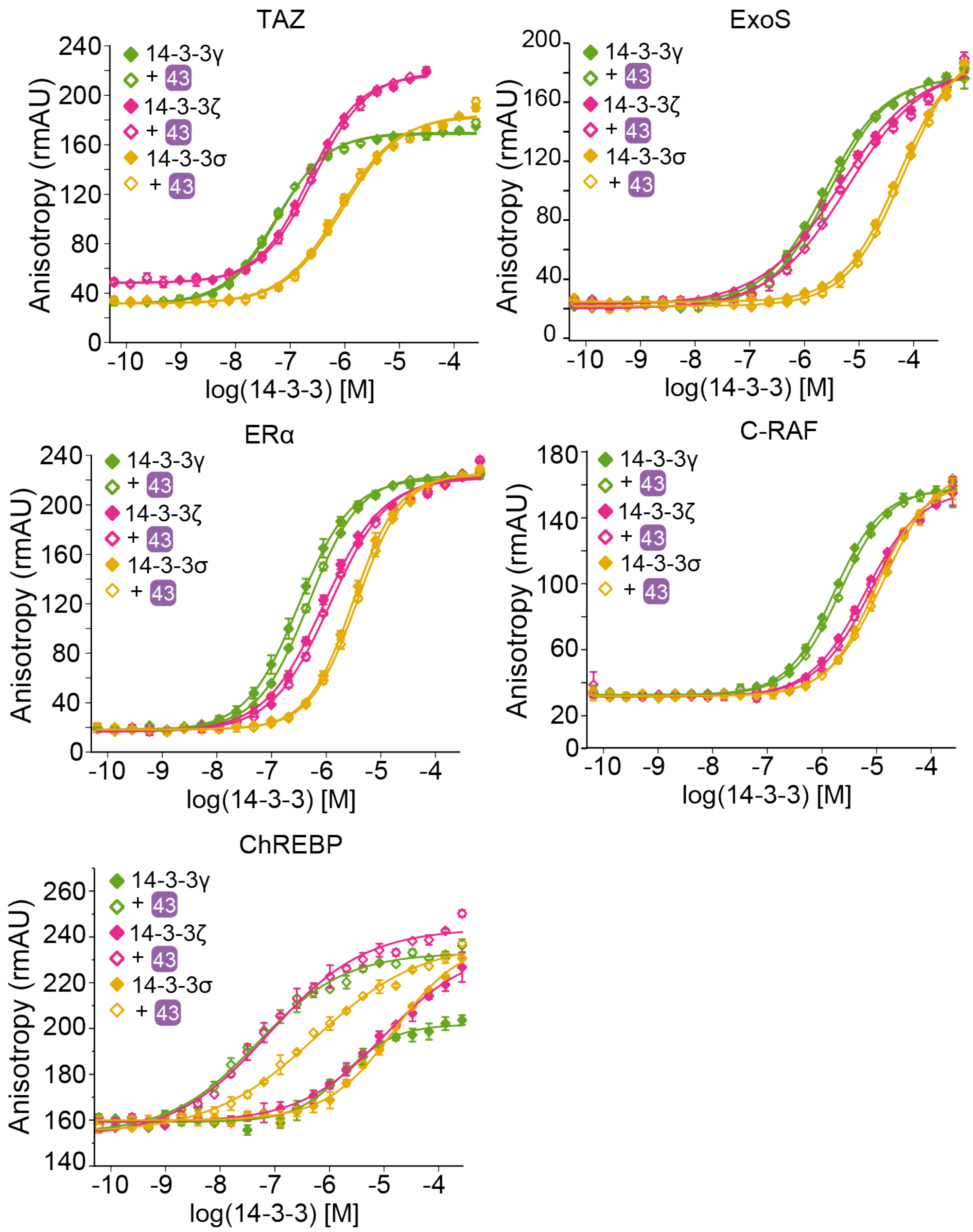


**Fig. S12 | Fluorescence Anisotropy (FA) measurements for studying the selectivity of compound 43 among different 14-3-3 isoforms (γ, ζ, σ).** Titration of the 14-3-3 isoforms to FITC-TAZ, FITC-ExoS, FITC-ERα, FITC-CRAF and FITC-ChREBP (10 nM) with and without 100 μM of compound **43** (mean ± SD, n = 2).

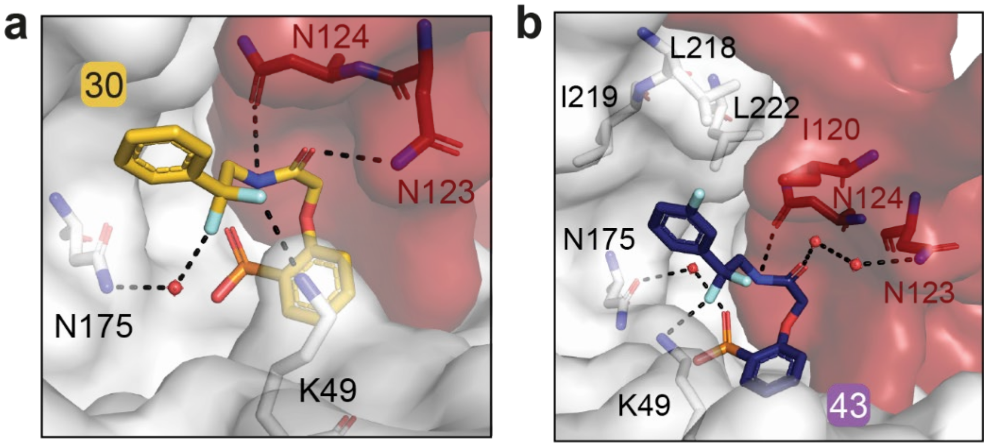


**Fig. S13 | Interactions of 30 and 43 at the ChREBPα/14-3-3 interface.** Structure of **30** (**c.** yellow) and **43** (**b.** purple) in complex with 14-3-3σ (white cartoon) and ChREBP (red cartoon), showing some conformations of K49 of 14-3-3 and N123 of ChREBP. This resulted in slightly different polar interactions with the stabilizers (black dashed lines).

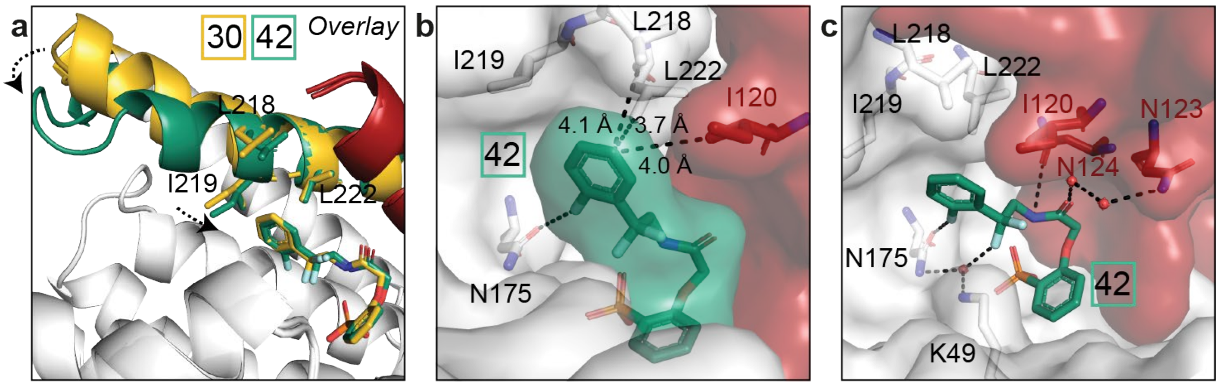


**Fig. S14 | Crystal structure of 42. a.** Crystallographic overlay of **30** (yellow) and **42** (green) in complex with 14-3-3σ (white cartoon) and ChREBP (red cartoon). The color of helix 9 of 14-3-3σ is matched to the corresponding small molecule bound, thus demonstrating the helical 'clamping' effect when **42** (green) is present. **b.** Surface representation of **42** (green) in complex with 14-3-3 (white) and ChREBP (red), showing the distances (black dashes) of the phenyl ring of **42** to the residues (sticks) of 14-3-3 and ChREBP. **c.** Interactions of **42** (green) with 14-3-3 (white) and ChREBP (red) (relevant side chains are displayed in stick representation, polar contacts are shown as black dashed lines).

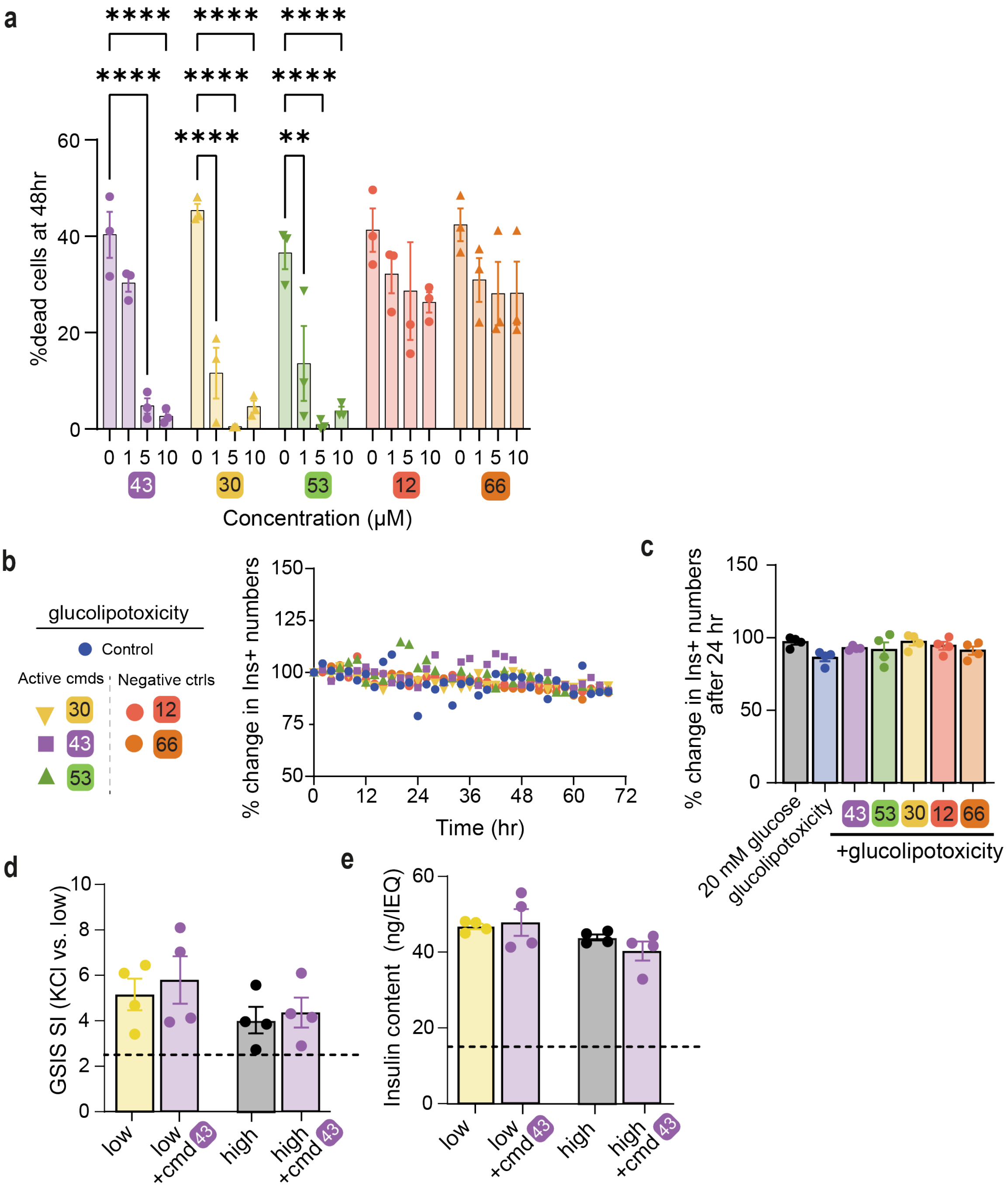


**Fig. S15 | Active compounds are effective at protecting INS-1 cells from glucolipotoxicity**. **a.** Dose response of active stabilizers: INS-1 cells were cultured in glucolipotoxic conditions (20 mM glucose+500 μM palmitate) for 48 hours, in the presence of control compounds (red) or active stabilizers (green) at indicated concentrations. Data is % Yoyo+ cells at 48 hr. Data are the means ± SEM, $n = 3$, **, $p < 0.01$; ****, $p < 0001$. **b.** Representative kinetic measurement of percent change in insulin positive (INS+) cell numbers over time. **c.** Overall change in β-cell numbers at 24 h. Data are means +/- SEM; n=4; *, $p < 0.05$ compared to glucolipotoxicity. **d.** Human islets were treated for 24 h as indicated, then GSIS was measured in KREBS buffer (2.8 mM glucose, 1% BSA) over 30 min. Following, 30 mM KCl was added to the islets for 30 min. Stimulation index (SI) is the amount of insulin released at high vs low glucose. **e.** Total insulin content in primary islets treated as in (**d**).

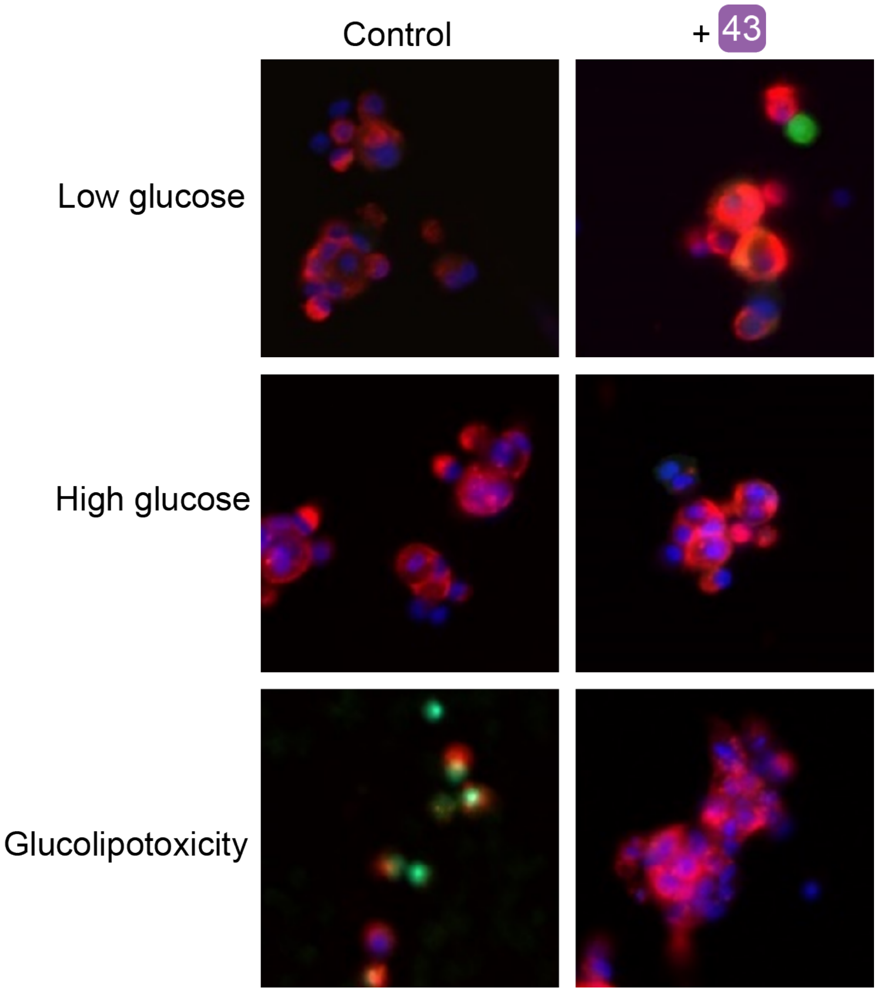


**Fig. S16 | Compound 43 rescues human islets from glucolipotoxicity.** Dispersed Human islets were cultured in low (5mM), high (20mM) or glucolipotoxic conditions (20 mM glucose+500 μM palmitate) in the presence or absence of 10 μM 43 for 24 h. Following, cells were fixed and stained for TUNEL (green) and immunostained for insulin (red), or Dapi (blue). Representative figures from n=2 human islet donors.

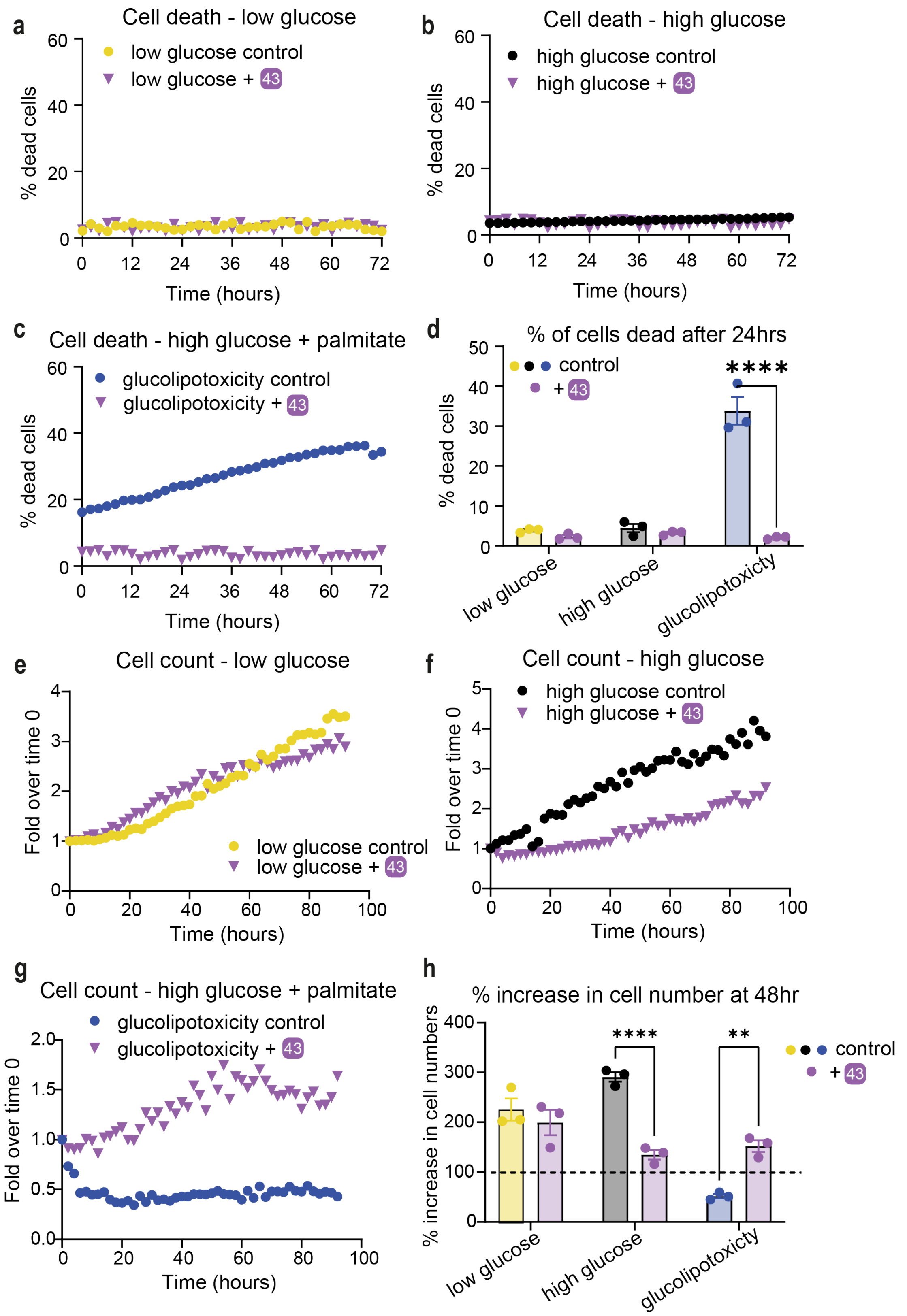


**Fig. S17 | Kinetics of cell proliferation and death in INS-1 cells.** INS-1 cells were stained with Syto21 to mark all live cells and Yoyo3 to mark dead cells. Cells were continuously monitored over 72 h and imaged every 2 h. (**a-d**) Dead cell quantifications (**e-h**) Total cell numbers quantification at Low (5 mM **a**, **e**, **d**, **h**)), High (20mM, **b**, **f**, **d**, **h**) glucose concentrations or glucoliptoxicity (20 mM glucose+500 μM palmitate, **c**, **g**, **d**, **h**). w/o 10 μM compound **43**.

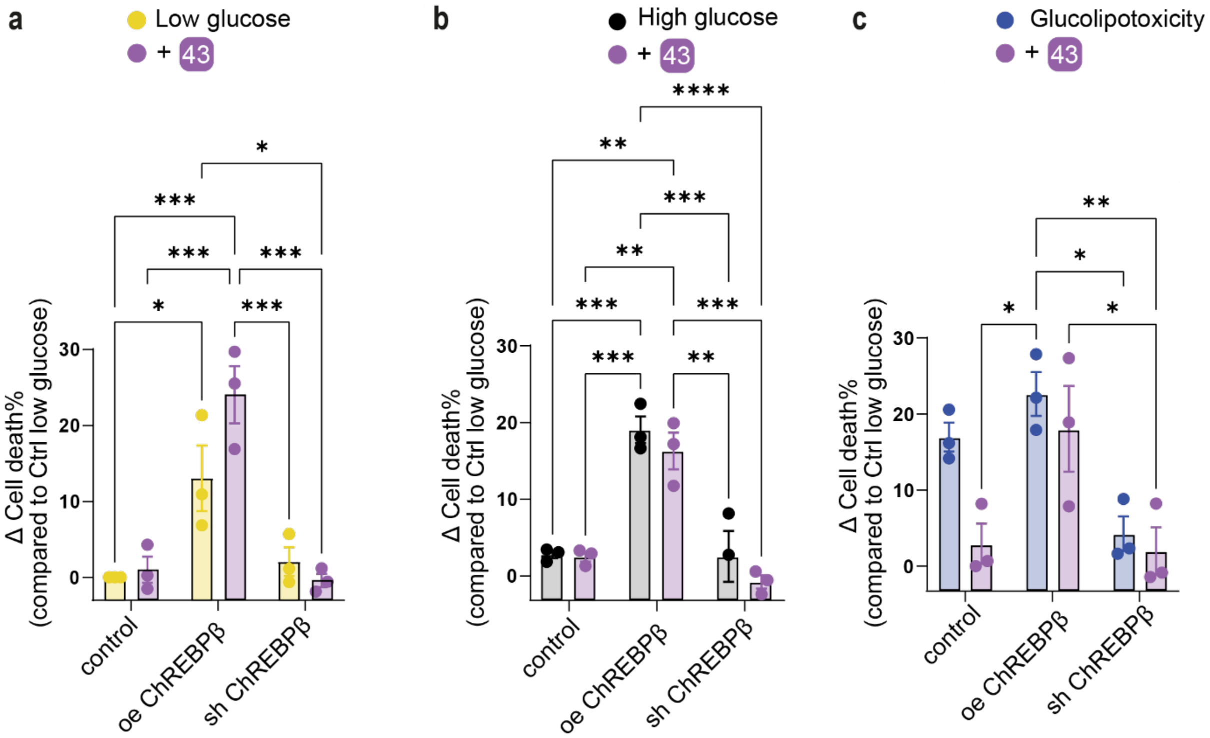


**Fig S18 | 43 activity is mediated by blocking the induction of ChREBPβ.** Dispersed human islet cells were transduced for 48 h with the indicated adenovirus (oe, overexpressed; sh, shRNA) and the RIP-zsGreen adenovirus to mark β-cells. Following, they were exposed to Low (5.5 mM) glucose (a), High (20 mM) glucose (b) or glucolipotoxicity (20 mM glucose and 0.5 mM palmitate, c) in the presence of 10 µM **43** and Yoyo3 to mark for dead cells. Cells were imaged by the Cytation 10 and data analyzed by Gen5. Presented here is the increase in β cell death 24 h following the indicated treatment. Data are means +/- SEM; n=3; *, P<0.5; **, P<0.01; ***, P<0.005; ****, P<0.001.

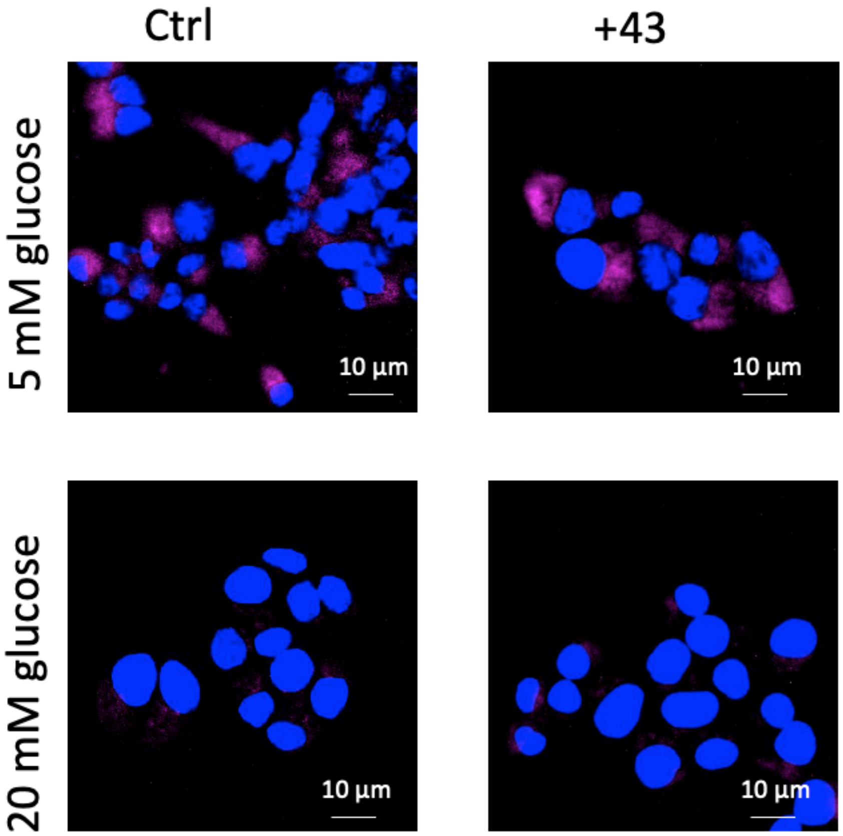


**Fig. S19 | Proximity ligation assay demonstrating 43 does not affect interaction or dissociation between 14-3-3 and NFATC1.** INS-1 cells were cultured overnight (ON) at 5.5 mM glucose and exposed to high (20 mM) glucose for the indicated times in the presence or absence of 10 µM **43.** Cells were fixed at 2 h following indicated treatment.

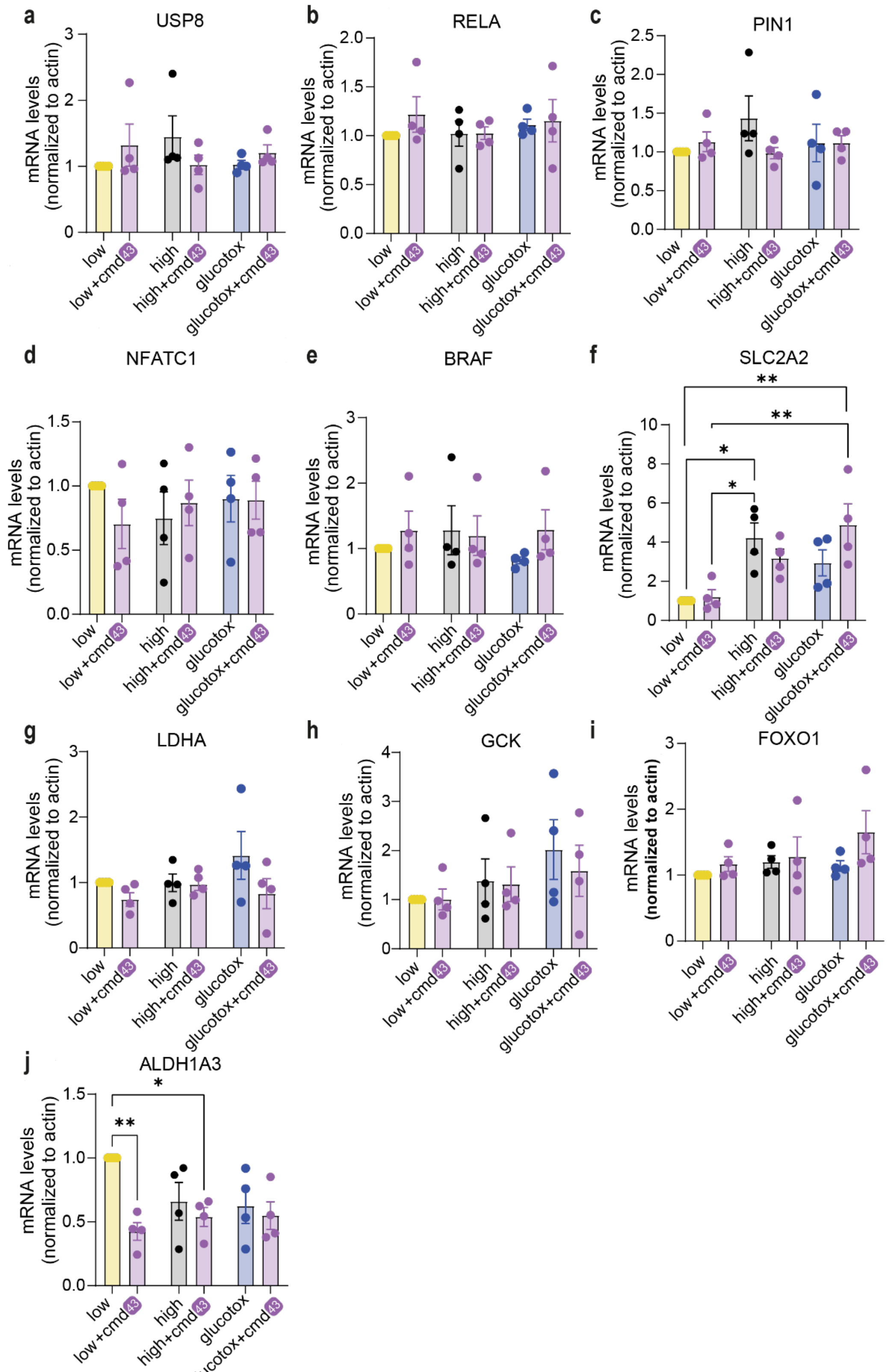


**Fig. S20 | Expression levels of 14-3-3 client protein genes, β-cell identity genes and disallowed genes.** mRNA fold-enrichment over control low glucose in human islets, treated with 10 μM **43** for 24 h; Low-5.5 mM glucose, High-20 mM glucose, glucotox-20 mM glucose+500 μM palmitate. Data are the means +/- SEM, n=4, *p<0.05, **p<0.01***, p<0.005, ****, p<0.001.

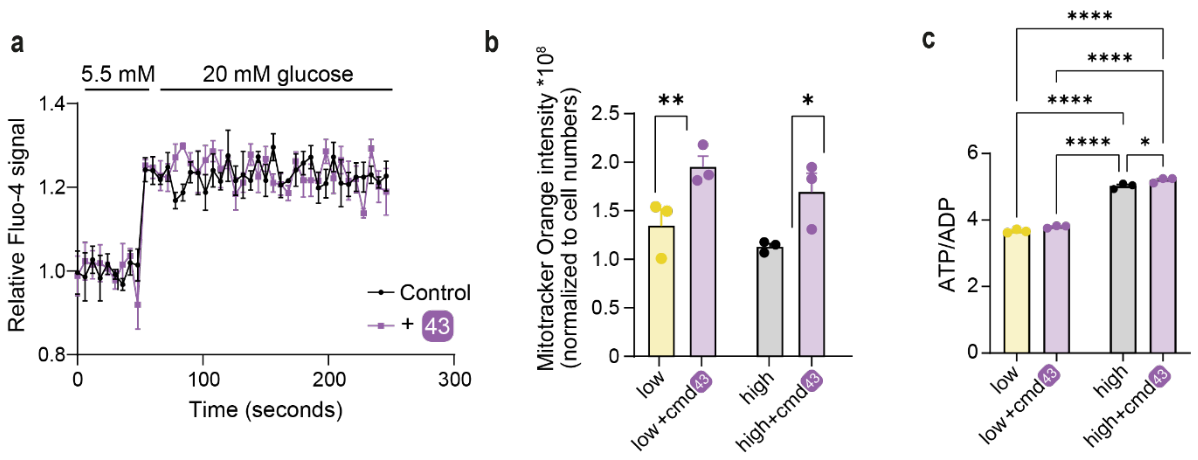


**Fig. S21 | 43 does not affect Ca$^{++}$ influx or mitochondrial activity. a.** Calcium imaging in INS-1 cells at indicated conditions. **b.** Active mitochondira in INS-1 cells using Mitotracker-Orange CM-H2TMRos. **c.** ATP/ADP ratio in INS-1 cells at low (5.5 mM) or high (20 mM) glucose w/o 10 µM 43. Data are means +/- SEM; n=3; *, P<0.5; **, P<0.01; ***, P<0.005; ****, P<0.001.

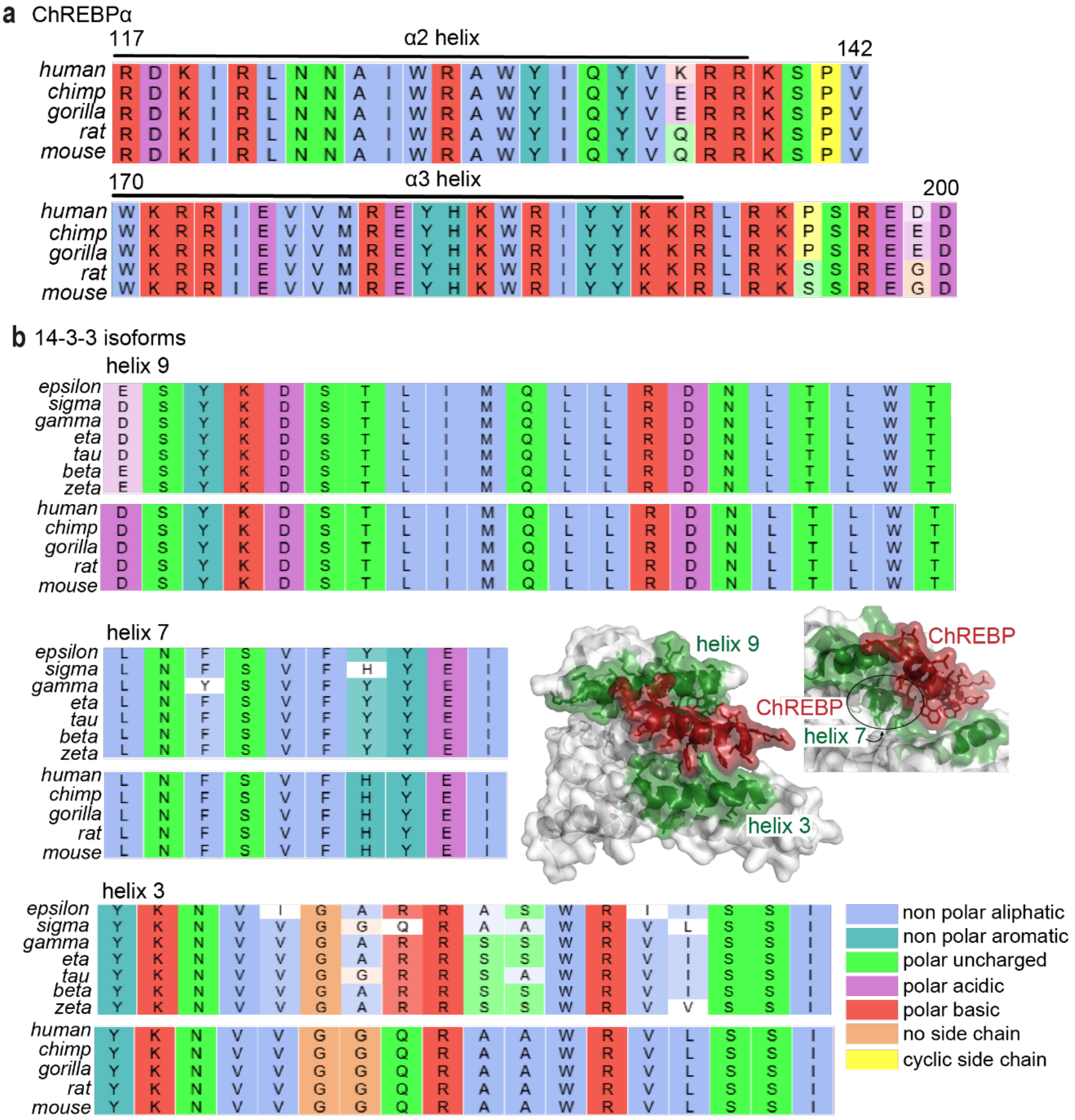


**Fig. S22 | Sequence alignment of interacting regions of 14-3-3 and ChREBPα. a.** Sequence alignment of the α2 and α3 helices of ChREBPα among species: human, chimpanzee, gorilla, rat, mouse. **b.** Sequence alignment of (part of) helix 9, 7 and 3 (green in crystal structure) of 14-3-3 (grey) which interact with the α2 helix of ChREBP (red). The first panel shows the sequence conservation among the seven human 14-3-3 isoforms. The second panel shows the alignment for the same helices among the species; human, chimpanzee, gorilla, rat and mouse, for one of the 14-3-3 isoforms (sigma).

### SUPPLEMENTAL TABLES

**Table S1 | Structure and activity of analogs of 1.** Compound **1** marked blue, and the two best stabilizers marked yellow. $EC_{50}$ in parenthesis with mean ± SD, n = 2. For FA titration graphs see **Fig. S4, S5**. Compounds with a star (*) were also described previously[12]. However, compared to the previously described compound titrations, the assay was adapted and performed at lower 14-3-3β concentrations to enlarge the window of detection of the improved stabilizers. Therefore, the reported $EC_{50}$ values are different compared to[12].

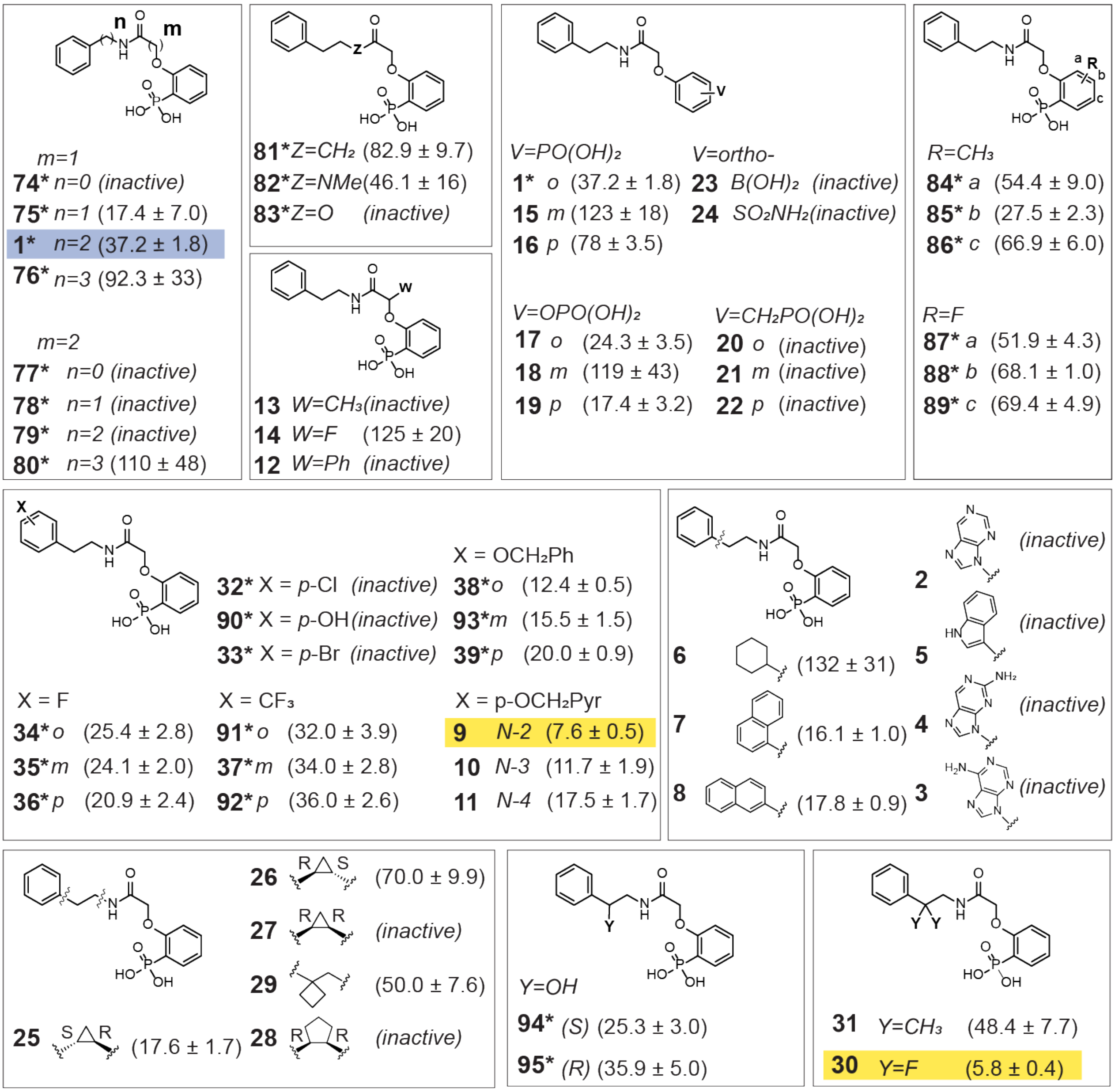


In parenthesis $EC_{50}$ (μM)
*described in *Nat. comm. (2020)*

**Table S2 | Structure and activity of sulfonamide and boric acid analogs.** $EC_{50}$ in parenthesis with mean ± SD, n = 2. For FA titration graphs see **Fig. S4, S5**.

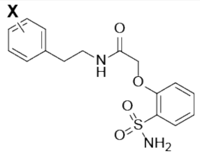

X=

| | | |
|---|---|---|
| **54** | *p*-F | (*inactive*) |
| **55** | *o*-$OCH_2Ph$ | (*inactive*) |
| **56** | *p*-$OCH_2Ph$ | (*inactive*) |
| **57** | *p*-$OCH_2Pyr$ (N-2) | (68.2 ± 7.9) |

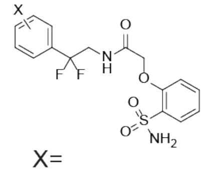

X=

| | | |
|---|---|---|
| **58** | H | (*inactive*) |
| **59** | *m*-F | (*inactive*) |
| **60** | *p*-F | (*inactive*) |
| **61** | *m*-$CF_3$ | (*inactive*) |
| **62** | *m*-$OCF_3$ | (*inactive*) |

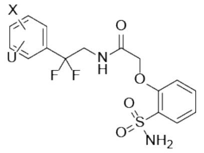

X=F, U=F

| | | |
|---|---|---|
| **63** | *o,o* | (*inactive*) |
| **64** | *o,m* | (*inactive*) |
| **65** | *m,m* | (*inactive*) |
| **66** | *m,p* | (*inactive*) |

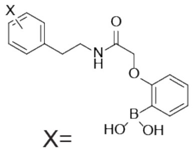

X=

**67** *p*-F (*inactive*)

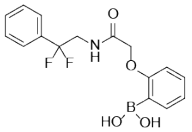

**68** (*inactive*)

**Table S3 | Structure and activity of fluorinated analogs described in Figure 2A.** $EC_{50}$ in parenthesis with mean ± SD, n = 2. For FA titration graphs see **Fig. S4, S5**. Compound numbers indicated with a star (*) are not visualized in **Fig. 3A**.

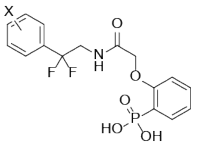

**30** X = H (5.7 ± 0.4)

| | | |
|---|---|---|
| X = Cl | | |
| **69*** | *o* | (12.0 ± 1.3) |
| **70*** | *m* | (9.3 ± 1.9) |
| **40** | *p* | (6.6 ± 0.9) |
| X = Br | | |
| **71*** | *o* | (18.5 ± 3.4) |
| **72*** | *m* | (7.6 ± 0.9) |
| **41** | *p* | (4.1 ± 0.4) |
| X = F | | |
| **42** | *o* | (9.6 ± 1.1) |
| **43** | *m* | (3.8 ± 0.2) |
| **44** | *p* | (7.5 ± 1.1) |
| X = $CF_3$ | | |
| **45** | *m* | (20.8 ± 2.3) |
| X = $OCH_2Ph$ | | |
| **46** | *o* | (10.6 ± 1.0) |
| **47** | *p* | (5.9 ± 0.7) |
| *o*-$OCH_2Pyr$ (N-2) | | |
| **73*** | | (11.0 ± 1.5) |

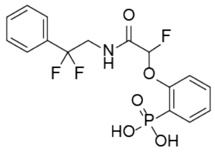

**48** (30.3 ± 1.3)

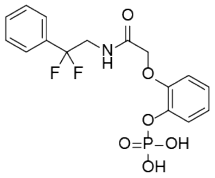

**49** (12.1 ± 3.4)

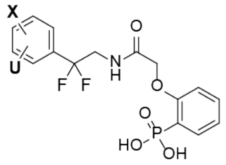

X=F U=F

| | | |
|---|---|---|
| **50** | *o,o* | (22.7 ± 4.2) |
| **51** | *o,m* | (20.0 ± 3.2) |
| **52** | *m,m* | (9.3 ± 1.1) |
| **53** | *m,p* | (3.1 ± 0.6) |

**Table S4 | XRD data collection and refinement statistics for ChREBPα–14-3-3σ structures.**

| 14-3-3σΔC/ ChREBP | 1 | 30 | 43 | 42 |
|---|---|---|---|---|
| PDB ID | **8BTQ** | **8C1Y** | **8BWE** | **8BWH** |
| **Data collection** | | | | |
| **Collection source** | DESY Petra III | DESY Petra III | ESRF ID23-2 | ESRF ID23-2 |
| **Collection date** | 04-11-2020 | 09-08-2020 | 29-01-2021 | 29-01-2021 |
| **Wavelength (Å)** | 1.033220 | 1.033220 | 8.873128 | 0.873128 |
| **Resolution (Å)[a]** | 1.60 – 45.36 (1.60 – 1.63) | 1.80 – 45.47 (1.80 – 1.84) | 2.00 – 46.41 (2.00 – 2.05) | 2.10 – 34.64 (2.10 – 2.16) |
| **Space group** | C 2 2 2 1 | C 2 2 2 1 | C 2 2 2 | C 2 2 2 |
| **Unit cell** | 81.979 111.932 62.339 | 81.626 111.651 62.827 | 63.316 147.471 76.968 | 63.166 147.336 77.070 |
| **Total reflections[a]** | 493511 (21933) | 351860 (22348) | 89504 (6796) | 88577 (7393) |
| **Unique reflections[a]** | 38129 (1837) | 26659 (1571) | 20127 (1550) | 20407 (1695) |
| **Redundancy[a]** | 12.9 (11.9) | 13.2 (14.2) | 4.4 (4.4) | 4.3 (4.4) |
| **Completeness (%)[a]** | 99.9 (99.6) | 98.9 (98.6) | 82.2 (85.8) | 96.7 (99.2) |
| **Average I/σ(I) [a]** | 26.0 (5.7) | 18.8 (1.5) | 9.0 (1.5) | 7.2 (1.6) |
| **Wilson B-factor (Å²)** | 17.20 | 30.70 | 24.54 | 28.57 |
| **$CC_{1/2}$[a,b]** | 0.999 (0.959) | 0.999 (0.635) | 0.992 (0.500) | 0.985 (0.613) |
| **$R_{merge}$[a,c,e]** | 0.066 (0.475) | 0.085 (1.854) | 0.127 (0.936) | 0.114 (0.717) |
| **$R_{meas}$[a,d,e]** | 0.071 (0.519) | 0.094 (1.994) | 0.154 (1.113) | 0.145 (0.894) |
| **Refinement** | | | | |
| **Reflections in set: Refinement / R-free[a]** | 38106 (3734) / 1856 (154) | 26651 (2628) / 1329 (125) | 20125 (2070) / 961 (83) | 20356 (2068) / 1058 (104) |
| **Non-H atoms: Overall / solvent** | 2246 / 222 | 2159 / 107 | 2180 / 124 | 2192 / 136 |
| **$R_{work}$ / $R_{free}$ [a]** | 0.2027 (0.2168) / 0.2307 (0.2529) | 0.2180 (0.3728) / 0.2469 (0.3932) | 0.2477 (0.3136) / 0.2777 (0.3435) | 0.2231 (0.2782) / 1058 (104) |
| **RMSD from ideal geometry:** | | | | |
| **Bond length (Å) / angles (°)** | 0.014 / 1.33 | 0.007 / 0.94 | 0.015 / 1.36 | 0.007 / 0.88 |
| **Average protein B-factor (Å²)** | 16.68 | 30.86 | 19.80 | 39.03 |
| **Ramachandran: Favored / outlier (%)** | 98.77 / 0.00 | 98.80 / 0.00 | 96.40 / 0.40 | 96.80 / 0.93 |
| **Clashscore** | 2.50 | 0.99 | 3.45 | 0.49 |

[a] Number in parentheses is for the highest resolution shell used in the refinement.

[b] $CC_{1/2}$ = Pearson's intra-dataset correlation coefficient, as described by Karplus and Diederichs.[1]

[c] $R_{merge}$ (= $R_{sym}$) = $\sum_h \sum_1 |I_{h1} - < I_h >| / \sum_h \sum_1 < I_h >$, where $I_{h1}$ is the intensity of the 1st observation of reflection h and $< I_h >$ is the average intensity of reflection h

[d] $R_{meas} = \sum_h \left| \sqrt{(n_h/(n_h-1)} \sum_1 |I_{h1} - < I_h >| \right| / \sum_h \sum_1 < I_h >$ where $n_h$ is the number of observations of reflection $h$

[e] Correlation of experimental intensities with intensities calculated from refined model, as described by Karplus and Diederichs.[1]

**Table S5 |** Human Islets used in this study.

| | | |
|---|---|---|
| **Unique identifiers** | HP-23285, HP-23301, HP-23350, SAMN38094145, SAMN36823227, SAMN35301006, SAMN36510137, HP-24184-01 , HP-24172-01, HP-24095-01, SAMN42008301 | **Aggregate Data** |
| **Origin, Islet isolation centers** | IIDP, Prodo Labs, Wisconsin, imagine pharama | |
| **Donor age (years)** | | Range 36-71, 54.5 ±5.76 (average ± SEM) |
| **Donor sex (M/F)** | | 36.4% F, 63.6% M |
| **Donor BMI (kg/m2)** | | Range 23-38.4, 31±1.4 (average ± SEM) |
| **Donor HbA1c** | | Range 4.4-6.1, 5.11±2.42 (average ± SEM) |
| **Estimated purity (%)** | | Range 75-95, 81.7±3.45 (average ± SEM) |
| **Estimated viability (%)** | | Range 85-98, 95.6±3.11 (average ± SEM) |
| **Total Culture Time (d)** | | 1-4 |

**Table S6 |** Primer used in real-time RT-PCR

| Gene | Forward Primer | Reverse Primer |
|---|---|---|
| ChREBPα | ACTCGGACTCGGACACAGAC | AGGCTCAAGCACTCGAAGAG |
| ChREBPβ | CTGCAGGTCGAGCGGATT | GTCTGTGTCCGAGTCCGAGT |
| PDX1 | ATTCACGAGCCAGTATGACCTTCAC | GAAGACAGACCTGGGATGCACA |
| INS | CGTCCACCCCGTCCACCTC | CCTCCGCCGCCCACACC |
| USP8 | TTCCATTCAATACTTGGACCTGG | CCAAAGAGCCTTTAGCCAATGT |
| RELA | ATGTGGAGATCATTGAGCAGC | CCTGGTCCTGTGTAGCCATT |
| PIN1 | TCAGGCCGAGTGTACTACTTC | TCTTCTGGATGTAGCCGTTGA |
| NFATc1 | CACCGCATCACAGGGAAGAC | GCACAGTCAATGACGGCTC |
| BRAF | AATACACCAGCAAGCTAGATGC | AATCAGTTCCGTTCCCCAGAG |
| SLC2A2 | GCTGCTCAACTAATCACCATGC | TGGTCCCAATTTTGAAAACCCC |
| LDHA | ATGGCAACTCTAAAGGATCAGC | CCAACCCCAACAACTGTAATCT |
| GCK | CCTGGGTGGCACTAACTTCAG | TAGTCGAAGAGCATCTCAGCA |
| FOXO1 | TCGTCATAATCTGTCCCTACACA | CGGCTTCGGCTCTTAGCAAA |
| ALDH1A3 | TGAATGGCACGAATCCAAGAG | CACGTCGGGCTTATCTCCT |
| β-actin | GTCTTCCCCTCCATCGTG | AGGTGTGGTGCCAGATTTTC |

### SUPPLEMENTAL METHODS

#### Organic Synthesis and Characterization

General Information (chemicals, materials and instrumentation)

*Reagents and dry solvents*

All reagents were purchased from ABCR, Acros Organics, Alfa Aesar, Carbolution Chemicals, Carl Roth, Fisher, Fluka, Fluorochem, Merck, Riedel de Häen, Sigma Aldrich, TCI Chemicals, Bernd Kraft or VWR Chemicals and were used without further purification. All dry solvents were purchased from the same supplier with the best quality available.

*Column chromatography*

Compound purification by column chromatography was achieved using glass columns filled with silica gel (particle size 35 – 70 µm, from *Acros Organics*) as stationary phase and eluent mixtures of different solvents as mobile phase. The exact ratios of the solvents are listed in the corresponding synthesis procedures.

*Thin layer chromatography (TLC)*

Thin layer chromatography was performed on silica coated aluminum plates (60 $F_{254}$) from *Merck*. Detection of substances was conducted with UV light (wave length 254 nm or 366 nm). The resulting $R_f$ values including the used solvents are listed in the corresponding synthesis procedures.

*Freeze-drying*

Freeze-drying of the products was carried out with a lyophilizer ALPHA 2-4 LD plus (CHRIST) at an ice condenser temperature of -80 °C. The drying process is favored by a large ice surface and a low ice thickness. To obtain the largest possible ice surface, an aqueous solution of the corresponding substance was frozen in liquid nitrogen under constant rotation. The corresponding frozen compounds were then put for 24 – 72 h on the lyophilizer.

*Reversed-phase liquid chromatography electrospray ionization mass spectrometry (LC-MS)*

Reaction control analyses were performed on a LC-MS system from *Thermo Scientific*. The system consisted of a Thermo Scientific Accela™ (peak detection at 210 nm) and a Thermo Scientific UltiMate™ 3000 (peak detection at 230 nm and 260 nm) equipped with an Eclipse XDB-C18 column (particle size 5 µm, from Agilent) and a *Thermo Scientific* LCQ Fleet™ ESI-MS. For analysis, a linear gradient of solvent B (0.1 % formic acid in acetonitrile) in solvent A (0.1 % formic acid in water) at flow rate of 1 mL $min^{-1}$ and the following gradient program: 0 min (10 % B) → 1 min (10 % B) → 10 min (100 % B) → 12 min (100 % B) → 15 min (10 % B) was used.

*High-resolution mass spectrometry (HRMS)*

HRMS spectra were recorded on an Exactive Plus EMR mass spectrometer from *Thermo Fisher with* an Advion TriVersa NanoMate ESI system from *Advion.*

*Preparative reversed-phase high performance liquid chromatography (prep HPLC)*

Compound purification by HPLC was achieved using the *Prominence UFLC* system from *Shimadzu* (peak detection at 210 nm and 254 nm). The system was equipped with a reversed-phase C18 column from *Phenomenex* (*Luna*® 5 µm C18(2), 100 x 21.20 mm). For purification a linear gradient of solvent B (0.1 % TFA in acetonitrile) in solvent A (0.1 % TFA in water) at a flow rate of 20 mL $min^{-1}$ was used.

*Nuclear magnetic resonance spectroscopy (NMR)*

Nuclear magnetic resonance (NMR) spectra were recorded on a *Bruker Avance II* 400 (400 MHz for $^{1}H$ NMR and 100 MHz for $^{13}C$ NMR) machine. As solvents deuterated chloroform-$d_1$, deuterated DMSO-$d_6$ or deuterated methanol-$d_4$ were used. The chemical shifts $\delta$ are reported in parts per million (ppm). The spectra were referenced to the residual signals of undeuterated solvents ($CDCl_3$: $\delta$ ($^{1}H$) = 7.26 and $\delta$ ($^{13}C$) = 77.16, DMSO: $\delta$ ($^{1}H$) = 2.50 and $\delta$ ($^{13}C$) = 39.52, MeOD: $\delta$ ($^{1}H$) = 4.87 and $\delta$ ($^{13}C$) = 49.00). The coupling constants $J$ are reported in Hertz (Hz). The $^{1}H$ NMR spectral data list the chemical shifts $\delta$, the multiplicities (s: singlet, d doublet, t: triplet, m: multiplet), the coupling constant $J$ and the number of protons. The $^{13}C$ NMR spectra list only the chemical shifts $\delta$.

General Procedures (GP)

*General Procedure A: Synthesis of the phosphate derivative*

The corresponding phenol derivative (1.0 eq) was dissolved in dry DCM (0.5 mL per mmol). Tetrachlormethane (1.0 eq) and triethylamine (1.0 eq) were added. The solution was cooled to 0 °C and dimethyl phosphite (1.5 eq) was added dropwise. The resulting solution was stirred for 16 h, allowing the mixture to slowly reach room temperature. The reaction mixture was washed with 5% $NaHCO_3$ solution (3x), the organic phase was dried over $MgSO_4$ and the solvent was removed under reduced pressure. The crude product was purified by column chromatography.

*General Procedure B: Synthesis of the phosphonate derivative*

Generation of LDA: Dry di-*iso*propyl amine (1.7 eq) was dissolved in dry THF (4.8 mL per mmol of the phosphate derivative) and cooled to -78 °C. *n*-BuLi (2.5 M, 1.5 eq) was added dropwise. The cooling bath was removed and the mixture was stirred for 1 h.

The LDA mixture was cooled to -78 °C again. The respective phosphate derivative (1.0 eq) was dissolved in dry THF (0.7 mL per mmol of the phosphate derivative), cooled to -78 °C and added to the LDA solution. The resulting mixture was stirred for 16 h, allowing the mixture to slowly reach room temperature. The reaction mixture was quenched with saturated $NH_4Cl$ solution (0.6 mL per mmol of the phosphate derivative), the organic phase was separated and the solvent was removed under reduced pressure. The crude product was purified by column chromatography.

*General Procedure C: Amide coupling with HOBt and EDC*

The carboxylic acid derivative (1.0 eq) was dissolved in DCM (50 mL per mmol of the carboxylic acid derivative). EDC (4.0 eq), HOBt (4.0 eq) and DIPEA (6.0 eq) were added. The corresponding amine (2.0 eq) was added and the mixture was stirred for 16 h at room temperature. The reaction mixture was washed with 5% $KHSO_4$ solution (3x) and 5% $NaHCO_3$ solution (3x). The organic phase was dried over $MgSO_4$ and the solvent was removed under reduced pressure.

*General Procedure D: Amide coupling with bromoacetyl bromide*

The amine derivative (1.0 eq) was dissolved in dry DCM (3 – 5 mL per mmol) and cooled to 0 °C. Triethylamine (1.1 eq) and bromoacetyl bromide (1.0 – 1.2 eq) were added and the resulting mixture was stirred for 16 h, allowing the mixture to slowly reach room temperature.

The reaction mixture was washed with saturated $NH_4Cl$ solution (3x) and the organic phase was dried over $MgSO_4$ and the solvent was removed under reduced pressure. The crude product was purified by HPLC.

*General Procedure E: Amide coupling with acid chloride*

Generation of the acid chloride: The acid derivative (1.5 eq.) was dissolved in dry DCM (5 mL per mmol), oxalyl chloride (3.0 eq) and some drops of dry DMF were added. The resulting mixture was stirred for 2 h at 40 °C. After cooling down to room temperature the volatiles were removed under reduced pressure.

The resulting acid chloride was redissolved in dry DCM (5 mL per mmol amine). The amine (1.0 eq.) was dissolved in dry DCM (9 mL per mmol) and cooled to 0 °C. Triethylamine (3.0 eq.) and the acid chloride solution were added and the resulting mixture was stirred for 16 h, allowing the mixture to slowly reach room temperature. The solvent was removed under reduced pressure and the crude product was purified by HPLC.

*General Procedure F: Amide coupling with isobutyl chloroformate*

The acid derivative (1.0 eq.) was dissolved in dry THF (10 mL per mmol acid). *N*-Methylmorpholine (1.0 eq.) was added and the mixture was cooled to -35 °C. Isobutyl chloroformate (1.0 eq) was added and the resulting mixture was stirred for 45 min. The amine (1.0 eq.) and *N*-Methylmorpholine (1.0 eq.) were added and the resulting mixture was stirred for 16 h, allowing the mixture to slowly reach room temperature. The solvent was removed under reduced pressure and the residue was poured into ethyl acetate. The organic phase was washed with saturated $NH_4Cl$ solution (3x), dried over $MgSO_4$ and the solvent was removed under reduced pressure.

*General Procedure G: Williamson ether synthesis*

Reaction in acetone: The respective nucleophile (1.0 eq) was dissolved in acetone (10 – 25 mL per mmol). Potassium carbonate (2.0 eq) was added and the resulting suspension was stirred for 20 min at room temperature. The respective bromide (1.0 – 1.3 eq) was added and the resulting mixture was stirred for 16 h at room temperature. The solvent was evaporated and the residue was suspended in ethyl acetate (25 mL per mmol of the phosphonate derivative). This suspension was washed with saturated $NH_4Cl$ solution (3x) and the organic phase was dried over $MgSO_4$ and the solvent was removed under reduced pressure.

Reaction in DMF: The respective nucleophile (1.0 eq) was dissolved in DMF (2 – 10 mL per mmol). Potassium carbonate (2.0 eq) was added and the resulting suspension was stirred for 20 min at room temperature. The respective bromide (1.0 – 1.3 eq) was added and the resulting mixture was stirred for 16 h at room temperature. The reaction mixture was poured into water (5 mL per mL DMF) and was extracted with diethyl ether (3x). The combined organic phases were dried over $MgSO_4$ and the solvent was removed under reduced pressure.

*General Procedure H: Deprotection of the phosphonates*

The protected phosphonate derivative (1.0 eq) was dissolved in dry DCM (6 mL per mmol) and cooled to 0 °C. TMSBr (5.0 – 50.0 eq) was added and the resulting solution was stirred for 4 h, allowing the mixture to slowly reach room temperature. The solvent was removed under reduced pressure. The residue was re-dissolved in a mixture MeOH/$H_2O$ (3:1, 6 mL per mmol of the protected phosphonate derivative) and stirred for 1 h at room temperature. The solvent was removed under reduced pressure and the crude product was purified by HPLC.

*General Procedure J: Substitution with 2-(Boc-amino)ethyl bromide*

The heteroaromatic (1.0 eq) was dissolved in DMF (8 mL per mmol). Potassium carbonate (2.4 eq) was added and the resulting suspension was stirred for 20 min at room temperature. 2-(Boc-amino)ethyl bromide (1.0 eq) was added and the resulting mixture was stirred for 48 h at room temperature. The reaction mixture was poured into water (5 mL per mL DMF) and was extracted with ethyl acetate (3x). The combined organic phases were dried over $MgSO_4$ and the solvent was removed under reduced pressure.

*General Procedure K: Boc protection*

The amine (1.0 eq.) and Di-*tert* butyl dicarbonate (1.03 eq.) was suspended in THF (4.3 mL per mmol amine) and cooled down to 0 °C. $NaHCO_3$ (1.11 eq.) was dissolved in $H_2O$ (2.2 mL per mmol amine) and added to the suspension dropwise. The resulting mixture was stirred for 16 h, allowing the mixture to slowly reach room temperature. The solvent was removed under reduced pressure and the residue was poured into ethyl acetate (10 mL per mmol amine). The organic phase was washed with 0.5M HCl (3x), $H_2O$ (1x) and brine (1x), dried over $MgSO_4$ and the solvent was removed under reduced pressure.

*General Procedure L: Boc deprotection*

Deprotection with dioxane: The Boc protected amine (1.0 eq.) was dissolved in dioxane (0.1 – 1.0 mL), 4M HCl/dioxane was added and the reaction was stirred for 3 h at room temperature. The suspension was poured in diethyl ether and 1 M NaOH (20 mL per mmol amine) was added. The phases were separated and the aqueous phase was extracted with diethyl ether (3x). The combined organic phases were dried over $MgSO_4$ and the solvent was removed under reduced pressure.

Deprotection with TFA: The Boc protected amine (1.0 eq.) was dissolved in DCM/TFA (1:1, 5 mL per mmol) and stirred for 16 h at room temperature. The solvent was removed under reduced pressure.

*General Procedure M: Henry reaction with sodium hydroxide*

The aldehyde (1.0 eq.) and nitromethane (1.1 eq.) were dissolved in ethanol (1.75 mL per mmol aldehyde) and cooled down to 0 °C. 10M NaOH (100 µL per mmol aldehyde) was added slowly and the resulting mixture was stirred for 2 h at 0 °C. The mixture was quenched with HCl/ $H_2O$ (1:1, 5 mL per mmol aldehyde) and stirred for 1 h at 0 °C. The resulting soild was filtered, washed with $H_2O$ and dried under reduced pressure.

*General Procedure N: Reduction with lithium aluminium hydride*

The nitrovinyl derivative (1.0 eq.) was dissolved in dry THF (2 mL per mmol nitrovinyl derivative). Lithium aluminium hydride was suspended in dry THF (2 mL per mmol nitrovinyl derivative) and cooled down to 0 °C). The nitrovinyl derivative solution was added slowly and the resulting mixture was stirred for 20 min at 0 °C and then for 3 h at 45 °C. The mixture was then cooled down to 0 °C again and quenched with 10 % NaOH solution (1.2 mL per mmol nitrovinyl derivative). The mixture was filtered over Celite and the filter cake was washed with ethyl acetate. The crude product was dried under reduced pressure and then purified by column chromatography or HPLC.

*General Procedure O: Copper mediated cross coupling*

Activated copper (2.6 – 3.9 eq.) was suspended in dry DMSO or DMF (1.6 – 2.6 mL per mmol iodobenzene derivative). The idoobenzene derivative (1.0 eq.) and ethyl bromodifluoroacetate (1.0 – 1.5 eq.) were added and the reaction mixture was stirred for 16 h at 60 °C. After cooling down to room temperature the reaction mixture was poured into ice/ saturated $NH_4Cl$ solution (1:1), the inorganic solid was filtered out and the filtrate was extracted with diethyl ether. The combined organic phases were dried over $MgSO_4$ and the solvent was removed under reduced pressure. The crude product was purified by column chromatography.

Copper activation: Copper was suspended in 1M HCl solution (0.4 mL per mmol copper). The suspension was stirred for 10 min at room temperature and the solvent was filtered out. The procedure was repeated one after another with $H_2O$, MeOH and Aceton. At the end the activated copper was dried under reduced pressure.

*General Procedure P: Reduction with sodium boronhydride*

The difluoroacetate derivative (1.0 eq.) was dissolved in MeOH (3.4 – 3.8 mL per mmol) und cooled down to 0 °C. $NaBH_4$ (1.0 eq.) was added and the mixture was stirred for 1 h at 0 °C. The reaction mixture was quenched with $H_2O$/ saturated $NH_4Cl$ solution (1:1, 1.4 mL per mmol) and with ethyl acetate extracted. The combined organic phases were dried over $MgSO_4$ and the solvent was removed under reduced pressure.

*General Procedure Q: Amination*

The alcohol (1.0 eq.) was dissolved in dry acetonitrile (1.8 mL per mmol alcohol) and pyridine (1.6 eq.) was added. The mixture was cooled down to 0 °C and $TF_2O$ (1.1 eq.) was added slowly. The mixture was stirred for 15 min at room temperature. $NH_4OH$ (28 %, 1.8 mL per mmol alcohol) was added and the mixture was stirred for 16 h at room temperature. The reaction mixture was extracted with DCM, the combined organic phases were dried over $MgSO_4$ and the solvent was removed under reduced pressure.

*General Procedure R: Esterification with chloromethyl pivalate*

The acid (1.0 eq.) was dissolved in dry DMF (15 mL per mmol acid) and triethylamine (2.0 eq.) was added. After 10 min stirring at room temperature chloromethyl pivalate (10.0 eq.) was added and the reaction mixture was stirred for 16 h at 60 °C. After cooling down to room temperature the mixture was poured into $H_2O$ and extracted with diethyl ether. The combined organic phases were dried over $MgSO_4$ and the solvent was removed under reduced pressure.

Detailed Synthetic Procedures and Characterization of Compounds

**Synthesis of intermediates for the derivatives 2 – 4**

Synthesis of dimethyl phenyl phosphate (**S-1**)

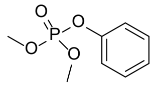
**S-1**

The phosphate **S-1** was synthesized via GP **A** using phenol (9.41 g, 100 mmol), tetrachlormethane (9.6 mL, 100 mmol), triethylamine (14.9 mL, 100 mmol) and dimethyl phosphite (13.8 mL, 150 mmol) in dry DCM (50 mL). The crude product was purified by column chromatography (cyclohexane/ethyl acetate 2:1 → 1:1) to obtain **S-1** (11.82 g, 58.5 mmol, 59 %) as a colorless oil. **TLC** (cyclohexane:ethyl acetate, 1:1 v/v): $R_f$ = 0.4. **LC-MS (ESI)**: $t_R$ = 7.13 min; *m/z* = 203.03 $[M + H]^+$. **$^1$H NMR** (400 MHz, $CDCl_3$): $\delta$ = 7.26 – 7.22 (m, 2H), 7.13 – 7.06 (m, 3H), 3.73 (s, 6H). **$^{13}$C NMR** (101 MHz, $CDCl_3$): $\delta$ = 150.81, 129.95, 125.33, 120.05, 120.00, 55.01. **HRMS (ESI)**: *m/z* = 203.0468 calcd. for $[C_8H_{11}O_4P + H]^+$; found: 203.0462.

Synthesis of dimethyl 2-hydroxyphenylphosphonate (**S-2**)

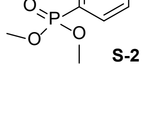
**S-2**

The phosphonate **S-2** was synthesized via GP **B** using di-*iso*propyl amine (13.8 mL, 98.5 mmol), *n*-BuLi (2.5 M, 34.8 mL, 87.8 mmol) and phosphate **S-1** (11.80 g, 58.4 mmol) in dry THF (280 mL + 40 mL). The crude product was purified by column chromatography (cyclohexane/ethyl acetate 2:1) to obtain **S-2** (9.92 g, 49.1 mmol, 84 %) as a white solid. **TLC** (cyclohexane:ethyl acetate, 2:1 v/v): $R_f$ = 0.4. **LC-MS (ESI)**: $t_R$ = 6.02 min; *m/z* = 202.98

[M + H]$^+$. **$^1$H NMR** (400 MHz, $CDCl_3$): $\delta$ = 10.06 (s, 1H), 7.47 – 7.43 (m, 1H), 7.35 (ddd, *J* = 14.3, 7.7, 1.7 Hz, 1H), 6.99 – 6.90 (m, 2H), 3.74 (d, *J* = 11.6 Hz, 6H). **$^{13}$C NMR** (101 MHz, $CDCl_3$): $\delta$ = 162.77, 135.94, 132.01, 120.22, 118.29, 77.80, 77.16, 53.47. **HRMS (ESI)**: *m/z* = 203.0468 calcd. for [$C_8H_{11}O_4P$ + H]$^+$; found: 203.0461.

#### Synthesis of 2-(2-(methoxyphosphono)phenoxy)acetic acid (**S-3**)

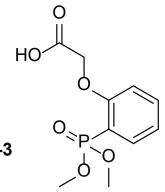


**S-3**

The phosphonate derivative **S-3** was synthesized via GP **G** using phosphonate **S-2** (1.822 g, 9.00 mmol), potassium carbonate (2.484 g, 18.0 mmol) and methyl bromoacetate (0.95 mL, 9.90 mmol) in acetone (225 mL). The crude intermediate was dissolved in a mixture THF/ methanol (1:1, 300 mL). Lithium hydroxide (0.647 g, 27.0 mmol, 3.0 eq) was added and the solution was stirred for 16 h at room temperature. The reaction mixture was acidified with 1M HCl solution (to a pH < 3) and the solvent was evaporated under reduced pressure. The crude product was purified by column chromatography (ethyl acetate/methanol 1:1) to obtain **S-3** (1.56 g, 6.00 mmol, 67 % over two steps) as a white solid. **TLC** (ethyl acetate:methanol, 1:1 v/v): $R_f$ = 0.5. **LC-MS (ESI)**: $t_R$ = 4.92 min; *m/z* = 261.00 [M + H]$^+$. **$^1$H NMR** (400 MHz, $CDCl_3$): $\delta$ = 10.93 (s, 1H), 7.65 – 7.59 (m, 1H), 7.55 – 7.51 (m 1H), 7.12 – 7.06 (m, 1H), 6.97 – 6.92 (m, 1H), 4.72 (d, J = 2.6 Hz, 2H), 3.78 (dd, *J* = 11.4, 2.9 Hz, 6H). **$^{13}$C NMR** (101 MHz, $CDCl_3$): $\delta$ = 169.86, 160.32, 135.53, 134.05, 122.67, 114.19, 67.73, 53.84. **HRMS (ESI)**: *m/z* = 261.0523 calcd. for [$C_{10}H_{13}O_6P$ + H]$^+$; found: 261.0514.

#### Synthesis of dimethyl (2-(2-((2-(9H-purin-9-yl)ethyl)amino)-2-oxoethoxy)phenyl)phosphonate (**S-4**)

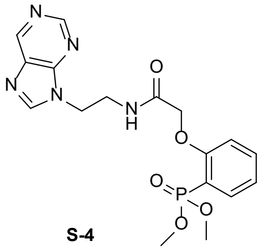


**S-4**

The phenylphosphonic acid derivative **S-4** was synthesized via GP **J**, GP **L** and GP **E**. GP **J** employed purine (0.48 g, 4.00 mmol), potassium carbonate (1.33 g, 9.60 mmol), 2-(Boc-amino)ethyl bromide (0.90 g, 4.00 mmol) in DMF (32 mL). The Boc protected intermediate was deprotected via GP **L** using DCM/TFA (20 mL). GP **E** converted the crude amine into the amide. The acid chloride was generated using the acid **S-3** (1.56 g, 6.00 mmol), oxaly chloride (1.03 mL, 12.00 mmol), dry DMF (3 drops) in dry DCM (40 mL). In addition the crude amine and triethylamine (1.66 mL, 12.00 mmol) in dry DCM (24 mL + 36 mL) were used. The crude product was purified by HPLC to obtain **S-4** (0.34 g, 0.83 mmol, 21 % over three steps) as a slightly yellow solid. **LC-MS (ESI)**: $t_R$ = 5.34 min; *m/z* = 406.08 [M + H]$^+$, calcd. for $C_{17}H_{20}N_5O_5P$: 405.12. **$^1$H NMR** (400 MHz, $CDCl_3$): $\delta$ = 9.81 (s, 1H), 9.33 (s, 1H), 9.03 (s, 1H), 8.64 (s, 1H), 7.60 – 7.54 (m, 2H), 7.16 – 7.11 (m, 1H), 7.01 – 6.93 (m, 1H), 4.77 (s, 2H), 4.67 – 4.59 (m, 2H), 3.89 – 3.84 (m, 6H), 3.72 – 3.68 (m, 2H). **$^{13}$C NMR** (101 MHz, $CDCl_3$): $\delta$ = 169.59, 159.79, 153.70, 141.72, 135.30, 133.49, 122.50, 116.08, 113.66, 67.20, 53.42, 44.09, 38.32.

#### Synthesis of dimethyl (2-(2-((2-(6-amino-9H-purin-9-yl)ethyl)amino)-2-oxoethoxy)phenyl)phosphonate (**S-5**)

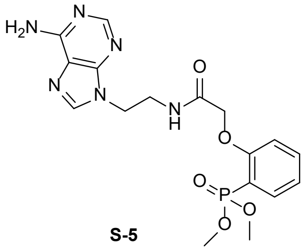


**S-5**

The phenylphosphonic acid derivative **S-5** was synthesized via GP **J**, GP **L** and GP **E**. GP **J** employed adenine (0.54 g, 4.00 mmol), potassium carbonate (1.33 g, 9.60 mmol), 2-(Boc-amino)ethyl bromide (0.90 g, 4.00 mmol.) in DMF (32 mL). The Boc protected intermediate was deprotected via GP **L** using DCM/TFA (20 mL). GP **E** converted the crude amine into the amide. The acid chloride was generated using the acid **S-3** (1.56 g, 6.00 mmol), oxaly chloride (1.03 mL, 12.00 mmol), dry DMF (3 drops) in dry DCM (40 mL). In addition the crude amine and triethylamine (1.66 mL, 12.00 mmol) in dry DCM (24 mL + 36 mL) were used. The crude product was purified by HPLC to obtain **S-5** (0.37 g, 0.87 mmol, 22 % over three steps) as a white solid. **LC-MS (ESI)**: $t_R$ = 4.85 min; *m/z* = 421.19 [M + H]$^+$, calcd. for $C_{17}H_{21}N_6O_5P$: 420.13. **$^1$H NMR** (400 MHz, $CDCl_3$): $\delta$ = 9.87 (s, 1H), 8.97 (t, *J* = 6.2 Hz, 1H), 8.60 (s, 1H), 8.36 – 8.25 (m, 2H), 7.64 – 7.56 (m, 2H), 7.19 – 7.12 (m, 1H), 6.99 – 6.33 (m, 1H), 4.66 (s, 2H), 4.54 (t, *J* = 5.3 Hz, 2H), 3.99 – 3.88 (m, 2H), 3.76 (dd, *J* = 11.6, 1.8 Hz, 6H). **$^{13}$C NMR** (101 MHz, $CDCl_3$): $\delta$ = 171.28, 161.00, 159.57, 150.95, 149.54, 145.60, 144.20, 136.37, 134.48, 123.17, 116.91, 114.06, 67.71, 54.04, 44.78, 39.28.

#### Synthesis of dimethyl (2-(2-((2-(2-amino-9H-purin-9-yl)ethyl)amino)-2-oxoethoxy)phenyl)phosphonate (**S-6**)

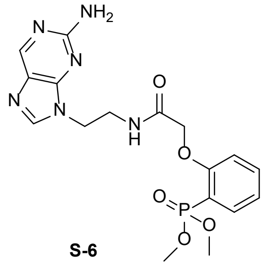

**S-6**

The phenylphosphonic acid derivative **S-6** was synthesized via GP **J**, GP **L** and GP **E**. GP **J** employed 9*H*-purine-amine (425 mg, 2.50 mmol, 1.0 Äq), potassium carbonate (830 mg, 6.00 mmol), 2-(Boc-amino)ethyl bromide (560 mg, 2.50 mmol) in DMF (20 mL). The Boc protected intermediate was deprotected via GP **L** using DCM/TFA (12.5 mL). GP **E** converted the crude amine into the amide. The acid chloride was generated using the acid **S-3** (976 mg, 3.75 mmol), oxaly chloride (643 µL, 7.50 mmol), dry DMF (1 drop) in dry DCM (20 mL). In addition the crude amine and triethylamine (1.04 mL, 7.50 mmol) in dry DCM (22 mL + 34 mL) were used. The crude product was purified by HPLC to obtain **S-6** (74 mg, 0.18 mmol, 7 % over three steps) as a white solid. **LC-MS (ESI)**: $t_R$ = 4.77 min; *m/z* = 421.11 [M + H]$^+$, calcd. for $C_{17}H_{21}N_6O_5P$: 420.13. **$^1$H NMR** (400 MHz, $CDCl_3$): $\delta$ = 9.27 (s, 1H), 8.49 (s, 1H), 8.14 (s, 1H), 7.69 – 7.54 (m, 3H), 7.16 – 7.12 (m, 1H), 7.02 – 6.91 (m, 2H), 4.65 (s, 2H), 4.36 (t, *J* = 5.2 Hz, 2H), 3.84 – 3.79 (m, 2H), 3.77 – 3.70 (m, 6H).

### Synthesis of the derivatives 2 – 4

#### Synthesis of (2-(2-((2-(9H-purin-9-yl)ethyl)amino)-2-oxoethoxy)phenyl)phosphonic acid (**2**)

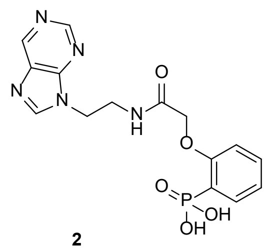

**2**

The phenylphosphonic acid derivative **2** was synthesized via GP **H** using the protected phosphonate **S-4** (337 mg, 0.834 mmol) and TMSBr (0.55 mL, 4.170 mmol) in dry DCM (5.0 mL) and MeOH/$H_2O$ (3:1, 5.0 mL). The crude product was purified by HPLC to obtain **2** (47 mg, 0.125 mmol, 15 %) as a white solid. **LC-MS (ESI)**: $t_R$ = 2.28 min; *m/z* = 378.04 [M + H]$^+$, calcd. for $C_{15}H_{16}N_5O_5P$: 377.09. **$^1$H NMR** (400 MHz, DMSO): $\delta$ = 9.39 (t, *J* = 5.8 Hz, 1H), 9.14 (s, 1H), 8.90 (s, 1H), 8.51 (s, 1H), 7.63 – 7.58 (m, 1H), 7.48 (t, *J* = 7.9 Hz, 1H), 7.12 – 7.05 (m, 2H), 4.56 (s, 2H), 4.41 (t, *J* = 5.9 Hz, 2H), 3.62 (m, 2H). **$^{13}$C NMR** (101 MHz, DMSO): $\delta$ = 168.53, 158.97, 151.48, 133.52, 133.21, 132.53, 122.91, 121.31, 121.18, 113.45, 67.84, 42.61, 37.94. **HRMS (ESI)**: *m/z* = 378.0962 calcd. for [$C_{15}H_{16}N_5O_5P$ + H]$^+$; found: 378.0960.

#### Synthesis of (2-(2-((2-(6-amino-9H-purin-9-yl)ethyl)amino)-2-oxoethoxy)phenyl)phosphonic acid (**3**)

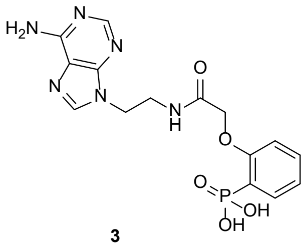

**3**

The phenylphosphonic acid derivative **3** was synthesized via GP **H** using the protected phosphonate **S-5** (365 mg, 0.869 mmol) and TMSBr (0.57 mL, 4.325 mmol) in dry DCM (5.0 mL) and MeOH/$H_2O$ (3:1, 5.0 mL). The crude product was purified by HPLC to obtain **3** (202 mg, 0.514 mmol, 59 %) as a white solid. **LC-MS (ESI)**: $t_R$ = 2.09 min; *m/z* = 393.12 [M + H]$^+$, calcd. for $C_{15}H_{17}N_6O_5P$: 392.10. **$^1$H NMR** (400 MHz, $CDCl_3$): $\delta$ = 9.41 (s, 1H), 9.02 (s, 1H), 8.34 (d, *J* = 10.1 Hz, 1H), 7.66 – 7.58 (m, 1H), 7.47 (t, *J* = 7.9 Hz, 1H), 7.13 – 7.01 (m, 3H), 4.57 (s, 2H), 4.33 (t, *J* = 5.8 Hz, 2H), 3.61 – 3.53 (m, 2H), 3.14 – 3.03 (m, 1H), 1.17 (t, *J* = 7.3 Hz, 1H). **$^{13}$C NMR** (101 MHz, $CDCl_3$): $\delta$ = 168.69, 158.97, 151.25, 145.89, 143.74, 133.12, 132.51, 123.11, 121.18, 113.33, 67.86, 43.06, 38.26. **HRMS (ESI)**: *m/z* = 393.1071 calcd. for [$C_{15}H_{17}N_6O_5P$ + H]$^+$; found: 393.1070.

#### Synthesis of (2-(2-((2-(2-amino-9H-purin-9-yl)ethyl)amino)-2-oxoethoxy)phenyl)phosphonic acid (**4**)

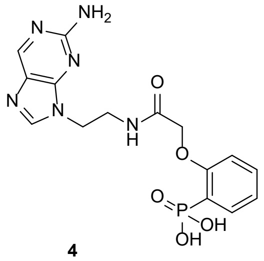

**4**

The phenylphosphonic acid derivative **4** was synthesized via GP **H** using the protected phosphonate **S-6** (68 mg, 0.162 mmol) and TMSBr (0.11 mL, 0.811 mmol) in dry DCM (6.0 mL) and MeOH/$H_2O$ (3:1, 6.0 mL). The crude product was purified by HPLC to obtain **4** (31 mg, 0.078 mmol, 48 %) as a white solid. **LC-MS (ESI)**: $t_R$ = 2.15 min; *m/z* = 393.04 [M + H]$^+$, calcd. for $C_{15}H_{17}N_6O_5P$: 392.10. $^1$**H NMR** (400 MHz, DMSO): *δ* = 9.37 (t, *J* = 5.6 Hz, 1H), 8.69 (s, 1H), 8.17 (s, 1H), 7.65 – 7.56 (m, 2H), 7.49 (t, *J* = 8.0 Hz, 1H), 7.14 – 7.03 (m, 3H), 4.59 (s, 2H), 4.18 (t, *J* = 5.9 Hz, 2H), 3.58 – 3.51 (m, 2H). $^{13}$**C NMR** (101 MHz, DMSO): *δ* = 168.56, 158.94, 145.88, 133.18, 132.51, 126.41, 121.20, 114.30, 67.83, 42.11, 37.78. **HRMS (ESI)**: *m/z* = 393.1071 calcd. for [$C_{15}H_{17}N_6O_5P$ + H]$^+$; found: 393.1069.

#### Synthesis of the derivatives 5 – 6

##### Synthesis of (2-(2-((2-(1H-indol-3-yl)ethyl)amino)-2-oxoethoxy)phenyl)phosphonic acid (**5**)

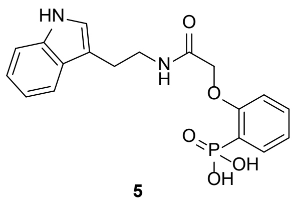

**5**

The phenylphosphonic acid derivative **5** was synthesized via GP **F** and GP **H**. GP **F** employed phosphonate **S-3** (260 mg, 1.00 mmol), *N*-Methylmorpholine (111 µL, 1.00 mmol) and isobutyl chloroformate (130 µL, 1.00 mmol) in dry THF (10 mL) as well as *N*-Methylmorpholine (111 µL, 1.00 mmol) and tryptamine (160 mg, 1.00 mmol). The crude intermediate was deprotected via GP **H** using TMSBr (1.32 mL, 10.00 mmol) in dry DCM (8 mL) and MeOH/$H_2O$ (3:1, 8 mL). The crude product was purified by HPLC to obtain **5** (118 mg, 0.31 mmol, 31 % over two steps) as a white solid. **LC-MS (ESI)**: $t_R$ = 5.93 min; *m/z* = 375.05 [M + H]$^+$, calcd. for $C_{18}H_{19}N_2O_5P$: 374.10. $^1$**H NMR** (400 MHz, DMSO): *δ* = 10.80 – 10.75 (m, 1H), 9.22 (t, *J* = 5.6 Hz, 1H), 7.69 – 7.62 (m, 1H), 7.56 – 7.46 (m, 2H), 7.32 (d, *J* = 8.1 Hz, 1H), 7.19 – 7.10 (m, 2H), 7.12 – 7.01 (m, 2H), 6.95 (t, *J* = 7.4 Hz, 1H), 4.63 (s, 2H), 3.42 (q, *J* = 8.4, 7.3 Hz, 2H), 2.87 (t, *J* = 7.7 Hz, 2H). $^{13}$**C NMR** (101 MHz, DMSO): *δ* = 167.79, 158.96, 136.21, 133.16, 132.65, 127.15, 122.66, 120.89, 118.22, 113.30, 111.65, 111.36, 67.79, 25.03. **HRMS (ESI)**: *m/z* = 375.1105 calcd. for [$C_{18}H_{19}N_2O_5P$ + H]$^+$; found: 375.1104.

##### Synthesis of (4-(2-oxo-2-(phenethylamino)ethoxy)phenyl)phosphonic acid (**6**)

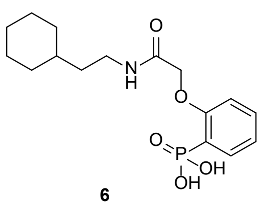

**6**

The phenylphosphonic acid derivative **6** was synthesized via GP **F** and GP **H**. GP **F** employed phosphonate **S-3** (260 mg, 1.00 mmol), *N*-Methylmorpholine (111 µL, 1.00 mmol) and isobutyl chloroformate (130 µL, 1.00 mmol) in dry THF (10 mL) as well as *N*-Methylmorpholine (111 µL, 1.00 mmol) and cyclohexylethylamine (150 mg, 1.00 mmol). The crude intermediate was deprotected via GP **H** using TMSBr (1.32 mL, 10.00 mmol) in dry DCM (8 mL) and MeOH/$H_2O$ (3:1, 8 mL). The crude product was purified by HPLC to obtain **6** (119 mg, 0.35 mmol, 35 % over two steps) as a white solid. **LC-MS (ESI)**: $t_R$ = 6.64 min; *m/z* = 342.19 [M + H]$^+$, calcd. for $C_{18}H_{19}N_2O_5P$: 341.14. $^1$**H NMR** (400 MHz, DMSO): *δ* = 8.88 (t, *J* = 5.6 Hz, 1H), 7.65 – 7.58 (m, 1H), 7.49 (td, *J* = 7.9, 1.7 Hz, 1H), 7.14 – 7.10 (m, 1H), 7.05 (td, *J* = 7.4, 2.9 Hz, 1H), 4.61 (s, 2H), 3.13 (q, *J* = 6.5 Hz, 2H), 1.67 – 1.53 (m, 5H), 1.32 (q, *J* = 7.1 Hz, 2H), 1.21 – 1.08 (m, 4H), 0.86 – 0.75 (m, 2H). $^{13}$**C NMR** (101 MHz, DMSO): *δ* = 167.57, 158.83, 133.06, 132.53, 123.14, 121.35, 121.12, 113.13, 67.56, 36.29, 36.01, 34.34, 32.59, 26.12, 25.68. **HRMS (ESI)**: *m/z* = 342.1465 calcd. for [$C_{18}H_{19}N_2O_5P$ + H]$^+$; found: 342.1463.

#### Synthesis of intermediates for the derivatives 7 – 8

##### Synthesis of 1-(2-nitrovinyl)naphthalene (**S-7**)

The nitrovinyl derivative **S-7** was synthesized via GP **M** using 1-Naphthaldehyde (1.36 mL, 10.00 mmol), nitromethane (0.51 mL, 11.00 mmol), 10M NaOH solution (1.01 mL) in ethanol (17.5 mL) and HCl/$H_2O$ (1:1, 50 mL). **S-7** (1.87 g, 9.39 mmol, 94 %) was obtained as a yellow solid. $^1$**H NMR** (400 MHz, $CDCl_3$): *δ* = 8.80 (d, *J* = 13.4 Hz, 1H), 8.11 (d, *J* = 8.4 Hz, 1H), 7.99 (d, *J* = 8.2 Hz, 1H), 7.91 (d, *J* = 8.1 Hz, 1H), 7.73 (d, *J* = 7.3 Hz, 1H), 7.66 – 7.56 (m, 3H), 7.53 7.55 – 7.47 (m, 1H). $^{13}$**C NMR** (101 MHz, $CDCl_3$): *δ* = 138.60, 136.21, 133.88, 132.67, 131.68, 129.18, 127.85, 127.09, 126.91, 126.50, 125.51, 123.08.

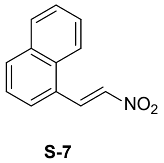

**S-7**

#### Synthesis of 2-(2-nitrovinyl)naphthalene (**S-8**)

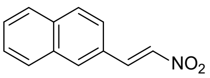

**S-8**

The nitrovinyl derivative **S-8** was synthesized via GP **M** using 2-Naphthaldehyde (1.56 g, 10.00 mmol), nitromethane (0.51 mL, 11.00 mmol), 10M NaOH solution (1.01 mL) in ethanol (17.5 mL) and HCl/$H_2O$ (1:1, 50 mL). **S-8** (1.54 g, 7.72 mmol, 77 %) was obtained as a yellow solid. **$^1$H-NMR** (400 MHz, $CDCl_3$): *δ* = 8.13 (d, *J* = 13.6 Hz, 1H), 8.00 (s, 1H), 7.90 – 7.86 (m, 3H), 7.70 (d, *J* = 13.6 Hz, 1H), 7.61 – 7.54 (m, 3H). **$^{13}$C-NMR** (101 MHz, $CDCl_3$): *δ* = 139.35, 137.25, 135.03, 133.32, 132.39, 129.48, 128.96, 128.51, 128.06, 127.66, 127.40, 123.44.4

#### Synthesis of 2-(naphthalen-1-yl)ethan-1-amine (**S-9**)

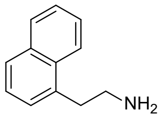

**S-9**

The amine **S-9** was synthesized via GP **N** using nitrovinyl derivative **S-7** (1.89 g, 9.50 mmol) and lithium aluminium hydride (0.72 g, 19.00 mmol) in dry THF (19 mL + 19 mL). The crude product was purified by HPLC to obtain **S-9** (0.78 g, 4.56 mmol, 48 %) as a white solid. **LC-MS (ESI)**: $t_R$ = 5.27 min; *m/z* = 171.86 $[M + H]^+$, calcd. for $C_{12}H_{13}N$: 171.10.
**$^1$H NMR** (400 MHz, DMSO): *δ* = 8.12 (d, *J* = 8.3 Hz, 1H), 8.03 (s, 2H), 7.96 (d, *J* = 7.6 Hz, 1H), 7.85 (d, *J* = 8.0 Hz, 1H), 7.62 – 7.53 (m, 2H), 7.49 – 7.41 (m, 2H), 3.37 – 3.33 (m, 2H), 3.17 – 3.10 (m, 2H). **$^{13}$C NMR** (101 MHz, DMSO): *δ* = 133.51, 133.32, 131.27, 128.76, 127.48, 126.96, 126.39, 125.86, 125.68, 123.33, 30.25.

#### Synthesis of 2-(naphthalen-2-yl)ethan-1-amine (**S-10**)

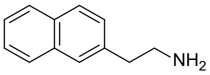

**S-10**

The amine **S-10** was synthesized via GP **N** using nitrovinyl derivative **S-8** (1.63 g, 8.16 mmol) and lithium aluminium hydride (0.62 g, 16.32 mmol) in dry THF (16 mL + 16 mL). The crude product was purified by HPLC to obtain **S-10** (0.71 g, 4.16 mmol, 51 %) as a white solid. **LC-MS (ESI)**: $t_R$ = 5.60 min; *m/z* = 171.84 $[M + H]^+$, calculated for $C_{12}H_{13}N$: 171.10. **$^1$H-NMR** (400 MHz, MeOD): *δ* = 7.90 – 7.85 (m, 3H), 7.78 (s, 1H), 7.53 – 7.47 (m, 2H), 7.43 (dd, *J* = 8.4, 1.9 Hz, 1H), 3.32 – 3.29 (m, 2H), 3.15 (t, *J* = 7.6 Hz, 2H).

### Synthesis of the derivatives 7 – 8

#### Synthesis of 2-(2-((2-(2-naphthyl)ethyl)amino)-2-oxoethoxy)phenyl)phosphonic acid (**7**)

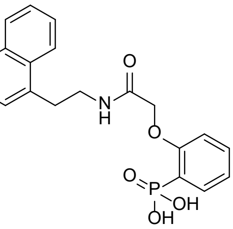

**7**

The phenylphosphonic acid derivative **7** was synthesized via GP **F** and GP **H**. GP **F** employed phosphonate **S-3** (130 mg, 0.50 mmo), *N*-Methylmorpholine (56 µL, 0.50 mmol) and isobutyl chloroformate (65 µL, 0.50 mmol) in dry THF (5.1 mL) as well as *N*-Methylmorpholine (56 µL, 0.50 mmol) and the amine **S-9** (86 mg, 0.61 mmol). The crude intermediate was deprotected via GP **H** using TMSBr (.33 mL, 2.50 mmol) in dry DCM (6 mL) and MeOH/$H_2O$ (3:1, 6 mL). The crude product was purified by HPLC to obtain **7** (31 mg, 0.08 mmol, 16 % over two steps) as a white solid. **LC-MS (ESI)**: $t_R$ = 6.98 min; *m/z* = 386.07 $[M + H]^+$, calcd. for $C_{20}H_{20}NO_5P$: 385.11. **$^1$H NMR** (400 MHz, DMSO): *δ* = 9.39 (t, *J* = 5.7 Hz, 1H), 8.19 (d, *J* = 8.0 Hz, 1H), 7.90 (d, *J* = 7.2 Hz, 1H), 7.78 (d, *J* = 7.1 Hz, 1H), 7.70 – 7.62 (m, 1H), 7.58 – 7.48 (m, 3H), 7.41 – 7.32 (m, 2H), 7.15 (t, *J* = 7.3 Hz, 1H), 7.09 (td, *J* = 7.5, 2.6 Hz, 1H), 4.64 (s, 2H), 3.49 – 3.40 (m, 2H), 3.27 – 3.15 (m, 2H). **$^{13}$C NMR** (101 MHz, DMSO): *δ* = 168.03, 159.04, 135.26, 133.45, 133.20, 132.67, 131.52, 128.62, 126.83, 126.59, 126.12, 125.63, 123.63, 123.14, 121.11, 113.10, 67.84, 48.65, 32.40. **HRMS (ESI)**: *m/z* = 386.1152 calcd. for $[C_{20}H_{20}NO_5P + H]^+$; found: 386.1149.

#### Synthesis of 2-(2-((2-(1-naphthyl)ethyl)amino)-2-oxoethoxy)phenyl)phosphonic acid (**8**)

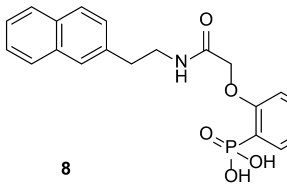

8

The phenylphosphonic acid derivative **8** was synthesized via GP **F** and GP **H**. GP **F** employed phosphonate **S-3** (159 mg, 0.61 mmol), *N*-Methylmorpholine (68 µL, 0.61 mmol) and isobutyl chloroformate (79 µL, 0.61 mmol) in dry THF (6.1 mL) as well as *N*-Methylmorpholine (68 µL, 0.61 mmol) and the amine **S-10** (104 mg, 0.61 mmol). The crude intermediate was deprotected via GP **H** using TMSBr (0.40 mL, 3.05 mmol) in dry DCM (3.7 mL) and MeOH/$H_2O$ (3:1, 3.7 mL). The crude product was purified by HPLC to obtain **8** (59 mg, 0.15 mmol, 25 % over two steps) as a white solid. **LC-MS (ESI)**: $t_R$ = 6.99 min; *m/z* = 386.14 $[M + H]^+$, calcd. for $C_{20}H_{20}NO_5P$: 385.11. **$^1$H NMR** (400 MHz, DMSO): $\delta$ = 9.25 (t, *J* = 5.5 Hz, 1H), 7.83 (dd, *J* = 22.9, 7.8 Hz, 3H), 7.74 – 7.64 (m, 2H), 7.51 – 7.40 (m, 3H), 7.38 (d, *J* = 8.4 Hz, 1H), 7.16 – 7.03 (m, 2H), 4.63 (s, 2H), 3.47 (t, *J* = 6.8 Hz, 2H), 2.95 (t, *J* = 7.5 Hz, 2H). **$^{13}$C NMR** (101 MHz, DMSO): $\delta$ = 168.15, 159.24, 137.24, 133.46, 133.02, 131.96, 127.98, 127.70, 126.93, 126.17, 125.58, 123.31, 121.47, 121.34, 113.61, 67.98, 40.25, 35.31. **HRMS (ESI)**: *m/z* = 386.1152 calcd. for $[C_{20}H_{20}NO_5P + H]^+$; found: 386.1151.

#### Synthesis of intermediates for the derivatives 9 – 11

##### Synthesis of N-(4-hydroxyphenethyl)-2-bromoacetamide (**S-11**)

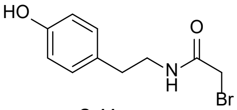

S-11

The bromoacetamide **S-11** was synthesized via GP **D** using tyramine (0.69 g, 5.00 mmol), triethylamine (0.76 mL, 5.50 mmol) and bromoacetyl bromide (0.44 mL, 5.00 mmol) in dry DCM (25 mL) and additional dry methanol (2.5 mL). The crude product was purified by HPLC to obtain **S-11** (0.62 g, 2.41 mmol, 48 %) as a white solid. **LC-MS (ESI)**: $t_R$ = 6.11 min; *m/z* = 258.07 $[M + H]^+$. **$^1$H NMR** (400 MHz, MeOD): $\delta$ = 7.05 (d, *J* = 8.5 Hz, 2H), 6.75 (d, *J* = 8.5 Hz, 2H), 3.81 (s, 2H), 3.40 (t, *J* = 7.3 Hz, 2H), 2.73 (t, *J* = 7.3 Hz, 2H). **$^{13}$C NMR** (101 MHz, MeOD): $\delta$ = 169.32, 156.94, 130.74, 116.24, 42.81, 35.34, 28.78. **HRMS (ESI)**: *m/z* = 258.0124 calcd. for $[C_{10}H_{12}BrNO_2 + H]^+$; found: 258.0122.

##### Synthesis of dimethyl 2-((4-hydroxyphenethylcarbamoyl)methoxy) phenylphosphonate (**S-12**)

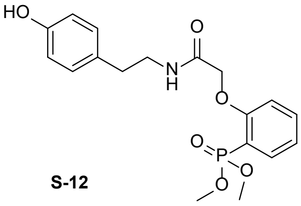

S-12

The phenylphosphonic acid derivative **S-12** was synthesized via GP **G** using phosphonate **S-2** (185 mg, 0.72 mmol), potassium carbonate (0.2 g, 1.44 mmol) and **S-11** (185 mg, 0.79 mmol) in acetone (20 mL). The crude product was purified by HPLC to obtain **S-12** (186 mg, 0.49 mmol, 69 %) as a white solid. **LC-MS (ESI)**: $t_R$ = 6.61 min; *m/z* = 380.00 $[M + H]^+$. **$^1$H NMR** (400 MHz, $CDCl_3$): $\delta$ = 8.43 (s, 1H), 7.72 – 7.66 (m, 1H), 7.59 – 7.49 (m, 2H), 7.12 (s, 1H), 6.98 – 6.92 (m, 3H), 6.74 (d, *J* = 7.1 Hz, 2H), 4.65 (s, 2H), 3.77 (d, *J* = 7.1 Hz, 6H), 3.57 – 3.52 (m, 2H), 2.78 (t, *J* = 7.2 Hz, 2H). **$^{13}$C NMR** (101 MHz, $CDCl_3$): $\delta$ = 169.08, 159.23, 155.01, 135.62, 134.42, 130.05, 129.90, 122.23, 115.53, 113.63, 112.76, 67.30, 53.47, 41.31, 34.62. **HRMS (ESI)**: *m/z* = 380.1258 calcd. for $[C_{18}H_{22}NO_6P + H]^+$; found: 380.1248.

#### Synthesis of the derivatives 9 – 11

##### Synthesis of (2-(2-oxo-2-((4-(pyridin-2-ylmethoxy)phenethyl)amino)ethoxy)-phenyl)phosphonic acid (**9**)

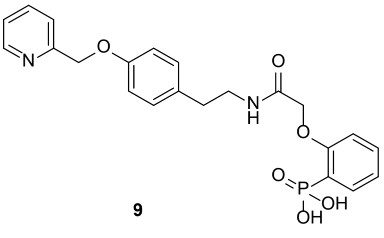

9

The phenylphosphonic acid derivative **9** was synthesized via GP **G** and GP **H**. GP **G** employed phosphonate **S-12** (0.74 g, 1.95 mmol), potassium carbonate (1.08 g, 7.78 mmol) and 2-(Bromomethyl)pyridine hydrobromide (0.54 g, 2.14 mmol) in DMF (60 mL). The crude intermediate was deprotected via GP **H** using TMSBr (1.29 mL, 9.75 mmol) in dry DCM (12.0 mL) and MeOH/$H_2O$ (3:1, 12.0 mL). The crude product was purified by HPLC to obtain **9** (0.86 g, 0.20 mmol, 10 % over two steps) as a white solid. **LC-MS (ESI)**: $t_R$ = 5.35 min; *m/z* = 443.14 $[M + H]^+$, calcd. for $C_{22}H_{23}N_2O_6P$: 442.13. **$^1$H NMR** (400 MHz, DMSO): $\delta$ = 9.11 (t, *J* = 5.6 Hz,

1H), 8.70 (d, $J$ = 5.2 Hz, 1H), 8.11 (t, $J$ = 7.8 Hz, 1H), 7.75 – 7.70 (m, 1H), 7.68 – 7.61 (m, 1H), 7.60 – 7.57 (m, 1H), 7.49 (t, $J$ = 8.0 Hz, 1H), 7.14 – 7.03 (m, 4H), 6.94 – 6.86 (m, 2H), 5.25 (s, 2H), 4.61 (s, 2H), 3.36 – 3.24 (m, 2H), 2.69 (t, $J$ = 7.6 Hz, 2H). **$^{13}$C NMR** (101 MHz, DMSO): $\delta$ = 168.09, 159.41, 156.46, 155.21, 146.88, 141.22, 133.46, 132.39, 129.97, 124.44, 123.36, 121.55, 114.93, 113.20, 68.78, 67.91, 40.52, 34.29. **HRMS (ESI)**: $m/z$ = 443.1367 calcd for $[C_{22}H_{23}N_2O_6P + H]^+$; found: 443.1360.

#### Synthesis of (2-(2-oxo-2-((4-(pyridin-3-ylmethoxy)phenethyl)amino)ethoxy)-phenyl)phosphonic acid (**10**)

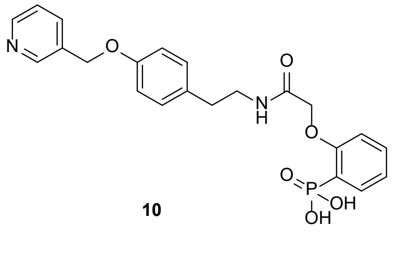

**10**

The phenylphosphonic acid derivative **10** was synthesized via GP **G** and GP **H**. GP **G** employed phosphonate **S-12** (650 mg, 1.70 mmol), potassium carbonate (940 mg, 6.81 mmol) and 3-(Bromomethyl)pyridine hydrobromide (470 mg, 1.87 mmol) in DMF (55 mL). The crude intermediate was deprotected via GP **H** using TMSBr (1.12 mL, 8.51 mmol) in dry DCM (10.0 mL) and MeOH/$H_2O$ (3:1, 10.0 mL). The crude product was purified by HPLC to obtain **10** (105 mg, 0.24 mmol, 14 % over two steps) as a white solid. **LC-MS (ESI)**: $t_R$ = 5.21 min; $m/z$ = 443.16 $[M + H]^+$, calcd. for $C_{22}H_{23}N_2O_6P$: 442.13. **$^1$H NMR** (400 MHz, DMSO): $\delta$ = 9.10 (m, 1H), 8.91 (s, 1H), 8.79 (d, $J$ = 5.4 Hz, 1H), 8.37 (d, $J$ = 8.0 Hz, 1H), 7.88 (dd, $J$ = 7.9, 5.6 Hz, 1H), 7.72 – 7.61 (m, 1H), 7.49 (t, $J$ = 7.9 Hz, 1H), 7.16 – 7.02 (m, 3H), 6.99 – 6.87 (m, 2H), 6.66 – 6.60 (m, 1H), 5.23 (s, 2H), 4.61 (s, 2H), 3.36 – 3.25 (m, 2H), 2.74 – 2.61 (m, 2H). **$^{13}$C NMR** (101 MHz, DMSO): $\delta$ = 167.93, 159.02, 156.24, 144.09, 143.50, 141.78, 135.93, 133.27, 132.24, 129.81, 129.52, 126.03, 123.14, 121.16, 117.60, 115.22, 114.78, 113.24, 67.73, 66.06, 40.35, 34.13. **HRMS (ESI)**: $m/z$ = 443.1367 calcd. for $[C_{22}H_{23}N_2O_6P + H]^+$; found: 443.1361.

#### Synthesis of (2-(2-oxo-2-((4-(pyridin-4-ylmethoxy)phenethyl)amino)ethoxy)-phenyl)phosphonic acid (**11**)

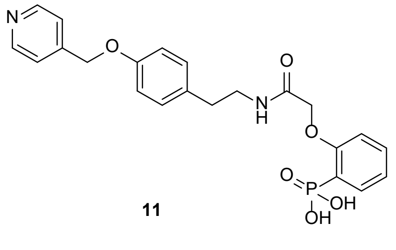

**11**

The phenylphosphonic acid derivative **11** was synthesized via GP **G** and GP **H**. GP **G** employed phosphonate **S-12** (0.77 g, 2.03 mmol), potassium carbonate (1.12 g, 8.12 mmol) and 4-(Bromomethyl)pyridine hydrobromide (0.56 g, 2.23 mmol) in DMF (65 mL). The crude intermediate was deprotected via GP **H** using TMSBr (1.34 mL, 10.15 mmol) in dry DCM (12 mL) and MeOH/$H_2O$ (3:1, 12 mL). The crude product was purified by HPLC to obtain **11** (81 mg, 0.18 mmol, 9 % over two steps) as a white solid. **LC-MS (ESI)**: $t_R$ = 4.88 min; $m/z$ = 443.12 $[M + H]^+$, calcd. for $C_{22}H_{23}N_2O_6P$: 442.13. **$^1$H NMR** (400 MHz, DMSO): $\delta$ = 9.12 (t, $J$ = 5.6 Hz, 1H), 8.83 (d, $J$ = 5.7 Hz, 2H), 7.90 (d, $J$ = 5.6 Hz, 2H), 7.70 – 7.59 (m, 1H), 7.49 (t, $J$ = 8.0 Hz, 1H), 7.16 – 7.02 (m, 4H), 6.94 – 6.86 (m, 2H), 5.35 (s, 2H), 4.60 (s, 2H), 3.30 (t, $J$ = 7.0 Hz, 2H), 2.69 (t, $J$ = 7.6 Hz, 2H). **$^{13}$C NMR** (101 MHz, DMSO): $\delta$ = 167.84, 158.95, 155.92, 154.66, 144.19, 133.18, 132.61, 132.32, 129.79, 123.58, 121.08, 114.71, 113.27, 67.72, 67.03, 40.22, 34.24. **HRMS (ESI)**: $m/z$ = 443.1367 calcd. for $[C_{22}H_{23}N_2O_6P + H]^+$; found: 443.1361.

### Synthesis of intermediates for the derivatives 12 – 14

#### Synthesis of 2-bromo-N-phenethyl-2-phenylacetamide (**S-13**)

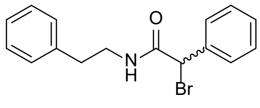

**S-13**

The acetamide **S-13** was synthesized via GP **E**. The acid chloride was generated using the 2-bromophenylacetic acid (11.08 g, 5.00 mmol), oxaly chloride (2 M, 3.00 mL, 6.00 mmol), dry DMF (3 drops) in dry DCM (2.5 mL). In addition phenylethylamine (0.63 mL, 5.00 mmol) and triethylamine (0.76 mL, 5.50 mmol) in dry DCM (7.5 mL + 15 mL) were used. The crude product was purified by column chromatography (cyclohexane/ethyl acetate 4:1) to obtain **S-13** (0.73 g, 2.31 mmol, 46 %) as a white solid. **TLC** (cyclohexane:ethyl acetate, 4:1 v/v): $R_f$ = 0.33. **LC-MS (ESI)**: $t_R$ = 9.59 min; $m/z$ = 318.14 $[M + H]^+$, calcd. for $C_{16}H_{16}BrNO$: 317.04. **$^1$H NMR** (400 MHz, $CDCl_3$): $\delta$ = 7.30 – 7.11 (m, 7H), 7.18 – 7.11 (m, 1H), 7.09 (d, $J$ = 7.1 Hz, 1H), 6.57 (t, $J$ = 6.1 Hz, 1H), 5.29 (d, $J$ = 21.5 Hz, 1H), 3.52 – 3.44 (m, 2H), 2.82 – 2.70 (m, 2H).

**$^{13}$C NMR** (101 MHz, $CDCl_3$): $\delta$ = 167.08, 138.46, 137.44, 129.13, 129.02, 128.96, 128.91, 128.81, 128.39, 127.83, 126.77, 51.63, 41.59, 35.44.

#### Synthesis of 2-bromo-N-phenethylpropanamid (**S-14**)

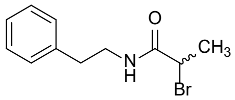

**S-14**

The bromoacetamide **S-14** was synthesized via GP **D** phenethylamine (0.63 mL, 5.00 mmol), triethylamine (0.76 mL, 5.50 mmol) and 2-bromopropionyl bromide (0.52 mL, 5.00 mmol) in dry DCM (25 mL). The crude product was purified by column chromatography (cyclohexane/ethyl acetate 4:1) to obtain **S-14** (1.03 g, 4.02 mmol, 80 %) as a white solid. **TLC** (cyclohexane:ethyl acetate, 4:1 v/v): $\mathbf{R_f}$ = 0.25. **LC-MS (ESI)**: $t_R$ = 8.33 min; *m/z* = 256.13 $[M + H]^+$, calcd. for $C_{11}H_{14}BrNO$: 255.03. **$^1$H NMR** (400 MHz, $CDCl_3$): $\delta$ = 7.28 – 7.19 (m, 2H), 7.20 – 7.08 (m, 3H), 6.41 (s, 1H), 4.26 (q, *J* = 7.0 Hz, 1H), 3.50 – 3.38 (m, 2H), 2.75 (t, *J* = 7.0 Hz, 2H), 1.74 (d, *J* = 7.1 Hz, 3H). **$^{13}$C NMR** (101 MHz, $CDCl_3$): $\delta$ = 169.33, 138.55, 128.88, 126.72, 45.30, 41.38, 35.48, 23.23.

#### Synthesis of 2-bromo-2-fluoro-N-phenethylacetamide (**S-15**)

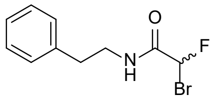

**S-15**

Erbium(III) trifluoromethanesulfonate (308 mg, 0.50 mmol) was dissolved in dry acetonitrile (10 mL). Ethyl bromofluoroacetate (1.18 mL, 10.00 mmol) and phenethylamine (1.51 mL, 12.00 mmol) were added and the resulting suspension was stirred for 12 h at 50 °C. After cooling down to room temperature the reaction mixture was poured into $H_2O$ and extracted with DCM. The combined organic phases dried over $MgSO_4$ and the solvent was removed under reduced pressure. The crude product was purified by column chromatography (cyclohexane/ethyl acetate 4:1) to obtain **S-15** (338 mg, 1.30 mmol, 13 %) as a slightly yellow solid. **TLC** (cyclohexane:ethyl acetate, 4:1 v/v): $\mathbf{R_f}$ = 0.52. **LC-MS (ESI)**: $t_R$ = 8.29 min; *m/z* = 259.87 $[M + H]^+$, calcd. for $C_{10}H_{11}BrFNO$: 259.00. **$^1$H NMR** (400 MHz, $CDCl_3$): $\delta$ = 7.40 – 7.33 (m, 2H), 7.30 – 7.21 (m, 3H), 6.59 (d, *J* = 50.9 Hz, 1H), 6.41 (s, 1H), 3.71 – 3.56 (m, 2H), 2.91 (t, *J* = 7.0 Hz, 2H). **$^{13}$C NMR** (101 MHz, $CDCl_3$): $\delta$ = 164.41, 138.09, 128.91, 126.94, 86.00, 40.93, 35.39.

### Synthesis of the derivatives 12 – 14

#### Synthesis of (2-(2-oxo-2-(phenethylamino)-1-phenylethoxy)phenyl)phosphonic acid (**12**)

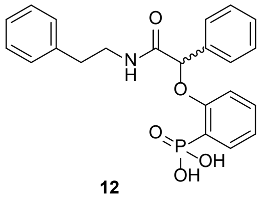

**12**

The phenylphosphonic acid derivative **12** was synthesized via GP **G** and GP **H**. GP **G** employed the phosphonate **S-2** (202 mg, 1.00 mol), potassium carbonate (276 mg, 2.00 mmol) and **S-13** (318 mg, 1.00 mmol) in acetone (15 mL). The crude intermediate was deprotected via GP **H** using TMSBr (1.32 mL, 10.00 mmol) in dry DCM (8 mL) and $MeOH/H_2O$ (3:1, 6 mL). The crude product was purified by HPLC to obtain **12** (167 mg, 0.41 mmol, 41 % over two steps) as a white solid. **LC-MS (ESI)**: $t_R$ = 7.03 min; *m/z* = 411.92 $[M + H]^+$, calcd. for $C_{22}H_{22}NO_5P$: 411.12. **$^1$H NMR** (400 MHz, $CDCl_3$): $\delta$ = 9.40 (s, 1H), 7.69 – 7.63 (m, 1H), 7.58 (d, *J* = 7.2 Hz, 1H), 7.43 – 7.29 (m, 4H), 7.22 – 7.10 (m, 5H), 7.07 – 7.00 (m, 2H), 5.90 (s, 1H), 3.20 (m, 2H), 2.73 (t, *J* = 7.5 Hz, 2H). **$^{13}$C NMR** (101 MHz, $CDCl_3$): $\delta$ = 168.77, 157.66, 139.23, 137.08, 132.96, 128.58, 128.17, 127.12, 125.99, 123.13, 121.34, 120.96, 113.24, 78.82, 66.95, 34.77, 31.31. **HRMS (ESI)**: *m/z* = 412.1309 calcd. for $[C_{22}H_{22}NO_5P + H]^+$; found: 412.1304.

#### Synthesis of (2-(2-Oxo-2-((2-phenylpropyl)amino)ethoxy)phenyl)phosphonic acid (**13**)

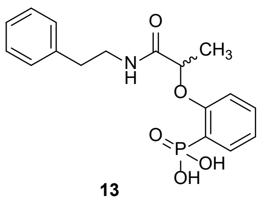

**13**

The phenylphosphonic acid derivative **13** was synthesized via GP **G** and GP **H**. GP **G** employed the phosphonate **S-2** (202 mg, 1.00 mol), potassium carbonate (276 mg, 2.00 mmol) and **S-14** (256 mg, 1.00 mmol) in acetone (15 mL). The crude intermediate was deprotected via GP **H** using TMSBr (1.32 mL, 10.00 mmol) in dry DCM (8 mL) and $MeOH/H_2O$ (3:1, 6 mL). The crude product was purified by HPLC to obtain **13** (47 mg, 0.13 mmol, 13 % over two steps) as a white solid. **LC-MS (ESI)**: $t_R$ = 5.97 min; *m/z* = 350.03 $[M + H]^+$, calcd. for $C_{17}H_{20}NO_5P$: 349.11. **$^1$H NMR** (400 MHz, DMSO): $\delta$ = 9.31 (t, *J* = 5.6 Hz, 1H), 7.67 – 7.61 (m, 1H), 7.48 (t, *J* = 7.3 Hz, 1H), 7.24 – 7.19 (m, 2H), 7.18 – 7.12 (m, 4H), 7.09 – 7.00 (m, 1H), 4.94 (q, *J* = 6.6 Hz, 1H), 3.31 – 3.24 (m, 2H), 2.71 (t,

$J$ = 7.6 Hz, 2H), 1.47 (d, $J$ = 6.7 Hz, 3H). **$^{13}$C NMR** (101 MHz, DMSO): $\delta$ = 170.94, 158.46, 139.32, 132.78, 128.55, 128.21, 125.98, 120.75, 112.92, 74.32, 34.86, 31.31, 18.45. **HRMS (ESI)**: *m/z* = 350.1152 calcd. for $[C_{17}H_{20}NO_5P + H]^+$; found: 350.1151.

##### Synthesis of (2-(1-fluoro-2-oxo-2-(phenethylamino)ethoxy)phenyl)phosphonic acid (**14**)

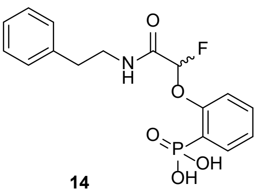

**14**

The phenylphosphonic acid derivative **43** was synthesized via GP **G** and GP **H**. GP **G** employed phosphonate **S-2** (202 mg, 1.00 mmol), potassium carbonate (276 mg, 2.00 mmol) and **S-15** (260 mg, 1.00 mmol) in DMF (6 mL). The crude intermediate was deprotected via GP **H** using TMSBr (1.32 mL, 10.00 mmol) in dry DCM (8 mL) and MeOH/$H_2O$ (3:1, 6 mL). The crude product was purified by HPLC to obtain **14** (170 mg, 0.48 mmol, 48 % over two steps) as a white solid. **LC-MS (ESI)**: $t_R$ = 5.92 min; *m/z* = 353.95 $[M + H]^+$, calcd. for $C_{16}H_{17}FNO_5P$: 353.08. **$^1$H NMR** (400 MHz, DMSO): $\delta$ = 9.64 (t, $J$ = 5.5 Hz, 1H), 7.01 (ddd, $J$ = 13.9, 7.6, 1.8 Hz, 1H), 7.60 (td, $J$ = 7.8, 1.7 Hz, 1H), 7.34 – 7.19 (m, 7H), 6.16 (d, $J$ = 58.7 Hz, 1H), 3.49 – 3.30 (m, 2H), 2.82 (tt, $J$ = 8.0, 3.7 Hz, 2H). **$^{13}$C NMR** (101 MHz, DMSO): $\delta$ = 163.51, 156.86, 139.16, 132.81, 128.41, 126.21, 123.75, 115.97, 104.75, 102.51, 99.56, 40.60, 34.64. **HRMS (ESI)**: *m/z* = 354.0901 calcd. for $[C_{16}H_{17}FNO_5P + H]^+$; found: 354.0898.

#### Synthesis of intermediate for the derivatives 15 – 16

##### Synthesis of 2-bromo-N-phenethylacetamide (**S-16**)

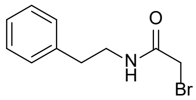

**S-16**

The bromoacetamide **S-16** was synthesized via GP **D** using phenethylamine (0.63 mL, 5.00 mmol), triethylamine (0.76 mL, 5.50 mmol) and bromoacatyl bromide (0.44 mL, 5.00 mmol), in dry DCM (25 mL). The crude product was purified by HPLC to obtain **S-16** (0.96 g, 3.99 mmol, 80 %) as a white solid. **LC-MS (ESI)**: $t_R$ = 7.75 min; *m/z* = 242.05 $[M + H]^+$. **$^1$H NMR** (400 MHz, $CDCl_3$): $\delta$ = 7.28 - 7.24 (m, 2H), 7.20 – 7.14 (m, 3H), 6.89 (s, 1H), 3.73 (s, 2H), 3.49 – 3.44 (m, 2H), 2.79 (t, $J$ = 7.0 Hz, 2H). **$^{13}$C NMR** (101 MHz, $CDCl_3$): $\delta$ = 165.79, 138.36, 128.66, 128.56, 126.53, 41.29, 35.24, 29.02. **HRMS (ESI)**: *m/z* = 242.0175 calcd. for $[C_{10}H_{12}BrNO + H]^+$; found: 242.017.

#### Synthesis of the derivatives 15 – 16

##### Synthesis of (3-(2-oxo-2-(phenethylamino)ethoxy)phenyl)phosphonic acid (**15**)

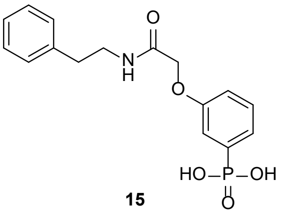

**15**

The phenylphosphonic acid derivative **15** was synthesized via GP **G** and GP **H**. GP **G** employed diethyl (3-hydroxyphenyl)phosphonate (230 mg, 1.00 mol), potassium carbonate (276 mg, 2.00 mmol) and **S-16** (267 mg, 1.10 mmol) in acetone (15 mL). The crude intermediate was deprotected via GP **H** using TMSBr (6.6 mL, 50.00 mmol) in dry DCM (8 mL) and MeOH/$H_2O$ (3:1, 8 mL). The crude product was purified by HPLC to obtain **15** (207 mg, 0.62 mmol, 62 % over two steps) as a white solid. **LC-MS (ESI)**: $t_R$ = 5.86 min; *m/z* = 336.14 $[M + H]^+$, calcd. for $C_{16}H_{18}NO_5P$: 335.09. **$^1$H NMR** (400 MHz, DMSO): $\delta$ = 8.23 (t, $J$ = 5.8 Hz, 1H), 7.41 – 7.34 (m, 1H), 7.31 – 7.23 (m, 4H), 7.22 – 7.17 (m, 3H), 7.08 – 7.01 (m, 1H), 4.48 (s, 2H), 3.37 (dt, $J$ = 8.2, 6.2 Hz, 2H), 2.76 (t, $J$ = 7.5 Hz, 2H). **$^{13}$C NMR** (101 MHz, DMSO): $\delta$ = 167.34, 157.16, 139.31, 136.73, 134.94, 129.50, 128.37, 126.12, 123.36, 117.18, 116.69, 66.98, 35.15. **HRMS (ESI)**: *m/z* = 336.0996 calcd. for $[C_{16}H_{18}NO_5P + H]^+$; found: 336.0991.

##### Synthesis of (3-(2-oxo-2-(phenethylamino)ethoxy)phenyl)phosphonic acid (**16**)

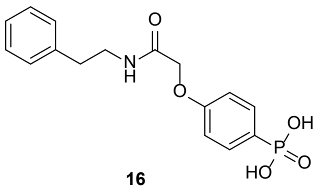

16

The phenylphosphonic acid derivative **16** was synthesized via GP **G** and GP **H**. GP **G** employed diethyl (4-hydroxyphenyl)phosphonate (230 mg, 1.00 mol), potassium carbonate (276 mg, 2.00 mmol) and **S-16** (267 mg, 1.10 mmol) in acetone (15 mL). The crude intermediate was deprotected via GP **H** using TMSBr (6.6 mL, 50.00 mmol) in dry DCM (8 mL) and MeOH/$H_2O$ (3:1, 8 mL). The crude product was purified by HPLC to obtain **16** (245 mg, 0.73 mmol, 73 % over two steps) as a white solid. **LC-MS (ESI)**: $t_R$ = 5.87 min; *m/z* = 336.10 [M + H]$^+$, calcd. for $C_{16}H_{18}NO_5P$: 335.09. **$^1$H NMR** (400 MHz, DMSO): $\delta$ = 8.20 (t, *J* = 5.8 Hz, 1H), 7.62 (dd, *J* = 12.5, 8.6 Hz, 2H), 7.28 (t, *J* = 7.5 Hz, 2H), 7.21 – 7.18 (m, 3H), 7.00 (dd, *J* = 8.7, 2.8 Hz, 2H), 4.52 (s, 2H), 3.38 (dt, *J* = 8.0, 6.2 Hz, 2H), 2.76 (t, *J* = 7.4 Hz, 2H). **$^{13}$C NMR** (101 MHz, DMSO): $\delta$ = 167.25, 159.68, 139.29, 132.40, 128.64, 128.37, 127.22, 126.14, 125.36, 114.24, 66.86, 35.11. **HRMS (ESI)**: *m/z* = 336.0996 calcd. for [$C_{16}H_{18}NO_5P$ + H]$^+$; found: 336.0993.

#### Synthesis of intermediates for the derivatives 17 – 19

##### Synthesis of 2-hydroxyphenyl dimethyl phosphate (**S-17**)

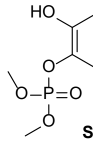

S-17

The phosphate **S-17** was synthesized via GP **A** using catechol (5.51 g, 50.0 mmol), tetrachlormethane (6.90 mL, 50.0 mmol), triethylamine (7.45 mL, 50.0 mmol) and dimethyl phosphite (4.80 mL, 15.0 mmol) in dry DCM (50 mL). The crude product was purified by column chromatography (ethyl acetate) to obtain **S-17** (7.15 g, 32.8 mmol, 66 %) as slightly brown solid. **TLC** (ethyl acetate, v/v): $R_f$ = 0.5. **LC-MS (ESI)**: $t_R$ = 6.16 min; *m/z* = 219.05 [M + H]$^+$, calcd. for $C_8H_{11}O_5P$: 218.03. **$^1$H NMR** (400 MHz, MeOD): $\delta$ = 7.12 (dt, *J* = 8.1, 1.5 Hz, 1H), 7.06 – 6.96 (m, 1H), 6.97 – 6.84 (m, 1H), 6.77 (td, *J* = 7.7, 1.6 Hz, 1H), 3.84 (d, *J* = 11.5 Hz, 6H). **$^{13}$C NMR** (101 MHz, MeOD): $\delta$ = 149.66, 139.86, 127.26, 122.29, 120.70, 118.23, 55.82.

##### Synthesis of 3-hydroxyphenyl dimethyl phosphate (**S-18**)

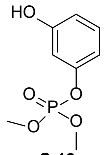

S-18

The phosphate **S-18** was synthesized via GP **A** using resorcinol (5.51 g, 50.0 mmol), tetrachlormethane (6.90 mL, 50.0 mmol), triethylamine (7.45 mL, 50.0 mmol) and dimethyl phosphite (4.80 mL, 15.0 mmol) in dry DCM (50 mL). The crude product was purified by column chromatography (ethyl acetate) to obtain **S-18** (1.01 g, 4.6 mmol, 9 %) as slightly yellow solid. **TLC** (ethyl acetate, v/v): $R_f$ = 0.5. **LC-MS (ESI)**: $t_R$ = 6.16 min; *m/z* = 219.12 [M + H]$^+$, calcd. for $C_8H_{11}O_5P$: 218.03. **$^1$H NMR** (400 MHz, MeOD): $\delta$ = 7.25 – 7.16 (m, 1H), 6.75 – 6.67 (m, 3H), 3.87 (d, *J* = 11.6 Hz, 16H). **$^{13}$C NMR** (101 MHz, MeOD): $\delta$ = 159.98, 152.52, 131.33, 113.60, 111.64, 108.20, 55.76.

##### Synthesis of 4-hydroxyphenyl dimethyl phosphate (**S-19**)

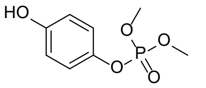

S-19

The phosphate **S-19** was synthesized via GP **A** using hydroquinone (1.10 g, 10.0 mmol), tetrachlormethane (0.96 mL, 10.0 mmol), triethylamine (1.49 mL, 10.0 mmol) and dimethyl phosphite (1.38 mL, 15.0 mmol) in dry DCM (10 mL). The crude product was purified by column chromatography (ethyl acetate) to obtain **S-19** (0.32 g, 1.5 mmol, 15 %) as slightly yellow solid. **TLC** (ethyl acetate, v/v): $R_f$ = 0.5. **LC-MS (ESI)**: $t_R$ = 5.91 min; *m/z* = 219.10 [M + H]$^+$, calcd. for $C_8H_{11}O_5P$: 218.03. **$^1$H NMR** (400 MHz, MeOD): $\delta$ = 7.07 (d, *J* = 8.5 Hz, 2H), 6.81 (dd, *J* = 9.7, 3.1 Hz, 2H), 3.89 (d, *J* = 11.7 Hz, 6H). **$^{13}$C NMR** (101 MHz, MeOD): $\delta$ = 156.11, 144.31, 121.85, 116.99, 55.70.

#### Synthesis of the derivatives 17 – 19

##### Synthesis of 2-(2-oxo-2-(phenethylamino)ethoxy)phenyl dihydrogen phosphate (**17**)

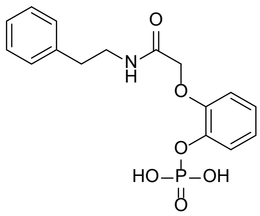


17

The dihydrogen phosphate derivative **17** was synthesized via GP **G** and GP **H**. GP **G** employed phosphate **S-17** (436 mg, 2.00 mol), potassium carbonate (553 mg, 4.00 mmol) and **S-16** (533 mg, 2.20 mmol) in acetone (30 mL). The crude intermediate was deprotected via GP **H** using TMSBr (2.64 mL, 20.00 mmol) in dry DCM (8 mL) and MeOH/$H_2O$ (3:1, 8 mL). The crude product was purified by HPLC to obtain **17** (429 mg, 1.22 mmol, 61 % over two steps) as a white solid. **LC-MS (ESI)**: $t_R$ = 5.58 min; *m/z* = 352.06 [M + H]$^+$, calcd. for $C_{16}H_{18}NO_6P$: 351.09 . **$^1$H NMR** (400 MHz, DMSO): *δ* = 8.18 (t, *J* = 5.8 Hz, 1H), 7.33 – 7.24 (m, 3H). 7.23 – 7.18 (m, 3H), 6.84 – 6.79 (m, 2H), 6.75 – 6.67 (m, 1H), 4.44 (s, 2H), 3.37 (dt, *J* = 8.0, 6.2 Hz, 2H), 2.76 (t, *J* = 7.5 Hz, 2H). **$^{13}$C NMR** (101 MHz, DMSO): *δ* = 167.31, 158.55, 152.43, 139.31, 129.94, 128.37, 126.13, 112.96, 109.92, 107.38, 67.12, 35.14. **HRMS (ESI)**: *m/z* = 352.0945 calcd. for [$C_{16}H_{18}NO_6P$ + H]$^+$; found: 352.0944.

#### Synthesis of 3-(2-oxo-2-(phenethylamino)ethoxy)phenyl dihydrogen phosphate (**18**)

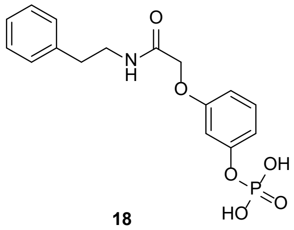


18

The dihydrogen phosphate derivative **18** was synthesized via GP **G** and GP **H**. GP **G** employed phosphate **S-18** (436 mg, 2.00 mol), potassium carbonate (553 mg, 4.00 mmol) and **S-16** (533 mg, 2.20 mmol) in acetone (30 mL). The crude intermediate was deprotected via GP **H** using TMSBr (2.64 mL, 20.00 mmol) in dry DCM (16 mL) and MeOH/$H_2O$ (3:1, 16 mL). The crude product was purified by HPLC to obtain **18** (150 mg, 0.43 mmol, 21 % over two steps) as a white solid. **LC-MS (ESI)**: $t_R$ = 5.68 min; *m/z* = 352.09 [M + H]$^+$, calcd. for $C_{16}H_{18}NO_6P$: 351.09 . **$^1$H NMR** (400 MHz, DMSO): *δ* = 8.69 (t, *J* = 5.7 Hz, 1H), 7.25 (t, *J* = 7.2 Hz, 2H), 7.22 – 7.13 (m, 4H), 7.15 – 7.06 (m, 1H), 7.05 – 6.91 (m, 2H), 4.55 (s, 2H), 3.33 (dt, *J* = 8.3, 6.1 Hz, 2H), 2.72 (t, *J* = 7.6 Hz, 2H). **$^{13}$C NMR** (101 MHz, DMSO): *δ* = 167.58, 148.99, 140.33, 139.28, 128.32, 126.06, 125.21, 122.22, 121.35, 113.44, 66.78, 34.90. **HRMS (ESI)**: *m/z* = 352.0945 calcd. for [$C_{16}H_{18}NO_6P$ + H]$^+$; found: 352.0943.

#### Synthesis of 4-(2-oxo-2-(phenethylamino)ethoxy)phenyl dihydrogen phosphate (**19**)

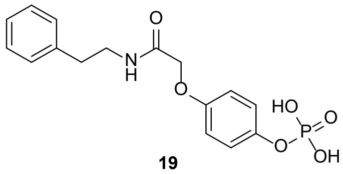


19

The dihydrogen phosphate derivative **19** was synthesized via GP **G** and GP **H**. GP **G** employed phosphate **S-19** (218 mg, 1.00 mol), potassium carbonate (276 mg, 2.00 mmol) and **S-16** (267 mg, 1.10 mmol) in acetone (15 mL). The crude intermediate was deprotected via GP **H** using TMSBr (1.32 mL, 10.00 mmol) in dry DCM (8 mL) and MeOH/$H_2O$ (3:1, 8 mL). The crude product was purified by HPLC to obtain **19** (241 mg, 0.69 mmol, 69 % over two steps) as a white solid. **LC-MS (ESI)**: $t_R$ = 5.52 min; *m/z* = 352.01 [M + H]$^+$, calcd. for $C_{16}H_{18}NO_6P$: 351.09 . **$^1$H NMR** (400 MHz, DMSO): *δ* = 8.13 (t, *J* = 5.8 Hz, 1H), 7.33 – 7.24 (m, 2H), 7.22 – 7.17 (m, 2H), 7.10 (d, *J* = 8.9 Hz, 2H), 6.91 (d, *J* = 9.0 Hz, 2H), 4.42 (s, 2H), 3.37 (dt, *J* = 7.9, 6.2 Hz, 2H), 2.76 (t, *J* = 7.4 Hz, 2H). **$^{13}$C NMR** (101 MHz, DMSO): *δ* = 167.56, 154.08, 145.62, 139.30, 128.64, 128.36, 126.13, 121.02, 115.41, 67.47, 35.12. **HRMS (ESI)**: *m/z* = 352.0945 calcd. for [$C_{16}H_{18}NO_6P$ + H]$^+$; found: 352.0943.

### Synthesis of intermediates for the derivatives 20 – 22

#### Synthesis of Diethyl (2-hydroxybenzyl)phosphonate (**S-20**)

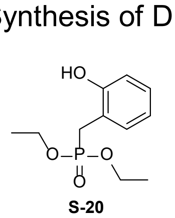


S-20

2-Hydroxybenzyl alcohol (1.24 g, 10.0 mmol) was dissolved in dry *o*-xylene (10mL) and heated up to 80 °C. Triethyl phosphite (2.07 mL, 12.0 mmol) was added and the reaction mixture was stirred for 1 h at 80 °C. After cooling down to room temperature the volatiles were removed under reduced pressure. The crude product was purified by column chromatography (cyclohexane/ethyl acetate 1:1) to obtain **S-20** (2.28 g, 9.3 mmol, 93 %) as a slightly yellow oil. **TLC** (cyclohexane:ethyl acetate, 1:1 v/v): **$R_f$** = 0.2. **LC-MS (ESI)**: $t_R$ = 6.96 min; *m/z* = 245.01 [M + H]$^+$, calcd. for $C_{11}H_{17}O_4P$: 244.09. **$^1$H NMR** (400 MHz, MeOD): *δ* = 7.22 (d, *J* = 7.6 Hz, 1H), 7.08 (t, *J* = 7.7 Hz, 1H), 6.88 – 6.75 (m, 2H), 4.03 (p, *J* = 7.2 Hz, 4H), 3.23 (d, *J* = 21.5 Hz, 2H), 1.24 (t, *J* = 7.3 Hz, 6H). **$^{13}$C NMR** (101 MHz, MeOD): *δ* = 156.43, 132.29, 129.20, 120.40, 116.12, 63.43, 26.26, 16.60.

#### Synthesis of Diethyl (3-hydroxybenzyl)phosphonate (**S-21**)

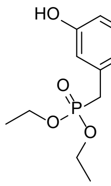

S-21

3-Hydroxybenzyl alcohol (1.24 g, 10.0 mmol) and $ZnBr_2$ (2.48 g, 11.00 mmol) were dissolved in dry DCM (30mL) and cooled down to 0 °C. Triethyl phosphite (8.65 mL, 50.00 mmol) was added and the reaction mixture was stirred for 16 h, allowing the mixture to slowly reach room temperature. The reaction mixture was poured into ice/HCl and extracted with DCM. The combined organic phases were dried over $MgSO_4$ and the solvent was removed under reduced pressure. The crude product was purified by column chromatography (ethyl acetate) to obtain **S-21** (1.29 g, 5.26 mmol, 53 %) as a yellow oil. **TLC** (ethyl acetate, v/v): $\mathbf{R_f}$ = 0.56. **LC-MS (ESI)**: $t_R$ = 6.66 min; *m/z* = 244.98 $[M + H]^+$, calcd. for $C_{11}H_{17}O_4P$: 244.09 . $^1$**H NMR** (400 MHz, $CDCl_3$): $\delta$ = 7.07 (t, *J* = 7.8 Hz, 1H), 6.89 (s, 1H), 6.71 (t, *J* = 7.7 Hz, 2H), 4.08 – 3.87 (m, 4H), 3.05 (d, *J* = 21.6 Hz, 2H), 1.19 (t, *J* = 7.2 Hz, 6H). $^{13}$**C NMR** (101 MHz, $CDCl_3$): $\delta$ = 157.33, 132.03, 129.48, 120.91, 116.83, 114.38, 62.63, 34.09, 16.20.

#### Synthesis of Diethyl (4-hydroxybenzyl)phosphonate (**S-22**)

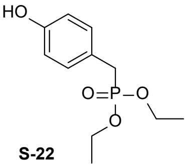

S-22

4-Hydroxybenzyl alcohol (1.24 g, 10.0 mmol) and $ZnBr_2$ (2.48 g, 11.00 mmol) were dissolved in dry DCM (30mL) and cooled down to 0 °C. Triethyl phosphite (8.65 mL, 50.00 mmol) was added and the reaction mixture was stirred for 16 h, allowing the mixture to slowly reach room temperature. The reaction mixture was poured into ice/HCl and extracted with DCM. The combined organic phases were dried over $MgSO_4$ and the solvent was removed under reduced pressure. The crude product was purified by column chromatography (ethyl acetate) to obtain **S-22** (1.10 g, 4.50 mmol, 45 %) as a yellow oil. **TLC** (ethyl acetate, v/v): $\mathbf{R_f}$ = 0.1 . **LC-MS (ESI)**: $t_R$ = 6.40 min; *m/z* = 244.95 $[M + H]^+$, calcd. for $C_{11}H_{17}O_4P$: 244.09 . $^1$**H NMR** (400 MHz, $CDCl_3$): $\delta$ = 7.04 (dd, *J* = 8.6, 2.8 Hz, 2H), 6.73 (d, *J* = 8.3 Hz, 2H), 4.07 – 3.89 (m, 4H), 3.08 (d, *J* = 20.9 Hz, 2H), 1.21 (t, *J* = 7.2 Hz, 6H). $^{13}$**C NMR** (101 MHz, $CDCl_3$): $\delta$ = 156.21, 130.58, 120.97, 115.80, 62.47, 33.18, 16.28.

### Synthesis of the derivatives 20 – 22

#### Synthesis of (2-(2-oxo-2-(phenethylamino)ethoxy)benzyl)phosphonic acid (**20**)

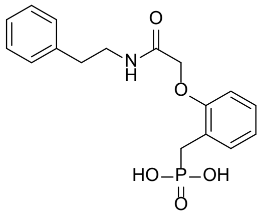

20

The phenylphosphonic acid derivative **20** was synthesized via GP **G** and GP **H**. GP **G** employed phosphonate **S-20** (181 mg, 0.74 mmol), potassium carbonate (204 mg, 1.48 mmol) and **S-16** (196 mg, 0.81 mmol) in acetone (11 mL). The crude intermediate was deprotected via GP **H** using TMSBr (5.30 mL, 37.00 mmol) in dry DCM (5.7 mL) and MeOH/$H_2O$ (3:1, 6 mL). The crude product was purified by HPLC to obtain **20** (181 mg, 0.52 mmol, 70 % over two steps) as a white solid. **LC-MS (ESI)**: $t_R$ = 6.34 min; *m/z* = 350.11 $[M + H]^+$, calcd. for $C_{17}H_{20}NO_5P$: 349.11 . $^1$**H NMR** (400 MHz, DMSO): $\delta$ = 9.03 (t, *J* = 5.6 Hz, 1H), 7.25 (dd, *J* = 7.9, 6.6 Hz, 2H), 7.23 – 7.12 (m, 4H), 6.91 (t, *J* = 7.4 Hz, 1H), 6.86 (d, *J* = 8.2 Hz, 1H), 4.52 (s, 2H), 3.33 – 3.23 (m, 2H), 3.09 (d, *J* = 21.4 Hz, 2H), 2.70 (dd, *J* = 8.8, 6.5 Hz, 2H). $^{13}$**C NMR** (101 MHz, DMSO): $\delta$ = 167.81, 155.03, 139.40, 131.51, 128.53, 127.70, 126.01, 122.24, 120.79, 110.48, 66.23, 34.88, 29.39. **HRMS (ESI)**: *m/z* = 350.1152 calcd. for $[C_{17}H_{20}NO_5P + H]^+$; found: 350.1151.

#### Synthesis of (3-(2-oxo-2-(phenethylamino)ethoxy)benzyl)phosphonic acid (**21**)

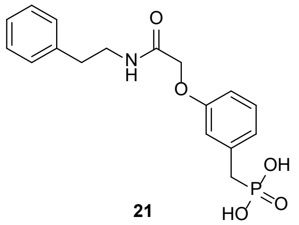

21

The phenylphosphonic acid derivative **21** was synthesized via GP **G** and GP **H**. GP **G** employed phosphonate **S-21** (488 mg, 2.00 mmol), potassium carbonate (553 mg, 4.00 mmol) and **S-16** (484 mg, 2.00 mmol) in acetone (30 mL). The crude intermediate was deprotected via GP **H** using TMSBr (10.00 mL, 76.00 mmol) in dry DCM (16 mL) and MeOH/$H_2O$ (3:1, 12 mL). The crude product was purified by HPLC to obtain **21** (137 mg, 0.39 mmol, 20 % over two steps) as a white solid. **LC-MS (ESI)**: $t_R$ = 6.25 min; *m/z* = 350.17 $[M + H]^+$, calcd. for $C_{17}H_{20}NO_5P$: 349.11 . $^1$**H NMR** (400 MHz, DMSO): $\delta$ = 8.14 (t, *J* = 5.8 Hz, 1H), 7.33 – 7.25 (m, 2H), 7.24 – 7.15 (m, 4H), 6.90 – 6.85 (m, 2H), 6.77 (d, *J* = 8.8 Hz, 1H), 4.41 (s, 2H), 3.37 (dt, *J* = 8.1, 6.2 Hz, 2H), 2.96 (d, *J* = 21.4 Hz, 2H), 2.76 (t, *J* = 7.5 Hz, 2H). $^{13}$**C NMR** (101 MHz, DMSO): $\delta$ = 167.51, 157.49, 139.32, 135.79, 128.98, 128.61, 128.35, 126.11, 122.83,

116.65, 111.86, 67.01, 36.04, 35.12. **HRMS (ESI)**: *m/z* = 350.1152 calcd. for $[C_{17}H_{20}NO_5P + H]^+$; found: 350.1150.

#### Synthesis of (4-(2-oxo-2-(phenethylamino)ethoxy)benzyl)phosphonic acid (**22**)

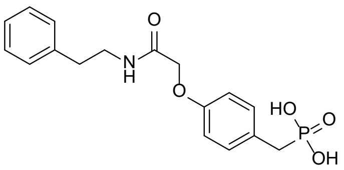

**22**

The phenylphosphonic acid derivative **22** was synthesized via GP **G** and GP **H**. GP **G** employed phosphonate **S-22** (293 mg, 1.20 mmol), potassium carbonate (332 mg, 2.40 mmol) and **S-16** (291 mg, 1.20 mmol) in acetone (18 mL). The crude intermediate was deprotected via GP **H** using TMSBr (6.00 mL, 45.50 mmol) in dry DCM (8 mL) and MeOH/$H_2O$ (3:1, 8 mL). The crude product was purified by HPLC to obtain **22** (93 mg, 0.27 mmol, 22 % over two steps) as a white solid. **LC-MS (ESI)**: $t_R$ = 6.22 min; *m/z* = 350.10 $[M + H]^+$, calcd. for $C_{17}H_{20}NO_5P$: 349.11 . **$^1$H NMR** (400 MHz, DMSO): $\delta$ = 8.13 (t, *J* = 5.8 Hz, 1H), 7.29 (t, *J* = 7.5 Hz, 2H), 7.21 – 7.15 (m, 5H), 6.86 (d, *J* = 8.4 Hz, 2H), 4.41 (s, 2H), 3.37 (dt, *J* = 8.1, 6.2 Hz, 2H), 2.92 (d, *J* = 21.0 Hz, 2H), 2.76 (t, *J* = 7.5 Hz, 2H). **$^{13}$C NMR** (101 MHz, DMSO): $\delta$ = 167.62, 156.11, 139.29, 130.61, 128.62, 126.84, 126.12, 114.32, 67.09, 35.11, 33.74. **HRMS (ESI)**: *m/z* = 350.1152 calcd. for $[C_{17}H_{20}NO_5P + H]^+$; found: 350.1150.

### Synthesis of intermediate for the derivative 23

#### Synthesis of 2-(4,4,5,5-Tetramethyl-1,3,2-dioxaborolan-2-yl)phenol (**S-23**)

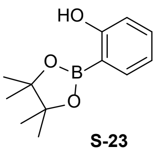

**S-23**

2-Hydroxybenzeneboronic acid (1.38 g, 10.0 mmol) and pinacol (1.30 g, 11.0 mmol) were dissolved in dry DCM (50 mL). The reaction mixture was stirred for 16 h at room temperature. The solvent was removed under reduced pressure to obtain **S-23** (2.07 g, 9.25 mmol, 93 %) as a slightly yellow oil. **LC-MS (ESI)**: $t_R$ = 10.81 min; *m/z* = 220.99 $[M + H]^+$, calcd. for $C_{12}H_{17}BO_3$: 220.13. **$^1$H NMR** (400 MHz, $CDCl_3$): $\delta$ = 7.84 (s, 1H), 7.65 (d, *J* = 7.3 Hz, 1H), 6.91 (t, *J* = 7.2 Hz, 2H), 1.39 (s, 12H). **$^{13}$C NMR** (101 MHz, $CDCl_3$): $\delta$ = 163.69, 135.76, 133.85, 119.54, 115.49, 84.47, 24.83.

### Synthesis of the derivative 23

#### Synthesis of (2-(2-Oxo-2-(phenethylamino)ethoxy)phenyl)boronic acid (**23**)

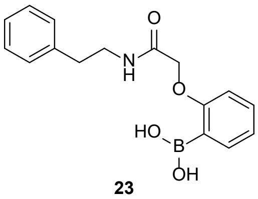

**23**

The boronic acid derivative **23** was synthesized via GP **G** using **S-23** (440 mg, 2.0 mmol), potassium carbonate (553 mg, 4.0 mmol) and **S-16** (533 mg, 2.2 mmol) in acetone (30 mL). For deprotection the crude intermediate was dissolved in DCM (18 mL) and cooled down to 0 °C. TFA (1.99 mL, 26.0 mmol) was added and the resulting mixture was stirred for 16 h, allowing the mixture to slowly reach room temperature. The solvent was removed under reduced pressure and the crude product was purified by HPLC to obtain **23** (202 mg, 0.68 mmol, 34 % over two steps) as a white solid. **LC-MS (ESI)**: $t_R$ = 7.79 min; *m/z* = 282.24 $[M – OH]^+$, calcd. for $C_{16}H_{18}BNO_4$: 299.13. **$^1$H NMR** (400 MHz, DMSO): $\delta$ = 8.41 – 8.32 (m, 1H), 8.14 (d, *J* = 4.5 Hz, 2H), 7.64 – 7.59 (m, 1H), 7.37 (td, *J* = 7.9, 7.2, 1.8 Hz, 1H), 7.30 – 7.25 (m, 2H), 7.22 – 7.18 (m, 3H), 6.98 (td, *J* = 7.3, 1.8 Hz, 1H), 6.88 (dd, *J* = 8.3, 1.9 Hz, 1H), 4.56 (s, 2H), 3.42 – 3.31 (m, 2H), 2.80 – 2.70 (m, 2H). **$^{13}$C NMR** (101 MHz, DMSO): $\delta$ = 168.31, 162.12, 139.17, 135.65, 131.62, 128.61, 128.32, 126.12, 121.12, 112.31, 67.15, 40.48, 34.99. **HRMS (ESI)**: *m/z* = 300.1402 calcd. for $[C_{16}H_{18}BNO_4 + H]^+$; found: 300.1396.

### Synthesis of intermediates for the derivative 24

#### Synthesis of 5-(tert-Butyl)-2-methoxybenzenesulfonamide (**S-24**)

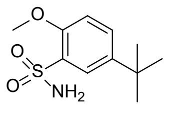

**S-24**

4-*tert*-butylanisole (4.11 g, 25.0 mmol) was dissolved in dry DCM (10 mL) and cooled down to 0 °C. Chlorsulfonic acid (5.66 mL, 85.0 mmol) was dissolved in dry DCM (6.0 mL) and added to te reaction mixture. The mixture was stirred for 1 h at 0 °C and for 1 h at room temperature. The reaction mixture was poured into ice-cooled water, the organic phase was quickly seperated washed with ice-cooled water (3x), dried over $MgSO_4$ and the solvent was removed under reduced pressure. The intermediate was directly dissolved in acetonitrile (10 mL) and cooled down to 0 °C. $NH_4OH$ solution (25%, 4.0 mL) was added and the resulting mixture was stirred for 16 h, allowing the mixture to slowly reach room temperature. The mixture was poured into ice-water and neutralized with concentrated HCl. The resulting solid was filtered and dried under reduced pressure to obtain **S-24** (3.96 g, 16.3 mmol, 65 %) as slightly yellow solid. **LC-MS (ESI)**: $t_R$ = 8.35 min; *m/z* = 243.72 $[M + H]^+$, calcd. for $C_{11}H_{17}NO_3S$: 243.09. **$^1$H NMR** (400 MHz, $CDCl_3$): $\delta$ = 7.90 (dd, *J* = 2.5, 1.1 Hz, 1H), 7.55 (dd, *J* = 8.7, 2.6 Hz, 1H), 6.99 (d, *J* = 8.7 Hz, 1H), 4.88 (bs, 2H), 3.99 (s, 3H), 1.30 (s, 9H). **$^{13}$C NMR** (101 MHz, $CDCl_3$): $\delta$ = 154.00, 144.08, 131.31, 129.41, 125.37, 112.05, 56.61, 34.54, 31.41.

#### Synthesis of 2-Hydroxybenzenesulfonamide (**S-25**)

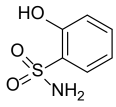

**S-25**

The protected sulfonamide **S-24** (3.89 g, 16.0 mmol) was dissolved in dry *m*-xylene (48 mL). $AlCl_3$ (8.85 g, 66.4 mmol) was added and the reaxtion mixture was stirred for 24 h at 60 °C. After cooling down to room temperature the black oil was poured into ice-water and hexane (20 mL) was used to rinse the reaction flask. The whole mixture was stirred at room temperature till a green suspension has been formed (about 30 min). The phases were seperated, the aqoueus phase was saturated with NaCl and extracted with diethyl ether. The combined organic diethyl ether phases were dired over $MgSO_4$ and the solvent was removed under reduced pressure to obtain **S-25** (1.54 g, 8.9 mmol, 56 %) as a white solid. **$^1$H NMR** (400 MHz, DMSO): $\delta$ = 10.62 (s, 1H), 7.64 (d, *J* = 7.9 Hz, 1H), 7.40 (t, *J* = 7.8 Hz, 1H), 6.98 (d, *J* = 8.2 Hz, 1H), 6.95 – 6.89 (m, 3H). **$^{13}$C NMR** (101 MHz, DMSO): $\delta$ = 154.36, 133.27, 129.25, 127.25, 118.26, 116.74.

### Synthesis of the derivative 24

#### Synthesis of N-phenethyl-2-(2-sulfamoylphenoxy)acetamide (**24**)

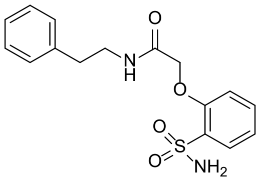

**24**

The sulfonamide derivative **24** was synthesized via GP **G** using **S-25** (384 mg, 2.22 mmol), potassium carbonate (608 mg, 4.44 mmol) and **S-16** (586 mg, 2.44 mmol) in acetone (35 mL). The crude product was purified by HPLC to obtain **24** (411 mg, 1.23 mmol, 55 %) as a white solid. **LC-MS (ESI)**: $t_R$ = 7.75 min; *m/z* = 334.95 $[M + H]^+$, calcd. for $C_{16}H_{18}N_2O_4S$: 334.10 . **$^1$H NMR** (400 MHz, DMSO): $\delta$ = 8.20 (t, *J* = 5.6 Hz, 1H), 7.77 (d, *J* = 7.7 Hz, 1H), 7.57 (t, *J* = 7.9 Hz, 1H), 7.35 (s, 2H), 7.32 – 7.24 (m, 2H), 7.21 – 7.12 (m, 5H), 4.70 (s, 2H), 3.47 – 3.28 (m, 2H), 2.74 (t, *J* = 7.4 Hz, 2H). **$^{13}$C NMR** (101 MHz, DMSO): $\delta$ = 167.52, 154.42, 139.08, 133.90, 131.77, 128.35, 127.54, 126.16, 121.04, 114.16, 67.55, 34.96. **HRMS (ESI)**: *m/z* = 335.1060 calcd. for $[C_{16}H_{18}N_2O_4S + H]^+$; found: 335.1056.

### Synthesis of intermediates for the derivatives 25 – 28

#### Synthesis of 2-Brom-N-((1R,2S)-2-phenylcyclopropyl)acetamide (**S-26**)

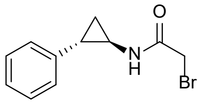

**S-26**

The bromoacetamide **S-26** was synthesized via GP **D** using *(1R,2S)-2-phenylcyclopropylamine (*100 mg, 751 µmol*),* triethylamine (135 µL, 976 µmol) and bromoacetyl bromide (79 µL, 901 µmol) in dry DCM (5 mL). The crude product was purified by HPLC to obtain **S-26** (129 mg, 507 µmol, 68 %) as a white solid. **LC-MS (ESI)**: $t_R$ = 8.01 min; *m/z* = 253.93 $[M + H]^+$, calcd. for $C_{11}H_{12}BrNO$: 253.01. **$^1$H NMR** (400 MHz, $CDCl_3$): $\delta$ = 7.27 – 7.08 (m, 2H), 7.12 – 6.96 (m, 3H), 6.82 (s, 1H), 3.76 (s, 2H), 2.84 (tt, *J* = 7.6, 3.5 Hz, 1H), 2.05 – 1.99 (m, 1H), 1.22 – 1.09 (m, 2H). **$^{13}$C NMR** (101 MHz, $CDCl_3$): $\delta$ = 166.89, 140.04, 128.50, 126.38, 32.46, 28.95, 24.62, 15.94.

#### Synthesis of 2-Brom-N-((1S,2R)-2-phenylcyclopropyl)acetamide (**S-27**)

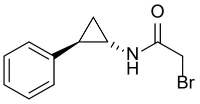

**S-27**

The bromoacetamide **S-27** was synthesized via GP **D** using *(1S,2R)-2-phenylcyclopropylamine (*100 mg, 751 µmol*),* triethylamine (135 µL, 976 µmol) and bromoacetyl bromide (79 µL, 901 µmol) in dry DCM (5 mL). The crude product was purified by HPLC to obtain **S-27** (147 mg, 578 µmol, 77 %) as a white solid. **LC-MS (ESI)**: $t_R$ = 8.03 min; *m/z* = 253.93 [M + H]+, calcd. for $C_{11}H_{12}BrNO$: 253.01. **$^1$H NMR** (400 MHz, $CDCl_3$): *δ* = 7.23 – 7.17 (m, 2H), 7.14 – 7.04 (m, 3H), 6.77 (s, 1H), 3.78 (s, 2H), 2.83 (tt, *J* = 7.5, 3.5 Hz, 1H), 2.06 – 1.99 (m, 1H), 1.22 – 1.11 (m, 2H). **$^{13}$C NMR** (101 MHz, $CDCl_3$): *δ* = 166.85, 140.04, 128.52, 126.57, 126.41, 32.47, 28.97, 24.66, 15.97.

#### Synthesis of (rac.)-2-Brom-N-(cis-2-phenylcyclopropyl)acetamide (**S-28**)

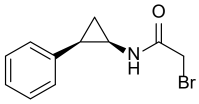

**S-28**

The bromoacetamide **S-28** was synthesized via GP **D** using *cis*-2-phenylcyclopropylamine hydrochloride (100 mg, 589 µmol), triethylamine (106 µL, 766 µmol) and bromoacetyl bromide (62 µL, 707 µmol) in dry DCM (5 mL). The crude product was purified by HPLC to obtain **S-28** (115 mg, 454 µmol, 77 %) as a white solid. **LC-MS (ESI)**: $t_R$ = 7.57 min; *m/z* = 253.87 [M + H]+, calcd. for $C_{11}H_{12}BrNO$: 253.01. **$^1$H NMR** (400 MHz, $CDCl_3$): *δ* = 7.29 – 7.21 (m, 2H), 7.20 – 7.10 (m, 3H), 6.12 (s, 1H), 3.64 – 3.51 (m, 2H), 3.10 – 2.99 (m, 1H), 2.36 – 2.25 (m, 1H), 1.38 – 1.26 (m, 1H), 1.04 – 0.98 (m, 1H). **$^{13}$C NMR** (101 MHz, $CDCl_3$): *δ* = 166.82, 135.66, 128.80, 128.55, 126.92, 42.41, 28.77, 21.71, 11.75.

#### Synthesis of2-Brom-N-((1R,2R)-2-phenylcyclopentyl)acetamide (**S-29**)

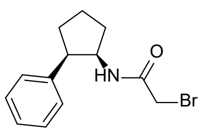

**S-29**

The bromoacetamide **S-29** was synthesized via GP **D** using (*1R,2R)-2-phenylcyclopentamine (*161 mg, 1.00 mol*),* triethylamine (180 µL, 1.30 mol) and bromoacetyl bromide (105 µL, 1.20 mmol) in dry DCM (5 mL). The crude product was purified by HPLC to obtain **S-29** (246 mg, 0.87 mmol, 87 %) as a slightly yellow oil. **LC-MS (ESI)**: $t_R$ = 8.86 min; *m/z* = 282.06 [M + H]+, calcd. for $C_{13}H_{16}BrNO$: 281.04. **$^1$H NMR** (400 MHz, $CDCl_3$): *δ* = 7.35 – 7.28 (m, 2H), 7.22 – 7.08 (m, 3H), 6.11 (s, 1H), 4.44 (p, *J* = 7.3 Hz, 1H), 3.63 (d, *J* = 13.7 Hz, 1H), 3.56 (d, *J* = 13.7 Hz, 1H), 3.37 (q, *J* = 7.5 Hz, 1H), 2.22 – 2.08 (m, 1H), 2.04 – 1.89 (m, 1H), 1.82 – 1.71 (m, 1H), 1,69 – 1.59 (m, 1H). **$^{13}$C NMR** (101 MHz, $CDCl_3$): *δ* = 164.78, 140.35, 128.55, 126.82, 54.41, 47.44, 31.84, 29.39, 22.46.

### Synthesis of the derivatives 25 – 28

#### Synthesis of (2-(2-Oxo-2-(((1R,2S)-2-phenylcyclopropyl)amino)ethoxy)phenyl)phosphonic acid (**25**)

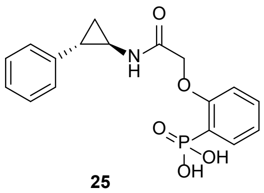

**25**

The phenylphosphonic acid derivative **25** was synthesized via GP **G** and GP **H**. GP **G** employed phosphonate **S-2** (156 mg, 0.78 mmol), potassium carbonate (214 mg, 1.55 mmol) and **S-26** (217 mg, 0.85 mmol) in acetone (15 mL). The crude intermediate was deprotected via GP **H** using TMSBr (273 µL, 2.07 mmol) in dry DCM (3.0 mL) and MeOH/$H_2O$ (3:1, 3.0 mL). The crude product was purified by HPLC to obtain **25** (64 mg, 0.19 mmol, 24 % over two steps) as a white solid. **LC-MS (ESI)**: $t_R$ = 5.99 min; *m/z* = 347.91 [M + H]+, cacld. for $C_{17}H_{18}NO_5P$: 347.09. **$^1$H NMR** (400 MHz, DMSO): *δ* = 9.41 (d, *J* = 5.3 Hz, 1H), 7.66 – 7.60 (m, 1H), 7.49 (t, *J* = 7.9 Hz, 1H), 7.25 (t, *J* = 7.5 Hz, 2H), 7.17 – 7.06 (m, 5H), 4.64 (s, 2H), 2.97 (s, 1H), 2.01 (s, 1H), 1.34 – 1.30 (m, 1H), 1.18 – 1.13 (m, 1H). **$^{13}$C NMR** (101 MHz, DMSO): *δ* = 168.90, 158.98, 141.41, 133.08, 132.68, 128.29, 125.80, 125.66, 123.28, 121.18, 113.26, 67.86, 32.46, 23.69, 14.95. **HRMS (ESI)**: *m/z* = 348.0996 calcd. for [$C_{17}H_{18}NO_5P$ + H]+; found: 348.0993.

#### Synthesis of (2-(2-Oxo-2-(((1S,2R)-2-phenylcyclopropyl)amino)ethoxy)phenyl)phosphonic acid (**26**)

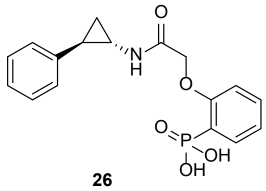

**26**

The phenylphosphonic acid derivative **26** was synthesized via GP **G** and GP **H**. GP **G** employed phosphonate **S-2** (174 mg, 0.64 mmol), potassium carbonate (239 mg, 1.73 mmol) and **S-27** (241 mg, 0.95 mmol) in acetone (16 mL). The crude intermediate was deprotected via GP **H** using TMSBr (0.57 mL, 4.32 mmol) in dry DCM (6.0 mL) and MeOH/$H_2O$ (3:1, 6.0 mL). The crude product was purified by HPLC to obtain **26** (71 mg, 0.21 mmol, 33 % over two steps) as a white solid. **LC-MS (ESI)**: $t_R$ = 6.07 min; *m/z* = 347.90 [M + H]$^+$, calcd. for $C_{17}H_{18}NO_5P$: 347.09. **$^1$H NMR** (400 MHz, DMSO): *δ* = 9.40 (d, *J* = 5.3 Hz, 1H), 7.66 – 7.60 (m, 1H), 7.49 (t, *J* = 8.1 Hz, 1H), 7.29 – 7.23 (m, 2H), 7.17 – 7.04 (m, 5H), 4.63 (s, 2H), 2.98 – 2.94 (m, 1H), 2.03 – 1.98 (m, 1H), 1.34 – 1.28 (m, 1H), 1.26 – 1.10 (m, 2H). **$^{13}$C NMR** (101 MHz, DMSO): *δ* = 168.69, 158.76, 141.20, 132.59, 128.07, 126.24, 125.58, 123.40, 121.16, 113.04, 67.64, 32.25, 23.47, 14.73. **HRMS (ESI)**: *m/z* = 348.0996 calcd. for [$C_{17}H_{18}NO_5P$ + H]$^+$; found: 348.0993.

#### Synthesis of (2-(2-Oxo-2-(((1R,2R)-2-phenylcyclopropyl)amino)ethoxy)phenyl)phosphonic acid (**27**)

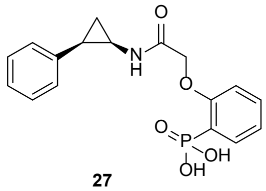

**27**

The phenylphosphonic acid derivative **27** was synthesized via GP **G** and GP **H**. GP **G** employed phosphonate **S-2** (83 mg, 413 µmol), potassium carbonate (115 mg, 826 µmol) and **S-28** (115 mg, 454 µmol) in acetone (8.0 mL). The crude intermediate was deprotected via GP **H** using TMSBr (273 µL, 2.07 mmol) in dry DCM (3.0 mL) and MeOH/$H_2O$ (3:1, 3.0 mL). The crude product was purified by HPLC to obtain **27** (47 mg, 134 µmol, 33 % over two steps) as a white solid. **LC-MS (ESI)**: $t_R$ = 5.69 min; *m/z* = 348.01 [M + H]$^+$, calcd. for $C_{17}H_{18}NO_5P$: 347.09. **$^1$H NMR** (400 MHz, DMSO): *δ* = 9.22 (d, *J* = 4.4 Hz, 2H), 7.68 – 7.62 (m, 1H), 7.45 – 7.41 (m, 2H), 7.12 – 6.95 (m, 7H), 4.45 (d, *J* = 15.3 Hz, 1H), 4.31 (d, *J* = 15.3 Hz, 1H), 3.18 (s, 1H), 2.15 (q, *J* = 7.8 Hz, 1H), 1.32 – 1.23 (m, 2H). **$^{13}$C NMR** (101 MHz, DMSO): *δ* = 169.21, 158.99, 137.65, 133.18, 132.67, 129.68, 127.81, 127.42, 125.16, 123.06, 121.19, 113.04, 67.42, 48.71, 30.06, 22.71, 10.53. **HRMS (ESI)**: *m/z* = 348.0996 calcd. for [$C_{17}H_{18}NO_5P$ + H]$^+$; found: 348.0994.

#### Synthesis of (2-(2-Oxo-2-(((1R,2R)-2-phenylcyclopentyl)amino)ethoxy)phenyl)phosphonic acid (**28**)

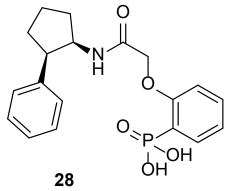

**28**

The phenylphosphonic acid derivative **28** was synthesized via GP **G** and GP **H**. GP **G** employed phosphonate **S-2** (178 mg, 790 µmol), potassium carbonate (249 mg, 1.59 mmol) and **S-29** (246 mg, 870 µmol) in acetone (10 mL). The crude intermediate was deprotected via GP **H** using TMSBr (1.04 mL, 7.90 mmol) in dry DCM (6.0 mL) and MeOH/$H_2O$ (3:1, 6.0 mL). The crude product was purified by HPLC to obtain **28** (159 mg, 424 µmol, 54 % over two steps) as a white solid. **LC-MS (ESI)**: $t_R$ = 6.48 min; *m/z* = 376.10 [M + H]$^+$, calcd. for $C_{19}H_{22}NO_5P$: 375.12. **$^1$H NMR** (400 MHz, DMSO): *δ* = 8.43 (d, *J* = 9.6 Hz, 1H), 7.72 – 7.66 (m, 1H), 7.45 (t, *J* = 7.9 Hz, 1H), 7.26 – 7.19 (m, 2H), 7.05 (d, *J* = 5.2 Hz, 4H), 6.94 (t, *J* = 7.2 Hz, 1H), 4.58 – 4.50 (m, 1H), 4.32 (d, *J* = 14.8 Hz, 1H), 4.21 (d, *J* = 14.9 Hz, 1H), 3.13 (dt, *J* = 12.8, 6.8 Hz, 1H), 2.22 – 1.95 (m, 3H), 1.91 – 1.83 (m, 1H), 1.75 – 1.68 (m, 1H), 1.64 – 1.56 (m, 1H). **$^{13}$C NMR** (101 MHz, DMSO): *δ* = 166.39, 158.60, 140.59, 133.08, 132.65, 128.74, 127.34, 125.63, 122.74, 120.95, 112.58, 66.89, 52.53, 49.58, 32.06, 29.51, 22.36. **HRMS (ESI)**: *m/z* = 376.1309 calcd. for [$C_{19}H_{22}NO_5P$ + H]$^+$; found: 376.1306.

#### **Synthesis of the derivatives 29 – 31**

#### Synthesis of (2-(2-oxo-2-(((1-phenylcyclobutyl)methyl)amino)ethoxy)phenyl)phosphonic acid (**29**)

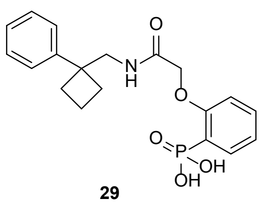

29

The phenylphosphonic acid derivative **29** was synthesized via GP **F** and GP **H**. GP **F** employed phosphonate **S-3** (260 mg, 1.00 mmol), *N*-Methylmorpholine (111 µL, 1.00 mmol) and isobutyl chloroformate (130 µL, 1.00 mmol) in dry THF (10 mL) as well as *N*-Methylmorpholine (111 µL, 1.00 mmol) and (1-phenylcylobutyl)methylamine (161 mg, 1.00 mmol). The crude intermediate was deprotected via GP **H** using TMSBr (1.32 mL, 10.00 mmol) in dry DCM (8 mL) and MeOH/$H_2O$ (3:1, 8 mL). The crude product was purified by HPLC to obtain **29** (227 mg, 0.60 mmol, 60 % over two steps) as a white solid. **LC-MS (ESI)**: $t_R$ = 6.67 min; *m/z* = 376.14 [M + H]$^+$, calcd. for $C_{19}H_{22}NO_5P$: 375.12 . **$^1$H NMR** (400 MHz, DMSO): $\delta$ = 8.83 (t, *J* = 6.2 Hz, 1H), 7.69 – 7.62 (m, 1H), 7.55 – 7.46 (m, 1H), 7.21 (t, *J* = 7.4 Hz, 2H), 7.16 – 7.03 (m, 5H), 4.55 (s, 2H), 3.39 (d, *J* = 6.2 Hz, 2H), 2.41 – 2.31 (m, 2H), 2.22 – 2.09 (m, 2H), 1.89 – 1.76 (m, 1H), 1.70 – 1.58 (m, 1H). **$^{13}$C NMR** (101 MHz, DMSO): $\delta$ = 167.69, 158.39, 148.05, 132.91, 132.49, 127.59, 125.59, 125.14, 120.82, 112.92, 67.08, 66.79, 47.67, 46.45, 31.10, 29.86, 14.98. **HRMS (ESI)**: *m/z* = 376.1309 calcd. for [$C_{19}H_{22}NO_5P$ + H]$^+$; found: 376.1307.

#### Synthesis of 2-(2-((2,2-Difluoro-2-phenylethyl)amino)-2-oxoethoxy)phenyl)phosphonic acid (**30**)

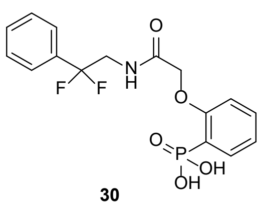

30

The phenylphosphonic acid derivative **30** was synthesized via GP **F** and GP **H**. GP **F** employed phosphonate **S-3** (260 mg, 1.00 mmol), *N*-Methylmorpholine (111 µL, 1.00 mmol) and isobutyl chloroformate (130 µL, 1.00 mmol) in dry THF (10 mL) as well as *N*-Methylmorpholine (111 µL, 1.00 mmol) and 2,2-difluoro-2-phenylethanamine (157 mg, 1.00 mmol). The crude intermediate was deprotected via GP **H** using TMSBr (1.32 mL, 10.00 mmol) in dry DCM (8 mL) and MeOH/$H_2O$ (3:1, 8 mL). The crude product was purified by HPLC to obtain **30** (158 mg, 0.43 mmol, 43 % over two steps) as a white solid. **LC-MS (ESI)**: $t_R$ = 6.11 min; *m/z* = 371.88 [M + H]$^+$, calcd. for $C_{16}H_{16}F_2NO_5P$: 371.07. **$^1$H NMR** (400 MHz, DMSO): $\delta$ = 9.39 (t, *J* = 6.3 Hz, 1H), 7.68 – 7.59 (m, 1H), 7.54 – 7.38 (m, 6H), 7.12 – 7.05 (m, 2H), 4.64 (s, 2H), 3.87 (td, *J* = 15.0, 6.3 Hz, 2H). **$^{13}$C NMR** (101 MHz, DMSO): $\delta$ = 168.65, 158.78, 134.68, 133.16, 132.59, 130.30, 128.56, 125.08, 123.14, 121.19, 120.57, 113.18, 67.52, 44.09. **HRMS (ESI)**: *m/z* = 372.0807 calcd. for [$C_{16}H_{16}F_2NO_5P$ + H]$^+$; found: 372.0806.

#### Synthesis of 2-(2-((2-Methyl-2-phenylpropyl)amino)-2-oxoethoxy)phenyl)phosphonic acid (**31**)

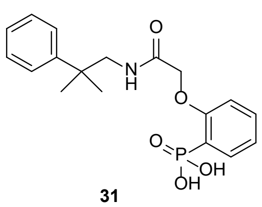

31

The phenylphosphonic acid derivative **31** was synthesized via GP **F** and GP **H**. GP **F** employed phosphonate **S-3** (260 mg, 1.00 mmol), *N*-Methylmorpholine (111 µL, 1.00 mmol) and isobutyl chloroformate (130 µL, 1.00 mmol) in dry THF (10 mL) as well as *N*-Methylmorpholine (111 µL, 1.00 mmol) and 2-Methyl-2-phenylpropanamine (149 mg, 1.00 mmol). The crude intermediate was deprotected via GP **H** using TMSBr (1.32 mL, 10.00 mmol) in dry DCM (8 mL) and MeOH/$H_2O$ (3:1, 8 mL). The crude product was purified by HPLC to obtain **31** (182 mg, 0.50 mmol, 50 % over two steps) as a white solid. **LC-MS (ESI)**: $t_R$ = 6.43 min; *m/z* = 364.09 [M + H]$^+$, calcd. for $C_{18}H_{22}NO_5P$: 363.12. **$^1$H NMR** (400 MHz, DMSO): $\delta$ = 8.64 (t, *J* = 6.3 Hz, 1H), 7.67 – 7.61 (m, 1H), 7.49 (t, *J* = 8.0 Hz, 1H), 7.35 (d, *J* = 7.5 Hz, 2H), 7.27 (t, *J* = 7.7 Hz, 2H), 7.21 – 7.12 (m, 1H), 7.14 – 7.01 (m, 2H), 4.60 (s, 2H), 3.30 (d, *J* = 6.3 Hz, 2H), 1.22 (s, 6H). **$^{13}$C NMR** (101 MHz, DMSO): $\delta$ = 167.98, 158.66, 147.61, 133.08, 132.57, 128.04, 125.70, 123.08, 121.30, 120.98, 112.87, 67.27, 49.45, 26.14. **HRMS (ESI)**: *m/z* = 364.1309 calcd. for [$C_{18}H_{22}NO_5P$ + H]$^+$; found: 364.1306.

#### **Synthesis of intermediates for the derivatives 40 – 45**

##### Synthesis of ethyl 2,2- difluoro-2-(4-chlorophenyl)acetate (**S-30**)

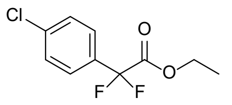

**S-30**

The phenylacetate derivative **S-30** was synthesized via GP **O** using 4-chloroiodobenzene (2.39 g, 10.00 mmol), ethyl bromodifluoroacetate (1.28 mL, 10.00 mmol) and activated copper (1.65 g, 26.00 mmol) in dry DMF (15 mL). The crude product was purified by column chromatography (cyclohexane/ethyl acetate 19:1) to obtain **S-30** (1.27 g, 5.41 mmol, 54 %) as a colorless oil. **TLC** (cyclohexane:ethyl acetate, 19:1 v/v): $R_f$ = 0.45. **$^1$H NMR** (400 MHz, $CDCl_3$): $\delta$ = 7.53 (d, $J$ = 8.8 Hz, 2H), 7.43 (d, $J$ = 8.8 Hz, 2H), 4.28 (q, $J$ = 7.1 Hz, 2H), 1.28 (t, $J$ = 7.2 Hz, 8H). **$^{13}$C NMR** (101 MHz, $CDCl_3$): $\delta$ = 170.77, 163.89, 137.42, 131.40, 129.06, 127.14, 113.05, 63.41, 13.91.

#### Synthesis of ethyl 2-(4-bromophenyl)-2,2-difluoracetate (**S-31**)

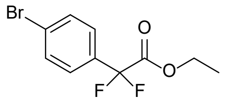

**S-31**

The phenylacetate derivative **S-31** was synthesized via GP **O** using 4-bromoiodobenzene (2.83 g, 10.00 mmol), ethyl bromodifluoroacetate (1.95 mL, 15.00 mmol) and activated copper (2.55 g, 39.00 mmol) in dry DMSO (26 mL). The crude product was purified by column chromatography (cyclohexane/ethyl acetate 19:1) to obtain **S-31** (2.38 g, 8.51 mmol, 85 %) as a colorless oil. **TLC** (cyclohexane:ethyl acetate, 19:1 v/v): $R_f$ = 0.50. **$^1$H NMR** (400 MHz, $CDCl_3$): $\delta$ = 7.58 (d, $J$ = 8.7 Hz, 1H), 7.49 (d, $J$ = 8.5 Hz, 2H), 4.30 (q, $J$ = 7.1 Hz, 2H), 1.29 (t, $J$ = 7.2 Hz, 3H). **$^{13}$C NMR** (101 MHz, $CDCl_3$): $\delta$ = 163.84, 132.05, 127.25, 125.79, 113.12, 63.42, 13.94.

#### Synthesis of ethyl 2,2-difluoro-2-(2-fluorophenyl)acetate (**S-32**)

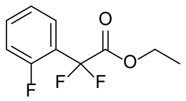

**S-32**

The phenylacetate derivative **S-32** was synthesized via GP **O** using 2-fluoroiodobenzene (1.16 mL, 10.00 mmol), ethyl bromodifluoroacetate (1.95 mL, 15.00 mmol) and activated copper (2.55 g, 39.00 mmol) in dry DMSO (26 mL). The crude product was purified by column chromatography (cyclohexane/ethyl acetate 19:1) to obtain **S-32** (1.80 g, 8.26 mmol, 83 %) as a colorless oil. **TLC** (cyclohexane:ethyl acetate, 19:1 v/v): $R_f$ = 0.37. **LC-MS (ESI)**: $t_R$ = 9.61 min; *m/z* = 218.58 [M + H]$^+$, calcd. for $C_{10}H_{19}F_3O_2$: 218.06. **$^1$H NMR** (400 MHz, $CDCl_3$): $\delta$ = 7.66 (t, $J$ = 7.6 Hz, 1H), 7.55 – 7.45 (m, 1H), 7.26 (d, $J$ = 15.4 Hz, 1H), 7.19 – 7.09 (m, 1H), 4.35 (q, $J$ = 7.2 Hz, 2H), 1.32 (t, $J$ = 7.2 Hz, 3H). **$^{13}$C NMR** (101 MHz, $CDCl_3$): $\delta$ = 163.32, 158.60, 133.14, 127.17, 124.37, 120.99, 116.25, 111.76, 63.42, 13.83.

#### Synthesis of ethyl 2,2-difluoro-2-(2-fluorophenyl)acetate (**S-33**)

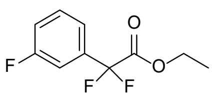

**S-33**

The phenylacetate derivative **S-33** was synthesized via GP **O** using 3-fluoroiodobenzene (1.17 mL, 10.00 mmol), ethyl bromodifluoroacetate (1.95 mL, 15.00 mmol) and activated copper (2.55 g, 39.00 mmol) in dry DMSO (26 mL). The crude product was purified by column chromatography (cyclohexane/ethyl acetate 19:1) to obtain **S-33** (10.99 g, 4.54 mmol, 45 %) as a colorless oil. **TLC** (cyclohexane:ethyl acetate, 19:1 v/v): $R_f$ = 0.43. **$^1$H NMR** (400 MHz, $CDCl_3$): $\delta$ = 7.46 – 7.38 (m, 2H), 7.31 (d, $J$ = 9.1 Hz, 1H), 7.19 (t, $J$ = 7.3 Hz, 1H), 4.30 (q, $J$ = 7.1 Hz, 2H), 1.31 (t, $J$ = 7.2 Hz, 3H). **$^{13}$C NMR** (101 MHz, $CDCl_3$): $\delta$ = 163.92, 161.46, 135.14, 130.62, 121.31, 118.38, 113.29, 112.74, 63.48, 13.96.

#### Synthesis of ethyl 2,2-difluoro-2-(4-fluorophenyl)acetate (**S-34**)

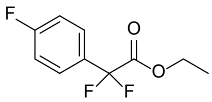

**S-34**

The phenylacetate derivative **S-34** was synthesized via GP **O** using 4-fluoroiodobenzene (1.15 mL, 10.00 mmol), ethyl bromodifluoroacetate (1.28 mL, 10.00 mmol) and activated copper (1.65 g, 26.00 mmol) in dry DMF (15 mL). The crude product was purified by column chromatography (cyclohexane/ethyl acetate 19:1) to obtain **S-34** (0.81 g, 3.71 mmol, 37 %) as a colorless oil. **TLC** (cyclohexane:ethyl acetate, 19:1 v/v): $R_f$ = 0.49. **LC-MS (ESI)**: $t_R$ = 9.98 min; *m/z* = 219.26 [M + H]$^+$, calcd. for $C_{10}H_9F_3O_2$: 218.06.
**$^1$H NMR** (400 MHz, $CDCl_3$): $\delta$ = 7.65 – 7.55 (m, 2H), 7.12 (t, $J$ = 8.6 Hz, 2H), 4.28 (q, $J$ = 7.1 Hz, 2H), 1.28 (t, $J$ = 7.2 Hz, 1H). **$^{13}$C NMR** (101 MHz, $CDCl_3$): $\delta$ = 165.75, 164.07, 163.11, 128.97, 127.85, 116.02, 113.15, 63.33, 13.87.

#### Synthesis of ethyl 2,2-difluoro-2-(3-(trifluoromethyl)phenyl)acetate (**S-35**)

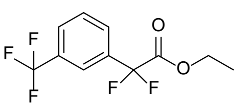

**S-35**

The phenylacetate derivative **S-35** was synthesized via GP **O** using 3-iodobenzotrifluorid (7.21 mL, 50.0 mmol), ethyl bromodifluoroacetate (9.62 mL, 75.0 mmol) and activated copper (12.75 g, 195.0 mmol) in dry DMSO (130 mL). The crude product was purified by column chromatography (cyclohexane/ethyl acetate 19:1) to obtain **S-35** (8.52 g, 31.8 mmol, 64 %) as a colorless oil. **TLC** (cyclohexane:ethyl acetate, 19:1 v/v): $\mathbf{R_f}$ = 0.50. $^{1}$**H NMR** (400 MHz, $CDCl_3$): *δ* = 7.88 (s, 1H), 7.68 (dd, *J* = 23.3, 8.0 Hz, 2H), 7.59 (t, *J* = 7.9 Hz, 1H), 4.30 (q, *J* = 7.2 Hz, 2H), 1.29 (t, *J* = 7.2 Hz, 1H). $^{13}$**C NMR** (101 MHz, $CDCl_3$): *δ* = 163.64, 134.11, 131.64, 129.54, 129.14, 127.96, 125.05, 122.77, 112.82, 63.59, 13.79.

#### Synthesis of 2,2-difluoro-2-(3-(trifluoromethyl)phenyl)ethanol (**S-36**)

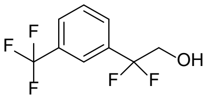

**S-36**

The alcohol **S-36** was synthesized via GP **P** using the acetate **S-35** (8.28 g, 30.88 mmol), and $NaBH_4$ (1.17 g, 30.88 mmol) in MeOH (100 mL). The crude product was purified by column chromatography (cyclohexane/ethyl acetate 2:1) to obtain **S-36** (5.09 g, 22.50 mmol, 73 %) as a colorless oil. **TLC** (cyclohexane:ethyl acetate, 2:1 v/v): $\mathbf{R_f}$ = 0.38. $^{1}$**H NMR** (400 MHz, MeOD): *δ* = 7.89 – 7.76 (m, 3H), 7.68 (q, *J* = 7.9, 7.3 Hz, 1H), 3.97 (t, *J* = 13.1 Hz, 4H). $^{13}$**C NMR** (101 MHz, MeOD): *δ* = 138.05, 132.03, 130.60, 127.84, 126.63, 123.70, 121.59, 119.17, 65.42.

#### Synthesis of 2-bromo-N-(2,2-difluoro-2-(4-chlorophenyl)ethyl)acetamide (**S-37**)

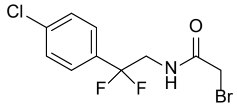

**S-37**

The bromoacetamide **S-37** was synthesized via GP **P**, GP **Q** and GP **D**. GP **P** employed the acetate **S-30** (877 mg, 3.67 mmol), and $NaBH_4$ (142 mg, 3.67 mmol) in MeOH (13 mL). The crude alcohol was converted to the amine via GP **Q** using pyridine (0.47 mL, 5.87 mmol) and $Tf_2O$ (0.68 mL, 4.04 mmol) in dry acetonirile (6.5 mL) as well as $NH_4OH$ (28%, 6.5 mL). The crude intermediate was coupled to the amide using triethylamine (0.56 mL, 4.04 mmol) and bromoacetyl bromide (0.32 mL, 4.04 mmol) in dry DCM (18 mL). The crude product was purified by column chromatography (cyclohexane/diethyl ether 1:4) to obtain **S-37** (216 mg, 0.69 mmol, 19 % over three steps) as an orange solid. **TLC** (cyclohexane:diethyl ether, 1:4 v/v): $\mathbf{R_f}$ = 0.43. **LC-MS (ESI)**: $t_R$ = 8.37 min; *m/z* = 311.45 $[M + H]^+$, calcd. for $C_{10}H_9BrClF_2NO$: 310.95. $^{1}$**H NMR** (400 MHz, $CDCl_3$): *δ* = 7.50 – 7.38 (m, 4H), 6.78 (s, 1H), 3.93 (td, *J* = 14.3, 6.3 Hz, 2H), 3.87 (s, 2H). $^{13}$**C NMR** (101 MHz, $CDCl_3$): *δ* = 165.76, 136.96, 132.83, 129.11, 126.82, 119.90, 45.91, 28.83.

#### Synthesis of 2-bromo-N-(2-(4-bromophenyl)-2,2-difluoroethyl)acetamide (**S-38**)

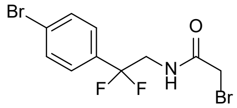

**S-38**

The bromoacetamide **S-38** was synthesized via GP **P**, GP **Q** and GP **D**. GP **P** employed the acetate **S-31** (2.12 g, 7.60 mmol), and $NaBH_4$ (0.29 g, 7.60 mmol) in MeOH (25 mL). The crude alcohol was converted to the amine via GP **Q** using pyridine (0.96 mL, 12.16 mmol) and $Tf_2O$ (1.44 mL, 8.36 mmol) in dry acetonirile (13 mL) as well as $NH_4OH$ (28%, 13 mL). The crude intermediate was coupled to the amide using triethylamine (1.16 mL, 8.36 mmol) and bromoacetyl bromide (0.73 mL, 8.36 mmol) in dry DCM (18 mL). The crude product was purified by column chromatography (cyclohexane/ethyl acetate 2:1) to obtain **S-38** (1.11 g, 3.10 mmol, 41 % over three steps) as an orange solid. **TLC** (cyclohexane:ethyl acetate, 2:1 v/v): $\mathbf{R_f}$ = 0.43. **LC-MS (ESI)**: $t_R$ = 9.04min; *m/z* = 355.58 $[M + H]^+$, calcd. for $C_{10}H_9Br_2F_2NO$: 354.90. $^{1}$**H NMR** (400 MHz, $CDCl_3$): *δ* = 7.56 (d, *J* = 8.2 Hz, 2H), 7.38 (d, *J* = 8.5 Hz, 1H), 6.88 (s, 1H), 3.92 (td, *J* = 14.3, 6.4 Hz, 2H), 3.85 (s, 2H). $^{13}$**C NMR** (101 MHz, $CDCl_3$): *δ* = 165.95, 133.33, 132.02, 127.12, 125.20, 119.92, 45.81, 28.72.

#### Synthesis of 2-bromo-N-(2,2-difluoro-2-(2-fluorophenyl)ethyl)acetamide (**S-39**)

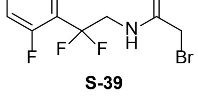

**S-39**

The bromoacetamide **S-39** was synthesized via GP **P**, GP **Q** and GP **D**. GP **P** employed the acetate **S-32** (1.45 g, 6.65 mmol), and $NaBH_4$ (0.25 g, 6.65 mmol) in MeOH (22 mL). The crude alcohol was converted to the amine via GP **Q** using pyridine (0.86 mL, 10.64 mmol) and $Tf_2O$ (1.23 mL, 7.32 mmol) in dry acetonirile (12 mL) as well as $NH_4OH$ (28%, 12 mL). The crude intermediate was coupled to the amide using triethylamine (1.01 mL, 7.32 mmol) and bromoacetyl bromide (0.64 mL, 7.32 mmol) in dry DCM (33 mL). The crude product was purified by column chromatography (cyclohexane/ethyl acetate 2:1) to obtain **S-39** (679 mg, 2.29 mmol, 34 % over three steps) as a white solid. **TLC** (cyclohexane:ethyl acetate, 2:1 v/v): $R_f$ = 0.29. **LC-MS (ESI)**: $t_R$ = 8.18 min; *m/z* = 295.62 [M + H]$^+$, calcd. for $C_{10}H_9BrF_3NO$: 294.98. $^1$**H NMR** (400 MHz, $CDCl_3$): $\delta$ = 7.53 (t, *J* = 7.6 Hz, 1H), 7.49 – 7.43 (m, 1H), 7.22 (t, *J* = 7.6 Hz, 1H), 7.18 – 7.12 (m, 1H), 6.82 (s, 1H), 6.24 (s, 0H), 4.07 (td, *J* = 14.6, 6.3 Hz, 2H), 3.85 (s, 2H). $^{13}$**C NMR** (101 MHz, $CDCl_3$): $\delta$ = 165.91, 158.60, 132.86, 127.50, 124.40, 121.69, 118.79, 116.85, 45.11, 28.77.

##### Synthesis of 2-bromo-N-(2,2-difluoro-2-(3-fluorophenyl)ethyl)acetamide (**S-40**)

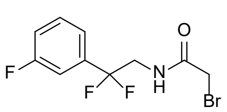

**S-40**

The bromoacetamide **S-40** was synthesized via GP **P**, GP **Q** and GP **D**. GP **P** employed the acetate **S-33** (0.73 g, 3.34 mmol), and $NaBH_4$ (0.13 g, 3.34 mmol) in MeOH (12 mL). The crude alcohol was converted to the amine via GP **Q** using pyridine (0.43 mL, 5.34 mmol) and $Tf_2O$ (0.62 mL, 3.67 mmol) in dry acetonirile (6 mL) as well as $NH_4OH$ (28%, 6 mL). The crude intermediate was coupled to the amide using triethylamine (0.51 mL, 3.67 mmol) and bromoacetyl bromide (0.32 mL, 3.67 mmol) in dry DCM (17 mL). The crude product was purified by column chromatography (cyclohexane/ethyl acetate 2:1) to obtain **S-40** (679 mg, 2.29 mmol, 34 % over three steps) as a white solid. **TLC** (cyclohexane:ethyl acetate, 2:1 v/v): $R_f$ = 0.40. **LC-MS (ESI)**: $t_R$ = 8.27 min; *m/z* = 295.61 [M + H]$^+$, calcd. for $C_{10}H_9BrF_3NO$: 294.98. $^1$**H NMR** (400 MHz, $CDCl_3$): $\delta$ = 7.46 – 7.40 (m, 1H), 7.29 (d, *J* = 7.8 Hz, 1H), 7.24 – 7.14 (m, 2H), 6.80 (s, 1H), 3.93 (td, *J* = 14.4, 6.3 Hz, 2H), 3.87 (s, 2H). $^{13}$**C NMR** (101 MHz, $CDCl_3$): $\delta$ = 165.83, 161.49, 136.63, 130.66, 121.15, 119.50, 117.70, 113.02, 45.92, 28.77.

##### Synthesis of 2-bromo-N-(2,2-difluoro-2-(4-fluorophenyl)ethyl)acetamide (**S-41**)

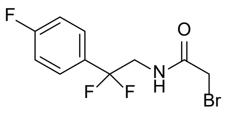

**S-41**

The bromoacetamide **S-41** was synthesized via GP **P**, GP **Q** and GP **D**. GP **P** employed the acetate **S-34** (561 mg, 2.57 mmol), and $NaBH_4$ (96 mg, 2,57 mmol) in MeOH (9 mL). The crude alcohol was converted to the amine via GP **Q** using pyridine (0.33 mL, 4.11 mmol) and $Tf_2O$ (0.48 mL, 2.83 mmol) in dry acetonirile (4.6 mL) as well as $NH_4OH$ (28%, 4.6 mL). The crude intermediate was coupled to the amide using triethylamine (0.39 mL, 2.83 mmol) and bromoacetyl bromide (0.22 mL, 2.83 mmol) in dry DCM (13 mL). The crude product was purified by column chromatography (cyclohexane/diethyl ether 1:4) to obtain **S-41** (200 mg, 0.67 mmol, 26 % over three steps) as a brown solid. **TLC** (cyclohexane:diethyl ether, 1:4 v/v): $R_f$ = 0.38. **LC-MS (ESI)**: $t_R$ = 8.23 min; *m/z* = 295.54 [M + H]$^+$, calcd. for $C_{10}H_9BrF_3NO$: 294.98. $^1$**H NMR** (400 MHz, $CDCl_3$): $\delta$ = 7.58 – 7.42 (m, 2H), 7.13 (t, *J* = 8.5 Hz, 2H), 6.80 (s, 1H), 3.92 (td, *J* = 14.3, 6.3 Hz, 2H), 3.87 (s, 2H). $^{13}$**C NMR** (101 MHz, $CDCl_3$): $\delta$ = 165.77, 130.35, 127.55, 119.97, 116.06, 115.84, 45.71, 28.84.

##### Synthesis of 2-bromo-N-(2,2-difluoro-2-(3-(trifluoromethyl)phenyl)ethyl)acetamide (**S-42**)

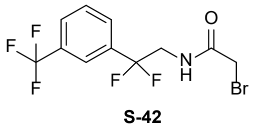

**S-42**

The bromoacetamide **S-42** was synthesized via GP **Q** and GP **D**. GP **Q** employed the alcohol **S-36** (4.94 g, 21.87 mmol), pyridine (2.82 mL, 34.99 mmol) and $Tf_2O$ (4.04 mL, 24.06 mmol) in dry acetonirile (40 mL) as well as $NH_4OH$ (28%, 40 mL). The crude intermediate was coupled to the amide using triethylamine (3.34 mL, 24.06 mmol) and bromoacetyl bromide (2.09 mL, 24.06 mmol) in dry DCM (90 mL). The crude product was purified by column chromatography (cyclohexane/ethyl acetate 2:1) to obtain **S-42** (2.89 g, 8.36 mmol, 38 % over two steps) as a white solid. **TLC** (cyclohexane:ethyl acetate, 2:1 v/v): $R_f$ = 0.20. **LC-MS (ESI)**: $t_R$ = 9.14 min; *m/z* = 345.61 [M + H]$^+$, calcd. for $C_{11}H_9BrF_5NO$: 344.98. $^1$**H NMR** (400 MHz, $CDCl_3$): $\delta$ = 7.80 – 7.68 (m, 3H), 7.60 (t, *J* = 7.8 Hz, 1H), 6.86 (s, 1H), 3.95 (td, *J* = 14.2, 6.3 Hz, 2H), 3.86 (s, 1H). $^{13}$**C NMR** (101 MHz, $CDCl_3$): $\delta$ = 166.04, 135.42, 131.25, 129.56, 128.86, 127.56, 125.05, 122.49, 119.62, 45.92, 28.63.

#### Synthesis of the derivatives 40 – 45

##### Synthesis of ((2-(2-((2-(4-chlorophenyl)-2,2-difluoroethyl)amino)-2-oxoethoxy)phenyl)phosphonic acid (**40**)

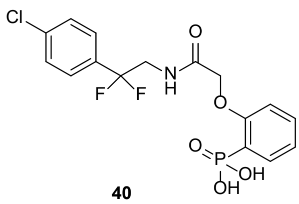

40

The phenylphosphonic acid derivative **40** was synthesized via GP **G** and GP **H**. GP **G** employed phosphonate **S-2** (123 mg, 0.61 mmol), potassium carbonate (169 mg, 1.22 mmol) and **S-37** (191 mg, 0.61 mmol) in DMF (6 mL). The crude intermediate was deprotected via GP **H** using TMSBr (0.81 mL, 6.10 mmol) in dry DCM (5 mL) and MeOH/$H_2O$ (3:1, 4 mL). The crude product was purified by HPLC to obtain **40** (138 mg, 0.34 mmol, 56 % over two steps) as a white solid. **LC-MS (ESI)**: $t_R$ = 6.67 min; *m/z* = 405.87 $[M + H]^+$, calcd. for $C_{16}H_{15}ClF_2NO_5P$: 405.03. **$^1$H NMR** (400 MHz, DMSO): *δ* = 9.46 (s, 1H), 7.71 – 7.58 (m, 1H), 7.53 – 7.40 (m, 5H), 7.16 – 7.06 (m, 2H), 4.65 (s, 2H), 3.88 (t, *J* = 12.6 Hz, 2H). **$^{13}$C NMR** (101 MHz, DMSO): *δ* = 168.58, 158.85, 135.01, 133.00, 128.51, 127.23, 121.21, 113.23, 99.50, 67.43, 43.84. **HRMS (ESI)**: *m/z* = 406.0417 calcd. for $[C_{16}H_{15}ClF_2NO_5P + H]^+$; found: 406.0417.

#### Synthesis of ((2-(2-((2-(4-bromophenyl)-2,2-difluoroethyl)amino)-2-oxoethoxy)phenyl)phosphonic acid (**41**)

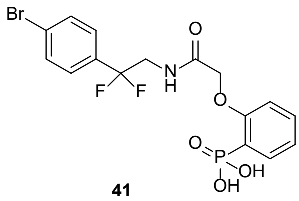

41

The phenylphosphonic acid derivative **41** was synthesized via GP **G** and GP **H**. GP **G** employed phosphonate **S-2** (202 mg, 1.00 mmol), potassium carbonate (276 mg, 2.00 mmol) and **S-38** (357 mg, 1.00 mmol) in DMF (6 mL). The crude intermediate was deprotected via GP **H** using TMSBr (1.32 mL, 10.00 mmol) in dry DCM (8 mL) and MeOH/$H_2O$ (3:1, 6 mL). The crude product was purified by HPLC to obtain **41** (181 mg, 0.40 mmol, 40 % over two steps) as a white solid. **LC-MS (ESI)**: $t_R$ = 6.82 min; *m/z* = 449.85 $[M + H]^+$, calcd. for $C_{16}H_{15}BrF_2NO_5P$: 448.98. **$^1$H NMR** (400 MHz, DMSO): *δ* = 9.44 (t, *J* = 6.2 Hz, 1H), 7.66 – 7.58 (m, 3H), 7.50 (t, *J* = 7.2 Hz, 1H), 7.38 (d, *J* = 8.3 Hz, 2H), 7.12 – 7.06 m, 2H), 4.64 (s, 2H), 3.88 (td, *J* = 14.4, 6.3 Hz, 2H). **$^{13}$C NMR** (101 MHz, $DMSO_3$): *δ* = 168.59, 158.77, 133.82, 133.05, 132.57, 131.47, 127.47, 123.81, 121.29, 113.24, 67.44, 43.83, 31.31. **HRMS (ESI)**: *m/z* = 449.9912 calcd. for $[C_{16}H_{15}BrF_2NO_5P + H]^+$; found: 449.9911.

#### Synthesis of (2-(2-((2,2-difluoro-2-(2-fluorophenyl)ethyl)amino)-2-oxoethoxy)phenyl)phosphonic acid (**42**)

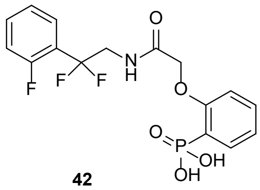

42

The phenylphosphonic acid derivative **42** was synthesized via GP **G** and GP **H**. GP **G** employed phosphonate **S-2** (202 mg, 1.00 mmol), potassium carbonate (276 mg, 2.00 mmol) and **S-39** (296 mg, 1.00 mmol) in DMF (6 mL). The crude intermediate was deprotected via GP **H** using TMSBr (1.32 mL, 10.00 mmol) in dry DCM (8 mL) and MeOH/$H_2O$ (3:1, 6 mL). The crude product was purified by HPLC to obtain **42** (158 mg, 0.41 mmol, 41 % over two steps) as a white solid. **LC-MS (ESI)**: $t_R$ = 6.11 min; *m/z* = 389.84 $[M + H]^+$, calcd. for $C_{16}H_{15}F_3NO_5P$: 389.06. **$^1$H NMR** (400 MHz, DMSO): *δ* = 9.46 (t, *J* = 6.3 Hz, 1H), 7.63 (ddd, *J* = 14.2, 7.7, 1.8 Hz, 1H), 7.60 – 7.44 (m, 2H), 7.38 (t, *J* = 7.7 Hz, 1H), 7.30 (dd, *J* = 11.4, 8.3 Hz, 1H), 7.22 (t, *J* = 7.6 Hz, 1H), 7.08 (td, *J* = 7.7, 7.3, 2.6 Hz, 2H), 4.61 (s, 2H), 3.96 (td, *J* = 14.8, 6.3 Hz, 2H). **$^{13}$C NMR** (101 MHz, DMSO): *δ* = 168.62, 158.73, 132.51, 127.37, 124.40, 121.14, 116.36, 113.16, 67.49, 43.10. **HRMS (ESI)**: *m/z* = 390.0713 calcd. for $[C_{16}H_{15}F_3NO_5P + H]^+$; found: 390.0712.

#### Synthesis of (2-(2-((2,2-difluoro-2-(3-fluorophenyl)ethyl)amino)-2-oxoethoxy)phenyl)phosphonic acid (**43**)

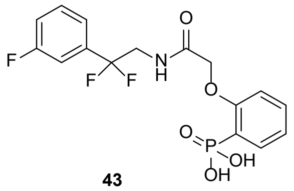

43

The phenylphosphonic acid derivative **43** was synthesized via GP **G** and GP **H**. GP **G** employed phosphonate **S-2** (202 mg, 1.00 mmol), potassium carbonate (276 mg, 2.00 mmol) and **S-39** (296 mg, 1.00 mmol) in DMF (6 mL). The crude intermediate was deprotected via GP **H** using TMSBr (1.32 mL, 10.00 mmol) in dry DCM (8 mL) and MeOH/$H_2O$ (3:1, 6 mL). The crude product was purified by HPLC to obtain **43** (111 mg, 0,28 mmol, 28 % over two steps) as a white solid. **LC-MS (ESI)**: $t_R$ = 6.21 min; *m/z* = 389.91 $[M + H]^+$, calcd. for $C_{16}H_{15}F_3NO_5P$: 389.06. **$^1$H NMR** (400 MHz, $CDCl_3$): *δ* = 9.42 (t, *J* = 6.3 Hz, 1H), 7.62 (ddd, *J* = 14.2, 7.4, 1.8 Hz, 1H), 7.53 – 7.42 (m, 2H), 7.37 – 7.24 (m, 3H), 7.08 (td, *J* = 8.1, 5.3 Hz, 2H), 4.64 (s, 2H), 3.90 (td, *J* = 14.5, 6.2 Hz, 2H).

**$^{13}$C NMR** (101 MHz, $CDCl_3$): *δ* = 168.64, 163.04, 160.60, 158.76, 136.97, 132.61, 132.55, 130.81, 121.18, 117.22, 113.16, 67.50, 43.48. **HRMS (ESI)**: *m/z* = 390.0713 calcd. for $[C_{16}H_{15}F_3NO_5P + H]^+$; found: 390.0714.

#### Synthesis of (2-(2-((2,2-difluoro-2-(4-fluorophenyl)ethyl)amino)-2-oxoethoxy)phenyl)phosphonic acid (**44**)

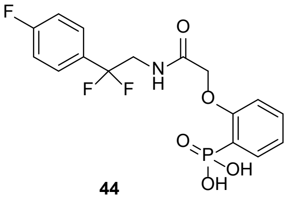

**44**

The phenylphosphonic acid derivative **44** was synthesized via GP **G** and GP **H**. GP **G** employed phosphonate **S-2** (117 mg, 0.58 mmol), potassium carbonate (160 mg, 1.04 mmol) and **S-41** (171 mg, 0.58 mmol) in DMF (6 mL). The crude intermediate was deprotected via GP **H** using TMSBr (0.77 mL, 5.80 mmol) in dry DCM (5 mL) and MeOH/$H_2O$ (3:1, 4 mL). The crude product was purified by HPLC to obtain **44** (92 mg, 0.24 mmol, 41 % over two steps) as a white solid. **LC-MS (ESI)**: $t_R$ = 6.25 min; *m/z* = 389.84 $[M + H]^+$, calcd. for $C_{16}H_{15}F_3NO_5P$: 389.06. **$^1$H NMR** (400 MHz, DMSO): *δ* = 9.41 (t, *J* = 6.3 Hz, 1H), 7.72 – 7.57 (m, 1H), 7.52 – 7.45 (m, 3H), 7.22 (t, *J* = 8.8 Hz, 2H), 7.14 – 7.02 (m, 2H), 4.63 (s, 2H), 3.86 (dt, *J* = 14.5, 7.2 Hz, 2H). **$^{13}$C NMR** (101 MHz, DMSO): *δ* = 168.58, 158.75, 133.07, 132.59, 127.76, 121.16, 115.57, 115.35, 113.24, 67.45, 43.97. **HRMS (ESI)**: *m/z* = 390.0713 calcd. for $[C_{16}H_{15}F_3NO_5P + H]^+$; found: 390.0711.

#### Synthesis of (2-(2-((2,2-difluoro-2-(3-(trifluoromethyl)phenyl)ethyl)amino)-2-oxoethoxy)phenyl)phosphonic acid (**45**)

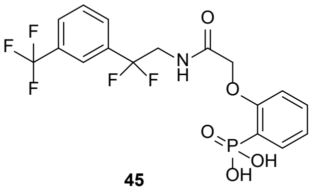

**45**

The phenylphosphonic acid derivative **45** was synthesized via GP **G** and GP **H**. GP **G** employed phosphonate **S-2** (202 mg, 1.00 mmol), potassium carbonate (276 mg, 2.00 mmol) and **S-42** (346 mg, 1.00 mmol) in DMF (6 mL). The crude intermediate was deprotected via GP **H** using TMSBr (1.32 mL, 10.00 mmol) in dry DCM (8 mL) and MeOH/$H_2O$ (3:1, 8 mL). The crude product was purified by HPLC to obtain **45** (194 mg, 0.44 mmol, 44 % over two steps) as a white solid. **LC-MS (ESI)**: $t_R$ = 7.49 min; *m/z* = 439.92 $[M + H]^+$, calcd. for $C_{17}H_{15}F_5NO_5P$: 439.06. **$^1$H NMR** (400 MHz, DMSO): *δ* = 9.50 (t, *J* = 6.3 Hz, 1H), 7.85 (d, *J* = 7.8 Hz, 1H), 7.81 – 7.71 (m, 2H), 7.71 – 7.58 (m, 2H), 7.48 (td, *J* = 7.9, 1.7 Hz, 1H), 7.06 (dt, *J* = 10.4, 6.9 Hz, 2H), 4.60 (s, 2H), 3.96 (td, *J* = 14.5, 6.3 Hz, 2H). **$^{13}$C NMR** (101 MHz, DMSO): *δ* = 168.67, 158.81, 135.74, 133.10, 132.55, 129.91, 129.62, 129.16, 127.18, 125.15, 123.18, 121.95, 121.32, 121.18, 113.24, 67.53, 43.83. **HRMS (ESI)**: *m/z* = 440.0681 calcd. for $[C_{17}H_{15}F_5NO_5P + H]^+$; found: 440.0687.

#### **Synthesis of intermediates for the derivatives 46 – 47**

#### Synthesis of ethyl 2,2-difluoro-2-(2-hydroxyphenyl)acetate (**S-43**)

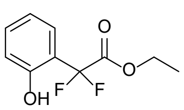

**S-43**

The phenylacetate derivative **S-43** was synthesized via GP **O** using 2-iodophenol (22.0 g, 100.0 mmol), ethyl bromodifluoroacetate (19.5 mL, 150.0 mmol) and activated copper (25.5 g, 390.0 mmol) in dry DMSO (260 mL). The crude product was purified by column chromatography (cyclohexane/ethyl acetate 4:1) to obtain **S-43** (6.73 g, 31.1 mmol, 31 %) as a yellow solid. **TLC** (cyclohexane:ethyl acetate, 4:1 v/v): $\mathbf{R_f}$ = 0.29. **$^1$H NMR** (400 MHz, $CDCl_3$): *δ* = 7.54 (dd, *J* = 7.9, 1.7 Hz, 1H), 7.38 – 7.22 (m, 1H), 7.02 – 6.89 (m, 2H), 4.34 (p, *J* = 7.1, 6.5 Hz, 2H), 1.46 – 1.22 (m, 3H). **$^{13}$C NMR** (101 MHz, $CDCl_3$): *δ* = 165.27, 154.13, 132.64, 126.25, 120.25, 117.09, 112.84, 63.59, 13.68.

#### Synthesis of ethyl 2,2-difluoro-2-(4-hydroxyphenyl)acetate (**S-44**)

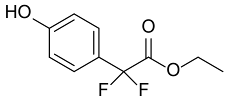

**S-44**

The phenylacetate derivative **S-44** was synthesized via GP **O** using 4-iodophenol (2.20 g, 10.00 mmol), ethyl bromodifluoroacetate (1.95 mL, 15.00 mmol) and activated copper (2.55 g, 39.00 mmol) in dry DMSO (26 mL). The crude product was purified by column chromatography (cyclohexane/ethyl acetate 4:1) to obtain **S-44** (565 mg, 2.62 mmol, 26 %) as a slightly yellow oil. **TLC** (cyclohexane:ethyl acetate, 4:1 v/v): $\mathbf{R_f}$ = 0.43. **$^1$H NMR** (400 MHz, $CDCl_3$): *δ* = 7.46 (d, *J* = 8.6 Hz, 2H), 6.88 (d, *J* = 8.6 Hz, 2H), 4.29 (q, *J* = 7.1 Hz,

2H), 1.30 (t, *J* = 7.2 Hz, 3H). **$^{13}$C NMR** (101 MHz, $CDCl_3$): *δ* = 164.99, 158.26, 127.33, 124.86, 115.66, 113.68, 63.39, 13.95.

#### Synthesis of 2-(2-(benzyloxy)phenyl)-2,2-difluoroethanol (**S-45**)

The alcohol **S-45** was synthesized via GP **G** and GP **P**. GP **G** employed acetate **S-43** (1.62 g, 7.50 mmol), potassium carbonate (2.08 g, 15.00 mmol) and benzylic bromide (0.89 mL, 7.50 mmol) in DMF (23 mL). The crude intermediate was reduced via GP **P** using $NaBH_4$ (0.56 g, 15.00 mmol) in MeOH (25 mL). The crude product was purified by column chromatography (cyclohexane/ethyl acetate 4:1) to obtain **S-45** (1.02 g, 3.87 mmol, 52 % over two steps) as a colorless oil. **TLC** (cyclohexane:ethyl acetate, 4:1 v/v): $R_f$ = 0.40. **LC-MS (ESI)**: $t_R$ = 9.33 min; *m/z* = 265.34 [M + H]$^+$, calcd. for $C_{15}H_{14}F_2O_2$: 264.10. **$^1$H NMR** (400 MHz, MeOD): *δ* = 7.56 (dd, *J* = 7.7, 1.8 Hz, 1H), 7.53 – 7.21 (m, 6H), 7.14 (d, *J* = 8.3 Hz, 1H), 7.03 (t, *J* = 7.6 Hz, 1H), 5.14 (s, 2H), 4.11 (t, *J* = 14.3 Hz, 2H). **$^{13}$C NMR** (101 MHz, MeOD): *δ* = 157.54, 138.16, 132.78, 129.55, 129.31, 128.93, 128.68, 128.41, 127.95, 121.53, 114.26, 71.41, 64.62.

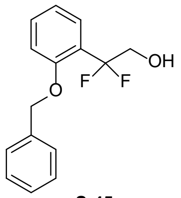


**S-45**

#### Synthesis of Ethyl 2-(4-(benzyloxy)phenyl)-2,2-difluoracetate (**S-46**)

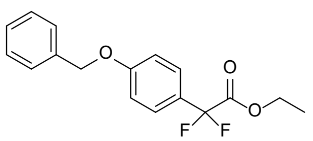


**S-46**

The acetate derivative **S-46** was synthesized via GP **G** using phenyl acetate **S-44** (4.00 g, 18.50 mmol), potassium carbonate (5.11 g, 37.00 mmol) and benzylic bromide (2.42 mL, 20.35 mmol) in DMF (55 mL). The crude product was purified by column chromatography (cyclohexane/ethyl acetate 19:1) to obtain **S-46** (1.70 g, 5.54 mmol, 30 %) as a colorless oil. **TLC** (cyclohexane:ethyl acetate, 19:1 v/v): $R_f$ = 0.43. **$^1$H NMR** (400 MHz, $CDCl_3$): *δ* = 7.54 (d, *J* = 8.7 Hz, 2H), 7.47 – 7.31 (m, 5H), 7.04 (d, *J* = 8.6 Hz, 2H), 5.10 (s, 2H), 4.29 (q, *J* = 7.2 Hz, 2H), 1.31 (dt, *J* = 27.1, 7.2 Hz, 3H). **$^{13}$C NMR** (101 MHz, $CDCl_3$): *δ* = 164.53, 160.89, 136.48, 128.79, 128.31, 127.58, 127.23, 125.29, 114.99, 70.23, 63.13, 14.01.

#### Synthesis of 2-(4-(benzyloxy)phenyl)-2,2-difluoroethanol (**S-47**)

The alcohol **S-47** was synthesized via GP **P** using the acetate **S-46** (1.70 g, 5.54 mmol), and $NaBH_4$ (0.21 g, 5.54 mmol) in MeOH (49 mL). The crude product was purified by column chromatography (cyclohexane/ethyl acetate 4:1) to obtain **S-47** (0.81 g, 3.07 mmol, 55 %) as a white solid. **TLC** (cyclohexane:ethyl acetate, 4:1 v/v): $R_f$ = 0.34. **LC-MS (ESI)**: $t_R$ = 9.41 min; *m/z* = 266.04 [M + H]$^+$, 244.99 [M-$H_2O$],$^+$, calcd. for $C_{15}H_{14}F_2O_2$: 264.10. **$^1$H NMR** (400 MHz, MeOD): *δ* = 7.48 – 7.40 (m, 4H), 7.38 – 7.30 (m, 3H), 7.05 (d, *J* = 8.6 Hz, 2H), 5.10 (s, 2H), 3.89 (t, *J* = 13.6 Hz, 2H). **$^{13}$C NMR** (101 MHz, MeOD): *δ* = 161.38, 138.33, 129.52, 128.94, 128.54, 128.15, 122.29, 115.69, 70.98, 65.94.

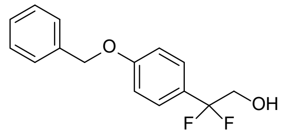


**S-47**

#### Synthesis of N-(2-(2-(benzyloxy)phenyl)-2,2-difluoroethyl)-2-bromoacetamide (**S-48**)

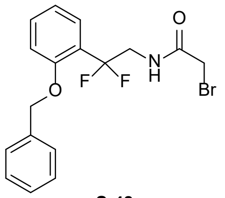


**S-48**

The bromoacetamide **S-48** was synthesized via GP **Q** and GP **D**. GP **Q** employed the alcohol **S-45** (983 mg, 3.72 mmol), pyridine (0.48 mL, 5.95 mmol) and $Tf_2O$ (0.69 mL, 4.09 mmol) in dry acetonirile (7.0 mL) as well as $NH_4OH$ (28%, 7.0 mL). The crude intermediate was coupled to the amide using triethylamine (0.57 mL, 4.09 mmol) and bromoacetyl bromide (0.36 mL, 4.09 mmol) in dry DCM (19 mL). The crude product was purified by column chromatography (cyclohexane/ethyl acetate 2:1) to obtain **S-48** (512 mg, 1.33 mmol, 36 % over two steps) as a white solid. **TLC** (cyclohexane:ethyl acetate, 2:1 v/v): $R_f$ = 0.43. **LC-MS (ESI)**: $t_R$ = 9.79 min; *m/z* = 383.69 [M + H]$^+$, calcd. for $C_{17}H_{16}BrF_2NO_2$: 383.03. **$^1$H NMR** (400 MHz, $CDCl_3$): *δ* = 7.53 (dd, *J* = 7.9, 1.8 Hz, 1H), 7.51 – 7.30 (m, 6H), 7.08 – 6.96 (m, 2H), 6.71 (t, *J* = 6.3 Hz, 1H), 5.17 (s, 2H), 4.17 (td, *J* = 14.4, 6.1 Hz, 2H), 3.79 (s, 2H). **$^{13}$C NMR** (101 MHz, $CDCl_3$): *δ* = 165.69, 156.19, 136.43, 132.24, 128.82, 128.22, 127.38, 120.86, 113.28, 70.77, 44.80, 28.91.

#### Synthesis of 2-(4-(benzyloxy)phenyl)-2,2-difluoroethanol (**S-49**)

The bromoacetamide **S-49** was synthesized via GP **Q** and GP **D**. GP **Q** employed the alcohol **S-46** (751 mg, 2.84 mmol), pyridine (0.37 mL, 4.54 mmol) and $Tf_2O$ (0.53 mL, 3.12 mmol) in dry acetonirile (5.0 mL) as well as $NH_4OH$ (28%, 5.0 mL). The crude intermediate was coupled to the amide using triethylamine (0.43 mL, 3.12 mmol) and bromoacetyl bromide (0.27 mL, 3.12 mmol) in dry DCM (14 mL). The crude product was purified by column chromatography (cyclohexane/ethyl acetate 2:1) to obtain **S-49** (450 mg, 1.17 mmol, 41 % over two steps) as a white solid.

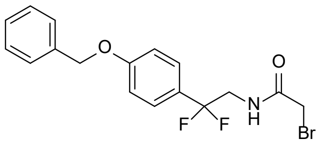

**S-49**

**TLC** (cyclohexane:ethyl acetate, 2:1 v/v): $R_f$ = 0.37. **LC-MS (ESI)**: $t_R$ = 9.76 min; *m/z* = 383.92 [M + H]$^+$, calcd. for $C_{17}H_{16}BrF_2NO_2$: 383.03. **$^1$H NMR** (400 MHz, $CDCl_3$): $\delta$ = 7.46 – 7.33 (m, 7H), 7.03 (d, *J* = 8.6 Hz, 2H), 6.80 (t, *J* = 6.2 Hz, 1H), 5.09 (s, 2H), 3.93 (td, *J* = 14.4, 6.2 Hz, 2H), 3.87 (s, 2H). **$^{13}$C NMR** (101 MHz, $CDCl_3$): $\delta$ = 165.90, 160.28, 136.57, 128.77, 128.27, 127.58, 126.87, 120.28, 114.99, 70.19, 46.15, 28.86.

### Synthesis of the derivatives 46 – 47

#### Synthesis of 2-((2-(2-(benzyloxy)phenyl)-2,2-difluoroethylcarbamoyl)methoxy)phenylphosphonic acid (**46**)

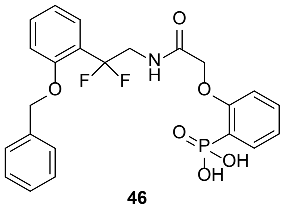

**46**

The phenylphosphonic acid derivative **46** was synthesized via GP **G** and GP **H**. GP **G** employed phosphonate **S-2** (187 mg, 0.93 mmol), potassium carbonate (256 mg, 1.86 mmol) and **S-48** (356 mg, 0.93 mmol) in DMF (6 mL). The crude intermediate was deprotected via GP **H** using TMSBr (1.22 mL, 9.30 mmol) in dry DCM (8 mL) and $MeOH/H_2O$ (3:1, 6 mL). The crude product was purified by HPLC to obtain **46** (109 mg, 0.23 mmol, 25 % over two steps) as a white solid. **LC-MS (ESI)**: $t_R$ = 7.61 min; *m/z* = 477.96 [M + H]$^+$, calcd. for $C_{23}H_{22}F_2NO_6P$: 477.12. **$^1$H NMR** (400 MHz, DMSO): $\delta$ = 9.20 (t, *J* = 6.3 Hz, 1H), 7.70 – 7.60 (m, 1H), 7.60 – 7.53 (m, 2H), 7.52 – 7.37 (m, 4H), 7.34 – 7.30 (m, 2H), 7.19 (d, *J* = 8.3 Hz, 1H), 7.05 (ddt, *J* = 14.0, 8.3, 4.4 Hz, 2H), 6.94 (t, *J* = 7.5 Hz, 1H), 5.21 (s, 2H), 4.59 (s, 2H), 4.10 (td, *J* = 14.7, 6.2 Hz, 2H). **$^{13}$C NMR** (101 MHz, DMSO): $\delta$ = 168.53, 158.78, 155.90, 136.83, 133.10, 132.66, 131.97, 128.56, 127.40, 123.24, 122.39, 121.25, 120.30, 113.48, 113.17, 69.83, 67.47, 42.64. **HRMS (ESI)**: *m/z* = 478.1226 calcd. for [$C_{23}H_{22}F_2NO_6P$ + H]$^+$; found: 478.1223.

#### Synthesis of 2-((2-(4-(benzyloxy)phenyl)-2,2-difluoroethylcarbamoyl)methoxy)phenylphosphonic acid (**47**)

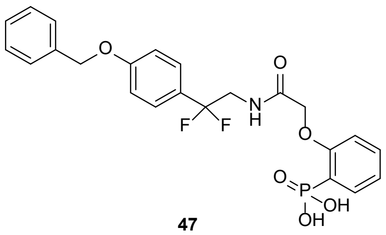

**47**

The phenylphosphonic acid derivative **47** was synthesized via GP **G** and GP **H**. GP **G** employed phosphonate **S-2** (202 mg, 1.00 mmol), potassium carbonate (276 mg, 2.00 mmol) and **S-49** (384 mg, 1.00 mmol) in DMF (6 mL). The crude intermediate was deprotected via GP **H** using TMSBr (1.32 mL, 10.00 mmol) in dry DCM (8 mL) and $MeOH/H_2O$ (3:1, 6 mL). The crude product was purified by HPLC to obtain **47** (110 mg, 0.23 mmol, 23 % over two steps) as a white solid. **LC-MS (ESI)**: $t_R$ = 7.73 min; *m/z* = 477.89 [M + H]$^+$, calcd. for $C_{23}H_{22}F_2NO_6P$: 477.12. **$^1$H NMR** (400 MHz, DMSO): $\delta$ = 9.38 (t, *J* = 6.3 Hz, 1H), 7.72 – 7.60 (m, 1H), 7.58 – 7.29 (m, 8H), 7.16 – 7.00 (m, 4H), 5.13 (s, 2H), 4.65 (s, 2H), 3.86 (td, *J* = 14.7, 6.2 Hz, 1H). **$^{13}$C NMR** (101 MHz, DMSO): $\delta$ = 168.57, 159.59, 158.75, 136.74, 133.04, 128.50, 127.76, 126.73, 121.12, 114.63, 69.35, 67.49, 44.08. **HRMS (ESI)**: *m/z* = 478.1226 calcd. for [$C_{23}H_{22}F_2NO_6P$ + H]$^+$; found: 478.1223.

### Synthesis of intermediates for the derivative 48

#### Synthesis of ethyl 2-(2-(dimethoxyphosphoryl)phenoxy)-2-fluoroacetate (**S-50**)

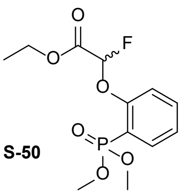

S-50

The fluoroacetate derivative **S-50** was synthesized via GP **G** using phosphonate **S-2** (1.01 g, 5.00 mmol), potassium carbonate (1.73 g, 12.50 mmol) and ethyl bromofluoroacetate (0.89 mL, 7.50 mmol) in DMF (15 mL). The crude product was purified by column chromatography (cyclohexane/ethyl acetate 2:1) to obtain **S-50** (1.41 g, 4.61 mmol, 92 %) as a colorless oil. **TLC** (cyclohexane:ethyl acetate, 2:1 v/v): $\mathbf{R_f}$ = 0.3. **LC-MS (ESI)**: $t_R$ = 7.39 min; *m/z* = 306.89 [M + H]$^+$, calcd. for $C_{12}H_{16}FO_6P$: 306.07. $^1$**H NMR** (400 MHz, MeOD): *δ* = 7.60 (ddt, *J* = 14.6, 7.6, 2.1 Hz, 1H), 7.46 (t, *J* = 8.3 Hz, 1H), 7.14 (t, *J* = 7.4 Hz, 1H), 7.11 – 7.05 (m, 1H), 5.99 (d, *J* = 58.6 Hz, 1H), 4.11 (q, *J* = 7.1 Hz, 2H), 3.56 (ddd, *J* = 11.4, 3.5, 1.7 Hz, 6H), 1.11 (t, *J* = 7.1 Hz, 3H). $^{13}$**C NMR** (101 MHz, MeOD): *δ* = 166.25, 158.68, 136.45, 135.94, 125.16, 117.32, 102.28, 63.61, 53.93, 14.29.

#### Synthesis of 2-(2-(Dimethoxyphosphoryl)phenoxy)-2-fluoroacetic acid (**S-51**)

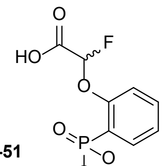

S-51

The fluoroacetate **S-50** (1.49 g, 4.86 mmol) was dissolved in MeOH (12 mL). An aqueous $K_2CO_3$ solution (1 M, 12 mL) was added and the mixture was stirred for 2 h at room temperature. The reaction mixture was acidified with 1M HCl solution and extracted with ethyl acetate. The combined organic phases were washed with brine, dried over $MgSO_4$ and the solvent was removed under reduced pressure to obtain **S-51** (0.90 g, 3.24 mmol, 67 %) as a colorless oil. **LC-MS (ESI)**: $t_R$ = 5.26 min; *m/z* = 278.85 [M + H]$^+$, calcd. for $C_{10}H_{12}FO_6P$: 278.04. $^1$**H NMR** (400 MHz, MeOD): *δ* = 7.64 (ddd, *J* = 14.6, 7.6, 1.8 Hz, 1H), 7.48 (t, *J* = 7.9 Hz, 1H), 7.17 (t, *J* = 7.5 Hz, 1H), 7.13 – 7.08 (m, 1H), 6.02 (d, *J* = 58.6 Hz, 1H), 3.58 (dt, *J* = 11.4, 2.3 Hz, 6H). $^{13}$**C NMR** (101 MHz, MeOD): *δ* = 166.28, 158.71, 136.47, 135.96, 125.17, 117.33, 104.57, 53.94.

### Synthesis of the derivative 48

#### Synthesis of 2-((2,2-difluoro-2-phenylethylcarbamoyl)fluoromethoxy)phenylphosphonic acid (**48**)

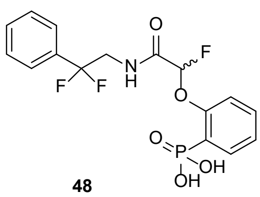

48

The phenylphosphonic acid derivative **48** was synthesized via GP **F** and GP **H**. GP **F** employed phosphonate **S-51** (278 mg, 1.00 mmol), *N*-Methylmorpholine (111 µL, 1.00 mmol) and isobutyl chloroformate (130 µL, 1.00 mmol) in dry THF (10 mL) as well as *N*-Methylmorpholine (111 µL, 1.00 mmol) and 2,2-difluoro-2-phenylethanamine (157 mg, 1.00 mmol). The crude intermediate was deprotected via GP **H** using TMSBr (1.32 mL, 10.00 mmol) in dry DCM (8 mL) and MeOH/$H_2O$ (3:1, 6 mL). The crude product was purified by HPLC to obtain **48** (70 mg, 18 mmol, 18 % over two steps) as a white solid. **LC-MS (ESI)**: $t_R$ = 6.25 min; *m/z* = 389.99 [M + H]$^+$, calcd. for $C_{16}H_{15}F_3NO_5P$: 389.06. $^1$**H NMR** (400 MHz, DMSO): *δ* = 10.01 (t, *J* = 6.3 Hz, 1H), 7.71 (ddd, *J* = 14.0, 7.5, 1.7 Hz, 1H), 7.66 – 7.38 (m, 6H), 7.26 (tt, *J* = 8.2, 4.3 Hz, 2H), 6.18 (d, *J* = 58.9 Hz, 1H), 4.07 – 3.71 (m, 2H). $^{13}$**C NMR** (101 MHz, DMSO): *δ* = 164.27, 156.79, 134.63, 132.82, 130.43, 128.64, 125.21, 123.98, 120.48, 116.66, 104.70, 102.45, 99.58, 91.34, 71.13, 44.59. **HRMS (ESI)**: *m/z* = 390.0713 calcd. for [$C_{16}H_{15}F_3NO_5P$ + H]$^+$; found: 390.0710.

### Synthesis of intermediates for the derivative 49

#### Synthesis of ethyl 2,2-difluoro-2-phenylacetate (**S-52**)

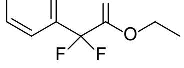

S-52

The phenylacetate derivative **S-52** was synthesized via GP **O** using iodobenzene (1.12 mL, 10.00 mmol), ethyl bromodifluoroacetate (1.95 mL, 15.00 mmol) and activated copper (2.55 g, 39.00 mmol) in dry DMSO (26 mL). The crude product was purified by column chromatography (cyclohexane/ethyl acetate 19:1) to obtain **S-52** (1.37 g, 6.83 mmol, 68 %) as a colorless oil. **TLC** (cyclohexane:ethyl acetate, 19:1 v/v): $\mathbf{R_f}$ = 0.43. **LC-MS (ESI)**: $t_R$ = 9.81 min; *m/z* = 200.74 [M + H]$^+$, calcd. for $C_{10}H_{10}F_2O_2$: 200.06. $^1$**H NMR** (400 MHz, $CDCl_3$): *δ* = 7.66 – 7.59 (m, 2H), 7.53 – 7.36 (m, 3H), 4.28 (q, *J* = 7.1 Hz, 2H), 1.29 (t, *J* = 7.2 Hz, 1H). $^{13}$**C NMR** (101 MHz, $CDCl_3$): *δ* = 164.26, 132.89, 131.07, 128.71, 125.53, 113.49, 63.19, 13.86.

#### Synthesis of 2,2-difluoro-2-phenylethanol (**S-53**)

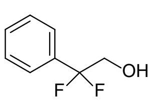

**S-53**

The alcohol **S-53** was synthesized via GP **P** using the acetate **S-52** (807 mg, 4.03 mmol), and $NaBH_4$ (152 mg, 4.03 mmol) in MeOH (14 mL). The crude product was purified by column chromatography (cyclohexane/ethyl acetate 2:1) to obtain **S-53** (493 mg, 3.12 mmol, 78 %) as a colorless oil. **TLC** (cyclohexane:ethyl acetate, 2:1 v/v): $\mathbf{R_f}$ = 0.57. **LC-MS (ESI)**: $t_R$ = 7.37 min; *m/z* = 159.62 $[M + H]^+$, calcd. for $C_8H_8F_2O$: 158.05. **$^1$H NMR** (400 MHz, $CDCl_3$): $\delta$ = 7.58 – 7.50 (m, 2H), 7.49 – 7.31 (m, 3H), 3.95 (t, *J* = 13.5 Hz, 1H), 2.41 – 2.29 (m, 1H). **$^{13}$C NMR** (101 MHz, $CDCl_3$): $\delta$ = 134.52, 130.41, 128.66, 125.59, 120.73, 66.06.

#### Synthesis of 2-bromo-N-(2,2-difluoro-2-phenylethyl)acetamide (**S-54**)

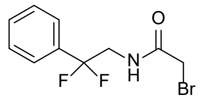

**S-54**

The bromoacetamide **S-54** was synthesized via GP **Q** and GP **D**. GP **Q** employed the alcohol **S-53** (1.08 g, 6.83 mmol), pyridine (0.88 mL, 10.93 mmol) and $Tf_2O$ (1.26 mL, 7.51 mmol) in dry acetonirile (12.4 mL) as well as $NH_4OH$ (28%, 12.4 mL). The crude intermediate was coupled to the amide using triethylamine (1.04 mL, 7.51 mmol) and bromoacetyl bromide (0.60 mL, 6.83 mmol) in dry DCM (34 mL). The crude product was purified by column chromatography (cyclohexane/ethyl acetate 2:1) to obtain **S-54** (0.88 g, 3.16 mmol, 46 % over two steps) as a yellow solid. **TLC** (cyclohexane:ethyl acetate, 2:1 v/v): $\mathbf{R_f}$ = 0.29. **LC-MS (ESI)**: $t_R$ = 8.11 min; *m/z* = 279.58 $[M + H]^+$, calcd. for $C_{10}H_{10}BrF_2NO$: 276.99. **$^1$H NMR** (400 MHz, $CDCl_3$): $\delta$ = 7.55 – 7.39 (m, 5H), 6.86 (s, 1H), 3.94 (td, *J* = 14.5, 6.2 Hz, 2H), 3.87 (s, 2H). **$^{13}$C NMR** (101 MHz, $CDCl_3$): $\delta$ = 165.85, 134.23, 130.66, 128.77, 125.32, 120.28, 46.09, 28.85.

### Synthesis of the derivative 49

#### Synthesis of 2-((2,2-difluoro-2-phenylethylcarbamoyl)methoxy)phenyl dihydrogen phosphate (**49**)

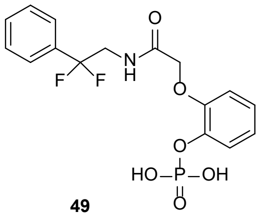

**49**

The dihydrogen phosphate derivative **49** was synthesized via GP **G** and GP **H**. GP **G** employed phosphate **S-17** (314 mg, 1.44 mmol), potassium carbonate (398 mg, 2.88 mmol) and **S-54** (398 mg, 1.44 mmol) in DMF (10 mL). The crude intermediate was deprotected via GP **H** using TMSBr (1.90 mL, 14.40 mmol) in dry DCM (8 mL) and MeOH/$H_2O$ (3:1, 6 mL). The crude product was purified by HPLC to obtain **49** (22 mg, 0.06 mmol, 4 % over two steps) as a white solid. **LC-MS (ESI)**: $t_R$ = 5.86 min; *m/z* = 387.90 $[M + H]^+$, calcd. for $C_{16}H_{16}F_2NO_6P$: 387.07. **$^1$H NMR** (400 MHz, MeOD): $\delta$ = 7.38 – 7.25 (m, 5H), 7.09 (s, 1H), 6.95 (t, *J* = 7.2 Hz, 1H), 6.85 – 6.75 (m, 2H), 4.41 (s, 2H), 3.77 (t, *J* = 13.9 Hz, 2H). **$^{13}$C NMR** (101 MHz, MeOD): $\delta$ = 163.33, 150.38, 136.26, 131.29, 129.54, 126.36, 123.20, 115.10, 101.34, 68.43. **HRMS (ESI)**: *m/z* = 388.0756 calcd. for $[C_{16}H_{16}F_2NO_6P + H]^+$; found: 388.0753.

### Synthesis of intermediates for the derivatives 50 – 53

#### Synthesis of ethyl 2-(2,6-difluorophenyl)-2,2-difluoroacetate (**S-55**)

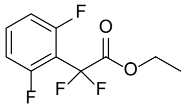

**S-55**

The phenylacetate derivative **S-55** was synthesized via GP **O** using 1,3-difluoro-2-iodobenzene (6.00 mL, 50.0 mmol), ethyl bromodifluoroacetate (9.62 mL, 75.0 mmol) and activated copper (12.75 g, 195.0 mmol) in dry DMSO (130 mL). The crude product was purified by column chromatography (cyclohexane/ethyl acetate 19:1) to obtain **S-55** (8.87 g, 37.55 mmol, 75 %) as a colorless oil. **TLC** (cyclohexane:ethyl acetate, 19:1 v/v): $\mathbf{R_f}$ = 0.25. **$^1$H NMR** (400 MHz, $CDCl_3$): $\delta$ = 7.48 – 7.41 (m, 1H), 6.97 (t, *J* = 8.9 Hz, 2H), 4.37 (q, *J* = 7.1 Hz, 2H), 1.33 (t, *J* = 7.2 Hz, 3H). **$^{13}$C NMR** (101 MHz, $CDCl_3$): $\delta$ = 162.85, 161.79, 159.33, 133.36, 112.77, 112.51, 111.27, 100.13, 63.70, 13.84.

#### Synthesis of ethyl 2-(2,5-difluorophenyl)-2,2-difluoroacetate (**S-56**)

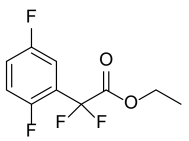
**S-56**

The phenylacetate derivative **S-56** was synthesized via GP **O** using 1,4-difluoro-3-iodobenzene (6.01 mL, 50.0 mmol), ethyl bromodifluoroacetate (9.62 mL, 75.0 mmol) and activated copper (12.75 g, 195.0 mmol) in dry DMSO (130 mL). The crude product was purified by column chromatography (cyclohexane/ethyl acetate 19:1) to obtain **S-56** (7.73 g, 37.7 mmol, 65 %) as a colorless oil. **TLC** (cyclohexane:ethyl acetate, 19:1 v/v): $\mathbf{R_f}$ = 0.38. **LC-MS (ESI)**: $t_R$ = 10.01 min; *m/z* = 236.66 [M + H]$^+$, calcd. for $C_{10}H_8F_4O_2$: 236.05. **$^1$H NMR** (400 MHz, $CDCl_3$): *δ* = 7.33 (ddd, *J* = 8.4, 5.6, 3.1 Hz, 1H), 7.22 – 7.04 (m, 2H), 4.34 (q, *J* = 7.2 Hz, 2H), 1.30 (t, *J* = 7.2 Hz, 3H). **$^{13}$C NMR** (101 MHz, $CDCl_3$): *δ* = 162.77, 159.68, 157.23, 119.85, 118.01, 114.40, 110.99, 108.49, 63.66, 13.82.

#### Synthesis of ethyl 2-(3,5-difluorophenyl)-2,2-difluoroacetate (**S-57**)

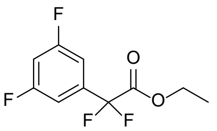
**S-57**

The phenylacetate derivative **S-57** was synthesized via GP **O** using 1,3-difluoro-5-iodobenzene (5.99 mL, 50.0 mmol), ethyl bromodifluoroacetate (9.62 mL, 75.0 mmol) and activated copper (12.75 g, 195.0 mmol) in dry DMSO (130 mL). The crude product was purified by column chromatography (cyclohexane/ethyl acetate 19:1) to obtain **S-57** (10.29 g, 43.6 mmol, 87 %) as a colorless oil. **TLC** (cyclohexane:ethyl acetate, 19:1 v/v): $\mathbf{R_f}$ = 0.50. **$^1$H NMR** (400 MHz, CDCl): *δ* = 7.13 (qd, *J* = 6.4, 5.2, 3.8 Hz, 2H), 6.93 (tt, *J* = 8.7, 2.4 Hz, 1H), 4.30 (q, *J* = 7.2 Hz, 2H), 1.31 (t, *J* = 7.2 Hz, 3H). **$^{13}$C NMR** (101 MHz, $CDCl_3$): *δ* = 164.28, 163.30, 161.79, 136.27, 114.69, 112.17, 109.19, 106.74, 63.71, 13.89.

#### Synthesis of ethyl 2-(3,4-difluorophenyl)-2,2-difluoroacetate (**S-58**)

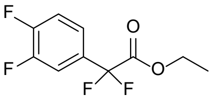
**S-58**

The phenylacetate derivative **S-58** was synthesized via GP **O** using 1,2-difluoroiodobenzene (6.03 mL, 50.0 mmol), ethyl bromodifluoroacetate (9.62 mL, 75.0 mmol) and activated copper (12.75 g, 195.0 mmol) in dry DMSO (130 mL). The crude product was purified by column chromatography (cyclohexane/ethyl acetate 19:1) to obtain **S-58** (9.42 g, 39.9 mmol, 80 %) as a colorless oil. **TLC** (cyclohexane:ethyl acetate, 19:1 v/v): $\mathbf{R_f}$ = 0.44. **$^1$H NMR** (400 MHz, $CDCl_3$): *δ* = 7.42 – 7.26 (m, 1H), 7.31 – 7.26 (m, 1H), 7.20 – 7.11 (m, 1H), 4.22 (q, *J* = 7.2 Hz, 2H), 1.23 (dd, *J* = 7.9, 6.6 Hz, 3H). **$^{13}$C NMR** (101 MHz, $CDCl_3$): *δ* = 163.46, 150.72, 148.93, 129.76, 122.25, 117.91, 115.35, 112.21, 63.43, 13.73.

#### Synthesis of 2-(2,6-difluorophenyl)-2,2-difluoroethanol (**S-59**)

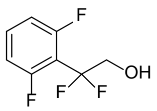
**S-59**

The alcohol **S-59** was synthesized via GP **P** using the acetate **S-55** (8.79 g, 37.21 mmol), and $NaBH_4$ (1.41 g, 37.21 mmol) in MeOH (130 mL). The crude product was purified by column chromatography (cyclohexane/ethyl acetate 2:1) to obtain **S-59** (4.32 g, 22.29 mmol, 60 %) as a colorless oil. **TLC** (cyclohexane:ethyl acetate, 2:1 v/v): $\mathbf{R_f}$ = 0.30. **$^1$H NMR** (400 MHz, MeOD): *δ* = 7.54 (tt, *J* = 8.4, 6.2 Hz, 1H), 7.13 – 7.00 (m, 2H), 4.05 (t, *J* = 13.8 Hz, 1H). **$^{13}$C NMR** (101 MHz, MeOD): *δ* = 163.10, 160.57, 133.90, 121.10, 113.68, 113.41, 112.42, 65.39.

#### Synthesis of 2-(2,5-difluorophenyl)-2,2-difluoroethanol (**S-60**)

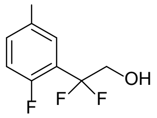
**S-60**

The alcohol **S-60** was synthesized via GP **P** using the acetate **S-56** (7.53 g, 31.90 mmol), and $NaBH_4$ (1.21 g, 31.90 mmol) in MeOH (110 mL). The crude product was purified by column chromatography (cyclohexane/ethyl acetate 2:1) to obtain **S-60** (4.84 g, 24.94 mmol, 78 %) as a white solid. **TLC** (cyclohexane:ethyl acetate, 2:1 v/v): $\mathbf{R_f}$ = 0.38. **$^1$H NMR** (400 MHz, DMSO): *δ* =δ 7.48 – 7.31 (m, 3H), 5.74 (t, *J* = 6.5 Hz, 1H), 3.91 (tdd, *J* = 14.2, 6.5, 1.2 Hz, 2H). **$^{13}$C NMR** (101 MHz, DMSO): *δ* = 158.96, 156.56, 123.43, 119.71, 119.16, 118.37, 115.02, 62.98.

#### Synthesis of 2-(3,5-difluorophenyl)-2,2-difluoroethanol (**S-61**)

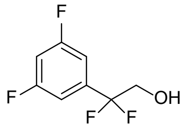

**S-61**

The alcohol **S-61** was synthesized via GP **P** using the acetate **S-57** (10.11 g, 42.80 mmol), and $NaBH_4$ (1.61 g, 42.80 mmol) in MeOH (150 mL). The crude product was purified by column chromatography (cyclohexane/ethyl acetate 2:1) to obtain **S-61** (7.57 g, 38.99 mmol, 91 %) as a colorless oil. **TLC** (cyclohexane:ethyl acetate, 2:1 v/v): $\mathbf{R_f}$ = 0.43. **$^1$H NMR** (400 MHz, $CDCl_3$): $\delta$ = 7.03 (dq, $J$ = 6.1, 2.3 Hz, 2H), 6.89 (tt, $J$ = 8.7, 2.3 Hz, 1H), 3.91 (ddd, $J$ = 14.6, 11.8, 1.4 Hz, 2H), 3.10 – 3.04 (m, 1H). **$^{13}$C NMR** (101 MHz, $CDCl_3$): $\delta$ 164.24, 161.75, 138.19, 119.49, 109.42, 109.15, 105.88, 65.47.

#### Synthesis of 2-(3,4-difluorophenyl)-2,2-difluoroethanol (**S-62**)

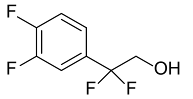

**S-62**

The alcohol **S-62** was synthesized via GP **P** using the acetate **S-58** (9.26 g, 39.24 mmol), and $NaBH_4$ (1.48 g, 39.24 mmol) in MeOH (135 mL). The crude product was purified by column chromatography (cyclohexane/ethyl acetate 2:1) to obtain **S-62** (6.01 g, 30.95 mmol, 79 %) as a colorless oil. **TLC** (cyclohexane:ethyl acetate, 2:1 v/v): $\mathbf{R_f}$ = 0.35. **$^1$H NMR** (400 MHz, MeOD): $\delta$ = 7.49 (ddd, $J$ = 11.6, 7.8, 2.1 Hz, 1H), 7.45 – 7.30 (m, 2H), 3.95 (t, $J$ = 13.2 Hz, 2H). **$^{13}$C NMR** (101 MHz, MeOD): $\delta$ = 152.55, 151.33, 133.78, 123.72, 121.15, 118.58, 116.59, 65.68.

#### Synthesis of Synthesis of 2-bromo-N-(2-(2,6-difluorophenyl)-2,2-difluoroethyl)acetamide (**S-63**)

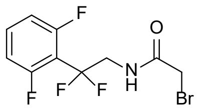

**S-63**

The bromoacetamide **S-63** was synthesized via GP **Q** and GP **D**. GP **Q** employed the alcohol **S-59** (4.19 g, 21.70 mmol), pyridine (2.80 mL, 34.72 mmol) and $Tf_2O$ (4.02 mL, 23.88 mmol) in dry acetonirile (39 mL) as well as $NH_4OH$ (28%, 39 mL). The crude intermediate was coupled to the amide using triethylamine (3.32 mL, 23.88 mmol) and bromoacetyl bromide (2.08 mL, 23.88 mmol) in dry DCM (80 mL). The crude product was purified by column chromatography (cyclohexane/ethyl acetate 2:1) to obtain **S-63** (1.65 g, 5.28 mmol, 24 % over two steps) as a white solid. **TLC** (cyclohexane:ethyl acetate, 2:1 v/v): $\mathbf{R_f}$ = 0.25. **LC-MS (ESI)**: $t_R$ = 8.36 min; $m/z$ = 313.73 $[M + H]^+$, calcd. for $C_{10}H_8BrF_4NO$: 312.97. **$^1$H NMR** (400 MHz, DMSO): $\delta$ = 8.86 (t, $J$ = 6.4 Hz, 1H), 7.60 (ddd, $J$ = 14.7, 8.4, 6.2 Hz, 1H), 7.18 (t, $J$ = 9.4 Hz, 2H), 3.94 (tt, $J$ = 17.7, 8.9 Hz, 2H), 3.82 (s, 2H). **$^{13}$C NMR** (101 MHz, DMSO): $\delta$ = 166.80, 160.87, 158.35, 133.56, 119.18, 113.03, 112.77, 110.19, 44.10, 28.56.

#### Synthesis of Synthesis of 2-bromo-N-(2-(2,5-difluorophenyl)-2,2-difluoroethyl)acetamide (**S-64**)

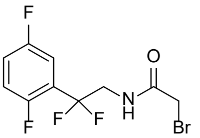

**S-64**

The bromoacetamide **S-64** was synthesized via GP **Q** and GP **D**. GP **Q** employed the alcohol **S-60** (4.65 g, 23.96 mmol), pyridine (3.10 mL, 38.34 mmol) and $Tf_2O$ (4.44 mL, 26.36 mmol) in dry acetonirile (43 mL) as well as $NH_4OH$ (28%, 43 mL). The crude intermediate was coupled to the amide using triethylamine (3.66 mL, 26.36 mmol) and bromoacetyl bromide (2.28 mL, 26.36 mmol) in dry DCM (112 mL). The crude product was purified by column chromatography (cyclohexane/ethyl acetate 1:1) to obtain **S-64** (3.89 g, 12.44 mmol, 52 % over two steps) as a white solid. **TLC** (cyclohexane:ethyl acetate, 1:1 v/v): $\mathbf{R_f}$ = 0.50. **LC-MS (ESI)**: $t_R$ = 8.58 min; $m/z$ = 313.79 $[M + H]^+$, calcd. for $C_{10}H_8BrF_4NO$: 312.97. **$^1$H NMR** (400 MHz, DMSO): $\delta$ = 8.78 (t, $J$ = 6.3 Hz, 1H), 7.49 – 7.34 (m, 2H), 7.32 (ddd, $J$ = 8.7, 5.8, 2.9 Hz, 1H), 3.94 (td, $J$ = 14.6, 6.3 Hz, 2H), 3.82 (s, 2H). **$^{13}$C NMR** (101 MHz, DMSO): $\delta$ = 166.77, 158.87, 156.47, 123.10, 119.44, 118.67, 118.42, 114.29, 43.41, 28.63.

#### Synthesis of Synthesis of 2-bromo-N-(2-(3,5-difluorophenyl)-2,2-difluoroethyl)acetamide (**S-65**)

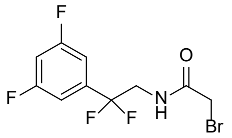

S-65

The bromoacetamide **S-65** was synthesized via GP **Q** and GP **D**. GP **Q** employed the alcohol **S-61** (7.46 g, 38.44 mmol), pyridine (4.96 mL, 61.52 mmol) and $Tf_2O$ (7.12 mL, 42.30 mmol) in dry acetonirile (68 mL) as well as $NH_4OH$ (28%, 68 mL). The crude intermediate was coupled to the amide using triethylamine (5.96 mL, 42.30 mmol) and bromoacetyl bromide (3.68 mL, 42.30 mmol) in dry DCM (120 mL). The crude product was purified by column chromatography (cyclohexane/ethyl acetate 2:1) to obtain **S-65** (4.40 g, 14.05 mmol, 37 % over two steps) as a white solid. **TLC** (cyclohexane:ethyl acetate, 2:1 v/v): $\mathbf{R_f}$ = 0.25. **LC-MS (ESI)**: $t_R$ = 8.71 min; *m/z* = 313.77 [M + H]$^+$, calcd. for $C_{10}H_8BrF_4NO$: 312.97. **$^1$H NMR** (400 MHz, DMSO): $\delta$ = 8.74 (t, *J* = 6.3 Hz, 1H), 7.38 (tt, *J* = 9.3, 2.4 Hz, 1H), 7.31 – 7.19 (m, 2H), 3.90 (td, *J* = 14.4, 6.2 Hz, 2H), 3.84 (s, 2H). **$^{13}$C NMR** (101 MHz, DMSO): $\delta$ = 166.78, 163.49, 161.15, 138.08, 119.61, 109.48, 109.20, 106.08, 44.06, 28.69.

#### Synthesis of 2-bromo-N-(2-(3,4-difluorophenyl)-2,2-difluoroethyl)acetamide (**S-66**)

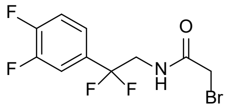

S-66

The bromoacetamide **S-66** was synthesized via GP **Q** and GP **D**. GP **Q** employed the alcohol **S-62** (5.79 g, 29.86 mmol), pyridine (3.86 mL, 47.78 mmol) and $Tf_2O$ (5.52 mL, 32.85 mmol) in dry acetonirile (54 mL) as well as $NH_4OH$ (28%, 54 mL). The crude intermediate was coupled to the amide using triethylamine (4.56 mL, 32.85 mmol) and bromoacetyl bromide (2.84 mL, 32.85 mmol) in dry DCM (108 mL). The crude product was purified by column chromatography (cyclohexane/ethyl acetate 2:1) to obtain **S-66** (3.76 g, 11.97 mmol, 40 % over two steps) as a yellow solid. **TLC** (cyclohexane:ethyl acetate, 2:1 v/v): $\mathbf{R_f}$ = 0.25. **LC-MS (ESI)**: $t_R$ = 8.47 min; *m/z* = 313.73 [M + H]$^+$, calcd. for $C_{10}H_8BrF_4NO$: 312.97. **$^1$H NMR** (400 MHz, $CDCl_3$): $\delta$ = 7.27 – 7.17 (m, 1H), 7.21 – 7.12 (m, 3H), 3.83 (td, *J* = 14.2, 6.3 Hz, 2H), 3.77 (s, 2H). **$^{13}$C NMR** (101 MHz, $CDCl_3$): $\delta$ = 166.51, 151.46, 148.84, 131.30, 122.01, 119.20, 117.90, 115.28, 45.29, 28.24.

### Synthesis of the derivatives 50 – 53

#### Synthesis of (2-(2-((2-(2,6-difluorophenyl)-2,2-difluoroethyl)amino)-2-oxoethoxy)phenyl)phosphonic acid (**50**)

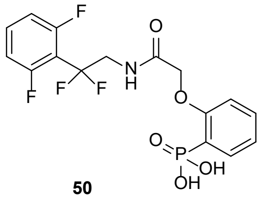

50

The phenylphosphonic acid derivative **50** was synthesized via GP **G** and GP **H**. GP **G** employed phosphonate **S-2** (202 mg, 1.00 mmol), potassium carbonate (276 mg, 2.00 mmol) and **S-63** (314 mg, 1.00 mmol) in DMF (6 mL). The crude intermediate was deprotected via GP **H** using TMSBr (1.32 mL, 10.00 mmol) in dry DCM (8 mL) and MeOH/$H_2O$ (3:1, 8 mL). The crude product was purified by HPLC to obtain **50** (60 mg, 0.15 mmol, 15 % over two steps) as a white solid. **LC-MS (ESI)**: $t_R$ = 6.68 min; *m/z* = 407.87 [M + H]$^+$, calcd. for $C_{16}H_{14}F_4NO_5P$: 407.05. **$^1$H NMR** (400 MHz, DMSO): $\delta$ = 9.57 (t, *J* = 6.4 Hz, 1H), 7.59 (dtd, *J* = 15.4, 7.9, 2.0 Hz, 2H), 7.48 (td, *J* = 7.9, 1.7 Hz, 1H), 7.18 – 7.01 (m, 4H), 4.63 (s, 2H), 3.94 (td, *J* = 14.8, 6.3 Hz, 2H). **$^{13}$C NMR** (101 MHz, DMSO): $\delta$ = 168.92, 158.84, 158.28, 133.49, 133.06, 132.64, 123.24, 121.44, 121.29, 121.16, 113.21, 113.13, 113.00, 112.76, 67.58, 43.63. **HRMS (ESI)**: *m/z* = 408.0618 calcd. for [$C_{16}H_{14}F_4NO_5P$ + H]$^+$; found: 408.0625.

#### Synthesis of (2-(2-((2-(2,5-difluorophenyl)-2,2-difluoroethyl)amino)-2-oxoethoxy)phenyl)phosphonic acid (**51**)

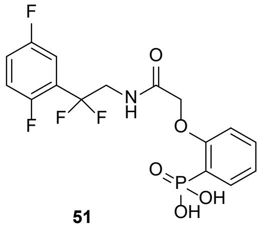

51

The phenylphosphonic acid derivative **51** was synthesized via GP **G** and GP **H**. GP **G** employed phosphonate **S-2** (202 mg, 1.00 mmol), potassium carbonate (276 mg, 2.00 mmol) and **S-64** (314 mg, 1.00 mmol) in DMF (6 mL). The crude intermediate was deprotected via GP **H** using TMSBr (1.32 mL, 10.00 mmol) in dry DCM (8 mL) and MeOH/$H_2O$ (3:1, 8 mL). The crude product was purified by HPLC to obtain **51** (146 mg, 0.36 mmol, 36 % over two steps) as a white solid. **LC-MS (ESI)**: $t_R$ = 6.89 min; *m/z* = 407.95 [M + H]$^+$, calcd. for $C_{16}H_{14}F_4NO_5P$: 407.05. **$^1$H NMR** (400 MHz, DMSO): $\delta$ = 9.47 (t, *J* = 6.4 Hz, 1H), 7.60 (ddd, *J* = 14.3, 7.5, 1.7 Hz, 1H), 7.47 (td, *J* = 7.9, 1.7 Hz, 1H), 7.41 – 7.35 (m, 2H), 7.16 (ddd, *J* = 8.6, 5.7, 3.0 Hz, 1H), 7.12 – 7.02 (m, 2H), 4.62 (s, 2H), 3.96 (td, *J* = 14.4, 6.3 Hz, 2H). **$^{13}$C NMR** (101 MHz,

DMSO): $\delta$ = 168.74, 158.74, 156.38, 133.09, 132.49, 123.10, 121.32, 121.19, 119.61, 119.28, 118.58, 118.25, 114.41, 113.22, 67.49, 42.88. **HRMS (ESI)**: *m/z* = 408.0618 calcd. for $[C_{16}H_{14}F_4NO_5P + H]^+$; found: 408.0586.

#### Synthesis of (2-(2-((2-(3,5-difluorophenyl)-2,2-difluoroethyl)amino)-2-oxoethoxy)phenyl)phosphonic acid (**52**)

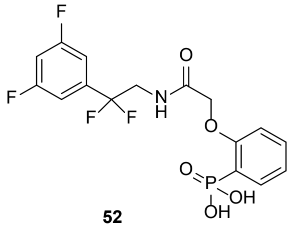


**52**

The phenylphosphonic acid derivative **52** was synthesized via GP **G** and GP **H**. GP **G** employed phosphonate **S-2** (202 mg, 1.00 mmol), potassium carbonate (276 mg, 2.00 mmol) and **S-65** (314 mg, 1.00 mmol) in DMF (6 mL). The crude intermediate was deprotected via GP **H** using TMSBr (1.32 mL, 10.00 mmol) in dry DCM (8 mL) and MeOH/$H_2O$ (3:1, 8 mL). The crude product was purified by HPLC to obtain **52** (133 mg, 0.33 mmol, 33 % over two steps) as a white solid. **LC-MS (ESI)**: $t_R$ = 7.33 min; *m/z* = 407.93 [M + H]$^+$, calcd. for $C_{16}H_{14}F_4NO_5P$: 407.05. **$^1$H NMR** (400 MHz, DMSO): $\delta$ = 9.43 (t, *J* = 6.3 Hz, 1H), 7.61 (ddd, *J* = 14.2, 7.4, 1.7 Hz, 1H), 7.48 (td, *J* = 7.9, 1.7 Hz, 1H), 7.38 (tt, *J* = 9.3, 2.4 Hz, 1H), 7.27 – 7.12 (m, 2H), 7.13 – 7.02 (m, 2H), 4.63 (s, 2H), 3.92 (td, *J* = 14.2, 6.3 Hz, 2H). **$^{13}$C NMR** (101 MHz, DMSO): $\delta$ = 168.71, 163.49, 160.91, 158.74, 133.05, 132.58, 123.24, 121.31, 119.50, 113.19, 109.44, 109.16, 106.04, 67.48, 43.59. **HRMS (ESI)**: *m/z* = 408.0618 calcd. for $[C_{16}H_{14}F_4NO_5P + H]^+$; found: 408.0593.

#### Synthesis of (2-(2-((2-(3,4-difluorophenyl)-2,2-difluoroethyl)amino)-2-oxoethoxy)phenyl)phosphonic acid (**53**)

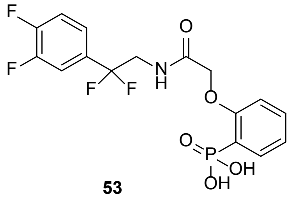


**53**

The phenylphosphonic acid derivative **53** was synthesized via GP **G** and GP **H**. GP **G** employed phosphonate **S-2** (202 mg, 1.00 mmol), potassium carbonate (276 mg, 2.00 mmol) and **S-66** (314 mg, 1.00 mmol) in DMF (6 mL). The crude intermediate was deprotected via GP **H** using TMSBr (1.32 mL, 10.00 mmol) in dry DCM (8 mL) and MeOH/$H_2O$ (3:1, 8 mL). The crude product was purified by HPLC to obtain **53** (261 mg, 0.64 mmol, 64 % over two steps) as a white solid. **LC-MS (ESI)**: $t_R$ = 6.43 min; *m/z* = 407.81 [M + H]$^+$, calcd. for $C_{16}H_{14}F_4NO_5P$: 407.05. **$^1$H NMR** (400 MHz, DMSO): $\delta$ = 9.40 (t, *J* = 6.4 Hz, 1H), 7.61 (ddd, *J* = 14.2, 7.5, 1.7 Hz, 1H), 7.60 – 7.36 (m, 3H), 7.28 (ddd, *J* = 8.8, 4.0, 1.9 Hz, 1H), 7.07 (td, *J* = 10.9, 9.6, 6.7 Hz, 2H), 4.63 (s, 2H), 3.9= (td, *J* = 14.1, 6.4 Hz, 2H). **$^{13}$C NMR** (101 MHz, DMSO): $\delta$ = 168.67, 158.75, 150.46, 148.01, 133.10, 132.54, 123.17, 122.77, 122.16, 121.19, 119.73, 117.76, 117.29, 113.15, 67.45, 43.76. **HRMS (ESI)**: *m/z* = 408.0618 calcd. for $[C_{16}H_{14}F_4NO_5P + H]^+$; found: 408.0617.

##### Synthesis of the derivatives 66

#### Synthesis of (N-(2-(3,4-difluorophenyl)-2,2-difluoroethyl)-2-(2-sulfamoylphenoxy)acetamide (**66**)

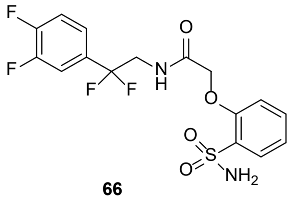


**66**

The sulfonamide derivative **66** was synthesized via GP **G** using sulfonamide **S-25** (173 mg, 1.00 mmol), potassium carbonate (276 mg, 2.00 mmol) and **S-66** (314 mg, 1.00 mmol) in DMF (6 mL). The crude product was purified by HPLC to obtain **66** (359 mg, 0.88 mmol, 88 %) as a white solid. **LC-MS (ESI)**: $t_R$ = 8.38 min; *m/z* = 406.75 [M + H]$^+$, calcd. for $C_{16}H_{14}F_4N_2O_4S$: 406.06. **$^1$H NMR** (400 MHz, DMSO): $\delta$ = 8.44 (t, *J* = 6.4 Hz, 1H), 7.77 (dd, *J* = 7.8, 1.7 Hz, 1H), 7.68 – 7.59 (m, 1H), 7.58 – 7.46 (m, 2H), 7.38 (ddd, *J* = 9.3, 4.0, 1.9 Hz, 1H), 7.28 (s, 2H), 7.18 – 7.06 (m, 2H), 4.76 (s, 2H), 3.99 (td, *J* = 14.3, 6.3 Hz, 2H). **$^{13}$C NMR** (101 MHz, DMSO): $\delta$ = 168.43, 154.40, 150.75, 148.30, 133.97, 131.92, 127.73, 122.98, 121.27, 119.91, 118.10, 115.71, 115.52, 114.09, 67.41, 43.88. **HRMS (ESI)**: *m/z* = 407.0683 calcd. for $[C_{16}H_{14}F_4N_2O_4S + H]^+$; found: 407.0687.
